# Topologically associating domains and their role in the evolution of genome structure and function in *Drosophila*

**DOI:** 10.1101/2020.05.13.094516

**Authors:** Yi Liao, Xinwen Zhang, Mahul Chakraborty, J.J. Emerson

**Affiliations:** Department of Ecology and Evolutionary Biology, University of California, Irvine, CA, USA; Center for Complex Biological Systems, University of California, Irvine, CA, USA

**Keywords:** *Drosophila*, TADs, Genome synteny, Structural variants, Gene regulation

## Abstract

Topologically associating domains (TADs) were recently identified as fundamental units of three-dimensional eukaryotic genomic organization, though our knowledge of the influence of TADs on genome evolution remains preliminary. To study the molecular evolution of TADs in *Drosophila* species, we constructed a new reference-grade genome assembly and accompanying high-resolution TAD map for *D. pseudoobscura*. Comparison of *D. pseudoobscura* and *D. melanogaster*, which are separated by ∼49 million years of divergence, showed that ∼30-40% of their genomes retain conserved TADs. Comparative genomic analysis of 17 *Drosophila* species revealed that chromosomal rearrangement breakpoints are enriched at TAD boundaries but depleted within TADs. Additionally, genes within conserved TADs exhibit lower expression divergence than those located in nonconserved TADs. Furthermore, we found that a substantial proportion of long genes (>50 kbp) in *D. melanogaster* (42%) and *D. pseudoobscura* (26%) constitute their own TADs, implying transcript structure may be one of the deterministic factors for TAD formation. Using structural variants (SVs) identified from 14 *D. melanogaster* strains, its 3 closest sibling species from the *D. simulans* species complex, and two obscura clade species, we uncovered evidence of selection acting on SVs at TAD boundaries, but with the nature of selection differing between SV types. Deletions are depleted at TAD boundaries in both divergent and polymorphic SVs, suggesting purifying selection, whereas divergent tandem duplications are enriched at TAD boundaries relative to polymorphism, suggesting they are adaptive. Our findings highlight how important TADs are in shaping the acquisition and retention of structural mutations that fundamentally alter genome organization.

## Introduction

Higher-order folding of eukaryotic genomes in the nucleus partitions the genome into multiple spatial layers, ranging from DNA loops to chromatin domains to compartments (Rowley and Corces 2018). A growing consensus has recognized chromatin domains, often referred to as topological associating domains (TADs), as fundamental units of three-dimensional (3D) genome organization (Szabo et al. 2018). Such domains are closely linked to important DNA-dependent and cellular processes, such as DNA replication (Pope et al. 2014), transcription (Schoenfelder and Fraser 2019), DNA-damage repair (Schmitt et al. 2016), development and cell differentiation (Zheng and Xie 2019). The presence of TADs or TAD-like domains has been widely characterized in species as varied as yeast (Mizuguchi et al. 2014), bacteria (Le et al. 2013), plants (Liu et al. 2017), and animals (Fishman et al. 2019), suggesting they represent a conserved feature of genome organization (Szabo et al. 2019).

TADs are thought to serve as regulatory units for controlling gene expression by promoting and constraining long-range enhancer-promoter interactions (Schoenfelder and Fraser 2019). Genes localizing within the same TAD tend to be co-regulated and co-expressed (Nora et al. 2012). Additionally, changes in TADs or their boundaries can alter expression of genes, including developmental and disease-related genes (Lupiáñez et al. 2015; Bonev et al. 2017; Akdemir et al. 2020). However, recent work re-examined the tight relationship between gene regulation and TADs by observing that disruption of TAD features can alter expression for only a small number of genes (Ghavi-Helm et al. 2019; Despang et al. 2019). Evolutionary comparisons across species (Krefting et al. 2018; Eres et al. 2019) permit us to quantify the nature of functional constraint in these domains, potentially reconciling the disparate observations above.

The conservation of TADs between distantly related species is associated with the preservation of synteny across vast spans of evolutionary time (Dixon et al. 2012). Genome rearrangement breakpoints are more common at TAD boundaries than inside TADs, implying that a subset of TADs is constrained to remain intact through evolution (Krefting et al. 2018; Lazar et al. 2018). In addition, deletions are depleted at TAD boundaries (Sadowski et al. 2019; Akdemir et al. 2020; Huynh and Hormozdiari 2019), presumably because deleting boundaries might destroy the insulating effect separating neighboring TADs. Thus, TAD boundaries share hallmarks of other functional genomic regions, like strong evolutionary constraints (Fudenberg and Pollard 2019). However, other than the dearth of deletions at TAD boundaries, it remains unclear whether natural selection drives the acquisition and fate of other classes of mutations, like duplications and transposable element (TE) insertions, when they occur at TAD boundaries.

TADs have been extensively analyzed using Hi-C in embryos (Sexton et al. 2012) and cell lines in *D. melanogaster* (Li et al. 2015; Chathoth and Zabet 2019; Wang et al. 2018). Hi-C data from other *Drosophila* species are primarily targeted for genome scaffolding and are too sparse in coverage for high resolution spatial organization analysis (Bracewell et al. 2019). In this study, we introduced a new genome assembly and created a high resolution Hi-C contact map for *D. pseudoobscura*, which diverged from *D. melanogaster* ∼49 million years ago (Thomas and Hahn 2017). We assessed 3D genome conservation between *D. pseudoobscura* and *D. melanogaster* using TADs annotated with high coverage Hi-C data. We also investigated the association between TADs and gene expression by analyzing public RNA-seq dataset (Yang et al. 2018). Additionally, we characterized genome rearrangement breakpoints in 17 *Drosophila* species (Miller et al. 2018; Mahajan et al. 2018) and their distribution along TAD regions. Finally, we investigated patterns of structural variants (SVs) at TAD boundaries with SV genotypes derived from reference-quality assemblies for intraspecies (14 *D. melanogaster* strains) (Chakraborty et al. 2019) and interspecies (*D. melanogaster* versus three simulans clade species; *D. pseudoobscura* versus *D. miranda*) (Chakraborty et al. 2020; Mahajan et al. 2018) comparisons. This study provides accurate, high-resolution resources for *Drosophila* genome structure research, and our findings highlight the evolutionary significance of TADs in shaping genome rearrangements, SVs, and gene expression.

## Results

### A new reference genome assembly for *D. pseudoobscura*

We *de novo* assembled the genome of *D. pseudoobscura* females from the strain MV-25-SWS-2005 using deep (280×) coverage Pacific Biosciences long reads (Supplemental Table S1). The resulting assembly consists of 72 contigs and is extremely contiguous, accurate, and complete (assembly size, ∼163.3 Mb; N50 contig length, 30.7 Mb; base-pair accuracy QV score, 52; and BUSCO (Simão et al. 2015), 99.6%) (Supplemental Tables S2, S3). The vast majority of three telocentric autosomes (Chr2, Chr3, and Chr4) and the dot chromosome, together with the complete circular mitochondrial DNA genome (mtDNA) are each assembled into a single contig. The X chromosome is represented in three contigs, including both arms of the euchromatic region and a repeat-rich contig (∼9.7 Mb) showing enrichment of centromere-specific repeats at both ends. These three contigs were combined into a single scaffold based on Hi-C contact maps (Supplemental Fig. S1). Additionally, we recovered 64 small and repetitive contigs (totaling ∼6.6 Mb) that were not scaffolded (Supplementary Table S4). The final assembly spans the majority of every major chromosome, interrupted only by two sequence gaps near the pericentromeric region of the X chromosome (Supplemental Fig. S2). While the middles of the chromosome arms are assembled contiguously, there are likely gaps remaining in the repetitive sequences at the ends of chromosomes, particularly in the centromeric and pericentromeric heterochromatin.

The current assembly is the most contiguous genome assembly for *D. pseudoobscura* (Supplemental Table S5). Approximately 26.9% (43.9 Mb) of the genome is annotated as repeat sequences, including 13.95% (22.77 Mb) derived from retrotransposons (Supplemental Table S6). We annotated 13,413 gene models using RNA-seq and full length mRNA sequencing (Iso-Seq) data (Supplemental Tables S6, S7). The largest structural difference between our assembly and two other assemblies from different strains (Bracewell et al. 2019; English et al. 2012) is an X-linked pericentromeric inversion (∼9.7 Mb). We verified this inversion with Hi-C contact frequency data and our sequenced strain shares a configuration with a closely related species, *D. miranda* (Mahajan et al. 2018) (Supplemental Fig. S2). The distribution of genomic features is generally conserved between *D. pseudoobscura* and *D. miranda*, despite several large chromosomal rearrangements reshuffling their genomes (Supplemental Fig. S2) (Bartolomé and Charlesworth 2006). We identified 1.43 million SNPs (∼10.3 per kbp), 0.53 million Indels (<50 bp; ∼3.7 per kbp), and 8,227 SVs (≥50 bp) that affected ∼1.8 Mb genomic regions between our assembly and the *Dpse*4.0 assembly from the reference strain (English et al. 2012) (Supplemental Fig. S3).

### Identification of TADs in *D. pseudoobscura* adult full bodies

We generated 397 million Hi-C paired-end reads (2×150 bp) from cross-linked DNA extracted from full bodies of adult females. We used the method of Arima Genomics that employs multiple restriction enzyme cutting sites for chromatin digestion, leading to a theoretical mean restriction fragment resolution of ∼160 bp. After filtering, about half of the raw data are retained for construction of the Hi-C contact map (Supplemental Table S8). This subset of read pairs yields a contact map with a maximal estimated “map resolution” of ∼800 bp, as calculated following the approach of Rao et al. (Rao et al. 2014).

TAD annotation may vary moderately among different computational methods (Forcato et al. 2017). To examine the consistency of TAD calls across methods, we identified TADs at 5-kbp resolution using three tools: HiCExplorer (Ramírez et al. 2018), Armatus (Filippova et al. 2014), and Arrowhead (Durand et al. 2016)(Fig. 1A), which inferred 1,013, 3,352, and 795 TADs, respectively. After excluding TADs shorter than 30 kbp, Armatus retained 858 TADs (mean length 123 kbp), Arrowhead retained 795 (mean length 140 kbp), and HiCExplorer retained 996 (mean length 146 kbp) (Fig. 1B, C). These tools yield results with different properties. For example, Arrowhead allows nested TADs and permits them to be spaced discretely, resulting in adjacent TADs separated by gaps. On the other hand, most adjacent TADs among HiCExplorer calls share boundaries. Partly for this reason, the calls from alternative methods do not overlap completely. Despite this, 589 TADs are shared by at least two tools, covering 57% (92.5/163 Mb) of the genome (Fig. 1B).

**Figure 1.**
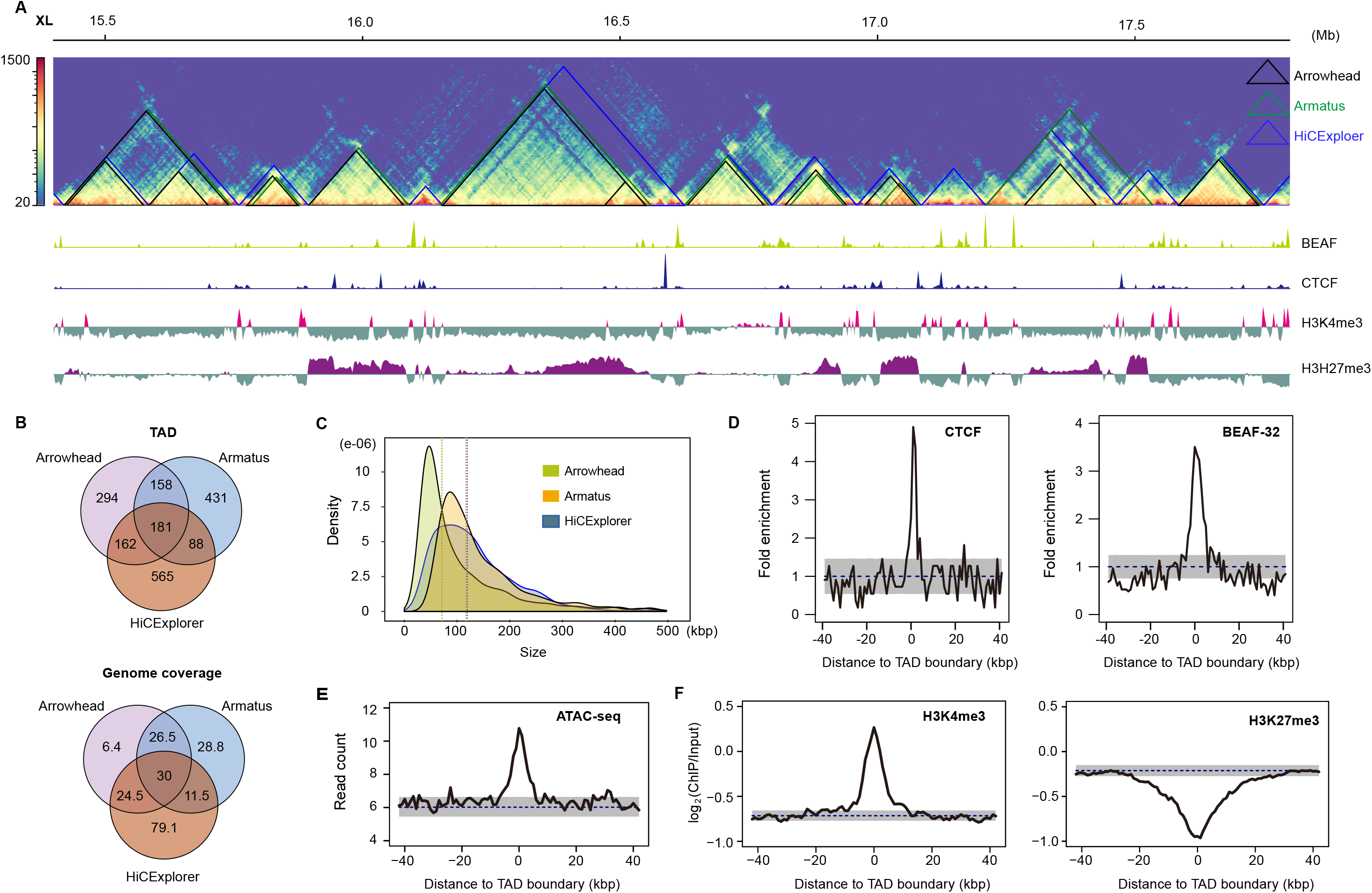
TADs annotated using full body samples in *D. pseudoobscura*. (*A*) Hi-C contact map (5-kbp resolution) on a ∼3 Mb region from the X chromosome with TADs annotated using Arrowhead (black), Armatus (green), and HiCExplorer (blue). The bottom browser tracks show the local profiles of binding sites of BEAF-32 and CTCF, as well as two histone marks (H3K4me3 and H3H27me3). (*B*) Overlap of TADs and their genome coverage annotated using three tools. (*C*) TAD size distribution for three tools. Vertical dashed lines represent the mean values. (*D*) Enrichment of CTCF and BEAF-32 binding sites at HiCExplorer TAD boundaries (*P* < 1.0 × 10^−4^). (*E*) Enrichment of ATAC-seq signal (open chromatin marks) at HiCExplorer TAD boundaries (*P* < 1.0 × 10^−4^). (*F*) TAD boundaries (HiCExplorer) are enriched in H3K4me3 (*P* < 1.0 × 10^−4^) but depleted for H3K27me3 marks (*P* < 1.0 × 10^−4^). *P*-values (*D, E, F*) were determined by permutation tests (n = 10,000); dashed lines represent mean values obtained from permutation tests; gray shaded areas, mean±SD in *D* and 95% intervals in *E* and *F*.

To determine whether TADs we inferred were supported by biological features, we investigated the chromatin landscape surrounding their boundaries using publicly available data (Schuettengruber et al. 2014; Yang et al. 2012; Ni et al. 2012). We found that TAD boundaries from all three tools are enriched for CTCF and BEAF-32 binding sites (Fig. 1D; Supplemental Fig. S4), as well as open and active chromatin as inferred from ATAC-seq data and H3K4me3 marks, respectively (Fig. 1E, F; Supplemental Fig. S4), but depleted for repressive chromatin marks, H3K27me3 (Fig. 1F; Supplemental Fig. S4). These observations are consistent with earlier studies in *D. melanogaster* (Wang et al. 2018; Chathoth and Zabet 2019; Hug et al. 2017), suggesting that Hi-C data from full bodies identifies a considerable amount of biologically meaningful TAD features in *Drosophila*. As HiCExplorer TAD calls are most strongly correlated with biological features (Fig. 1D-F; Supplemental Fig. S4), we used this set of TADs in most of the subsequent analysis, unless otherwise noted.

### Evolutionary conservation of TADs between *D. pseudoobscura* and *D. melanogaster*

We investigated the TAD conservation between two distantly related *Drosophila* species, *D. pseudoobscura* and *D. melanogaster*. We first identified blocks of synteny between their genomes. The resulting synteny map consists of 985 orthologous blocks larger than 10 kbp, with an average size of 101 kbp in *D. melanogaster* and 109 kbp in *D. pseudoobscura*, spanning 74.6% (100/134 Mb) of the *D. melanogaster* genome and 70% (110/157 Mb) of the *D. pseudoobscura* genome, respectively (Supplemental Table S9). The high-quality genome assemblies increased the average length of synteny blocks by 20% (100-83/83 kbp) compared to an earlier study (Richards et al. 2005). The synteny map (Fig. 2A) reveals extensive genome shuffling between species, but most of the orthologous blocks are located in the same Muller elements, suggesting that even on a small scale, translocations rarely occur between chromosomes in the course of *Drosophila* genome evolution, consistent with previous observations (Muller 1940; Schaeffer 2018).

**Figure 2.**
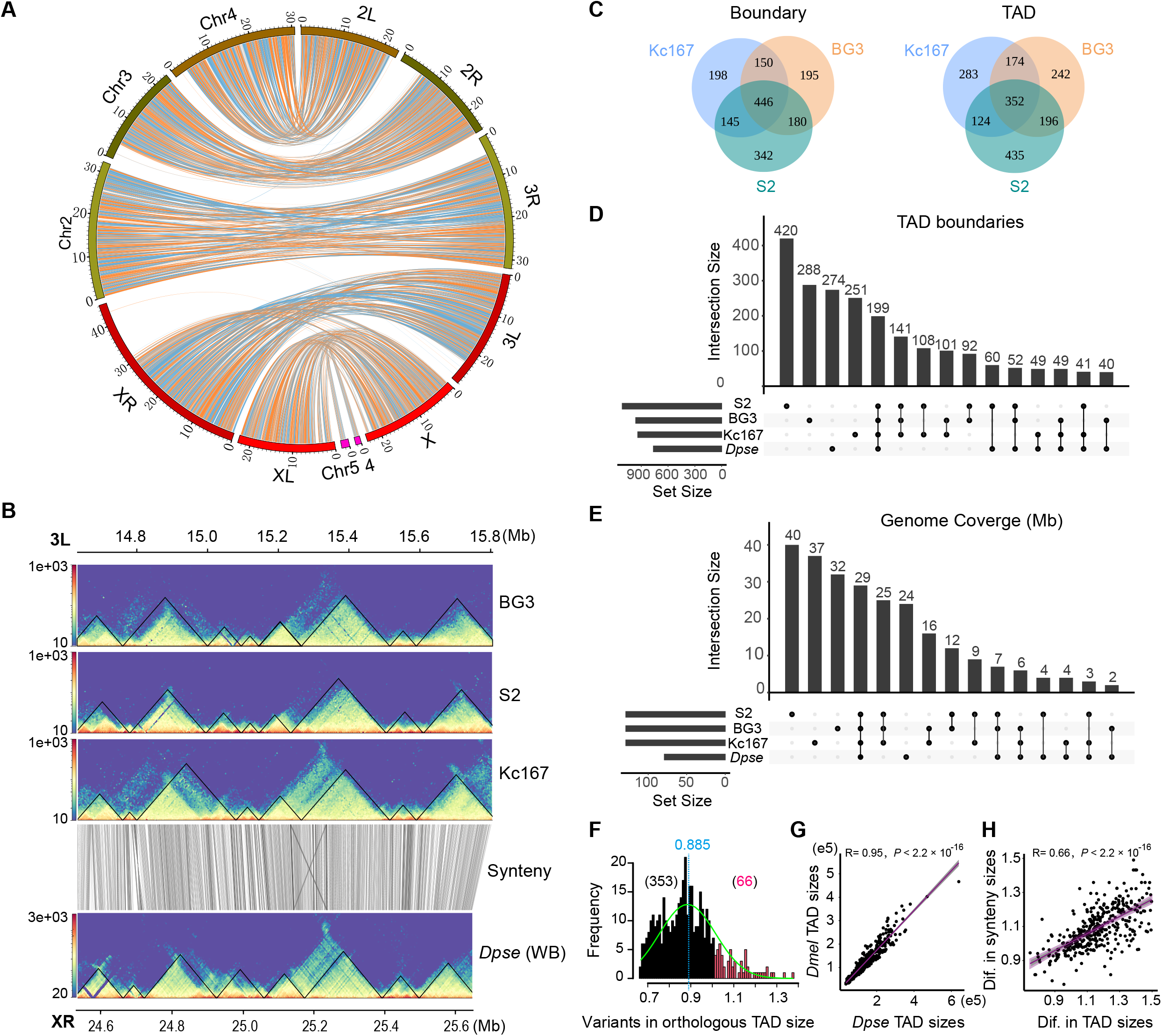
Evolutionary conservation of TADs between *D. pseudoobscura* (Dpse) and *D. melanogaster* (Dmel). (*A*) Genome synteny map between Dmel and Dpse constructed using 985 syntenic blocks larger than 10 kbp. (*B*) Conservation of TADs on a ∼1.2 Mb orthologous region between Dmel and Dpse. (*C*) Overlap of TAD features across three *D. melanogaster* cell lines. (*D*) Upset plot showing the overlap of TAD boundaries across three *D. melanogaster* cell lines and *D. pseudoobscura* whole body. 5-kbp boundaries for Dmel and 10-kbp boundaries for Dpse. (*E*) Upset plot showing genome coverage that maintained conserved TADs across three *D. melanogaster* cell lines and *D. pseudoobscura*. (*F*) Distribution of size variation (Dmel size divided by Dpse size) of orthologous TADs between Dmel and Dpse. (*G*) Correlation of the size of conserved TADs between *Dmel* and *Dpse*. (*H*) Correlation between the size difference of the orthologous TADs and the size difference of local synteny blocks where the orthologous TADs located between Dmel and Dpse. Dif., Difference.

For *D. melanogaster*, we annotated TADs using published Hi-C data from three cell lines (S2, Kc167, and BG3) (Chathoth and Zabet 2019; Wang et al. 2018) (Supplemental Table S10). TADs were annotated using HiCExplorer as described for our *D. pseudoobscura* data. Consistent with previous studies (Ulianov et al. 2016), TADs and their boundaries are largely shared across cell lines (Fig. 2B; Supplemental Figs. S5, S6). About 68% of TADs and 76% of their boundaries (10-kbp boundaries) are shared at least in two cell lines, whereas 32% and 24% of the TADs and boundaries are cell line-specific (Fig. 2C).

For each interspecific comparison, genomic coordinates of TAD features (i.e. body and boundary) were lifted over between species (Table 1). The success rate for each lift is 66.9-71.5% for bodies and 72.5-75.4% for boundaries, comparable to their shared genome synteny fraction (70-74.6%). Of TADs annotated across the three *D. melanogaster* cell lines, we found that 29.4-32% of body annotations and 31.5-35.3% of boundary annotations were conserved with *D. pseudoobscura* (Table 1). The conserved TADs cover at least 34.5% (46.2/134 Mb) of the *D. melanogaster* genome. In the reciprocal comparison, ∼41.4% of TADs, spanning ∼39.7% (62.4/157 Mb) of the *D. pseudoobscura* genome, and 48.5% of the boundaries were conserved with *D. melanogaster* (Table 1). Such rates of conservation are substantially higher than the background levels observed from permuted genomic coordinates based on two different null hypotheses (Table 1; Methods). We note that adjusting the stringency of the reciprocal overlap requirement from 80% to 90% did not qualitatively alter the results but decreased the conservation rate (e.g. genome coverage) from 39.7% to 27.8% (Supplemental Table S12). Thus, the conservative estimate is that nearly half of the syntenic regions of the genome (30-40% of the total genome) retained conserved TADs between *D. melanogaster* and *D. pseudoobscura* (Fig. 2D, E; Supplemental Fig. S7), and TADs in the remaining half may have diverged. This estimate is consistent with our visual inspection of Hi-C contact maps across the long syntenic regions between these two species (Fig. 2B and Supplemental Fig. S5). The inferred conservation magnitude is comparable to some regulatory phenotypes between these two species, such as functional enhancer conservation (46%) (Arnold et al. 2014) and CTCF binding sites (30%) (Ni et al. 2012), but lower than others, such as BEAF-32 (>70%) (Yang et al. 2012) and alternative splicing (∼80%) (Malko 2006).

**Table 1.** Summary of TAD annotation (HiCExplorer), lift and conservation between *D. melanogaster* (Release 6) and *D. pseudoobscura* (this study).

| TAD Features | Species (sample) | Total | Lifted | Conserved | Genome Cov. [C/L/T] (Mb) | P-values |
| --- | --- | --- | --- | --- | --- | --- |
| Body | *Dmel* (Kc167) | 933 | 640 (68.6%) | 291 (31.2%) | 43.6/88.2/131.4 | $1 \times 10^{-4}$ |
| | *Dmel* (BG3) | 956 | 668 (69.3%) | 308 (32.0%) | 46.2/90.0/131.3 | $1 \times 10^{-4}$ |
| | *Dmel* (S2) | 1,107 | 792 (71.5%) | 325 (29.4%) | 43.0/91.5/132.4 | $1 \times 10^{-4}$ |
| | *Dpse* (WB) | 1,013 | 678 (66.9%) | 419 (41.4%) | 62.4/90.5/145.6 | $1 \times 10^{-4}$ |
| Boundary<sup>a</sup> | *Dmel* (Kc167) | 939 | 683 (72.7%) | 331 (35.3%) | - | $2.2 \times 10^{-16}$ |
| | *Dmel* (BG3) | 962 | 722 (75.1%) | 339 (35.2%) | - | $2.2 \times 10^{-16}$ |
| | *Dmel* (S2) | 1,113 | 807 (72.5%) | 351 (31.5%) | - | $2.2 \times 10^{-16}$ |
| | *Dpse* (WB) | 1,019 | 768 (75.4%) | 494 (48.5%) | - | $2.2 \times 10^{-16}$ |

353/419 conserved TADs are larger in *D. pseudoobscura* than their orthologs in *D. melanogaster*, whereas the rest are larger in *D. melanogaster* (Fig. 2F). The sizes of the orthologous TADs are correlated with the sizes of the local syntenic blocks (Fig. 2G, H), suggesting that TADs may expand or contract proportionally with their local genomic regions. On average, *D. psedoobscura* TADs are ∼10% larger than their counterparts in *D. melanogaster*, smaller than the ∼17% difference (157-134/134 Mb) observed between their genome sizes, suggesting that size variation among homologous TADs is more strongly constrained than genome size variation elsewhere.

### Conservation of different TAD boundary classes

TAD boundaries may differ in insulation strengths, binding affinity of insulators, occurrence across cell types, and the flanking chromatin states. These properties may distinguish their evolutionary conservation. To test this, we first assessed the conservation of boundaries overlapping with binding sites for six insulator proteins in each of the three *D. melanogaster* cell lines (Kc167, BG3, and S2) (Supplementary Table S13). We found that TAD boundaries overlapping BEAF-32, CP190, and Chromator binding sites are more frequently shared between species than those lacking these sites. However, we observed no such pattern for CTCF, Su(Hw), and Trl (Fig. 3A; Supplemental Fig. S8). These findings are consistent with previous observations that BEAF-32, CP190, and Chromator are better predictors of TAD boundaries in *Drosophila* (Wang et al. 2018; Ramírez et al. 2018).

**Figure 3.**
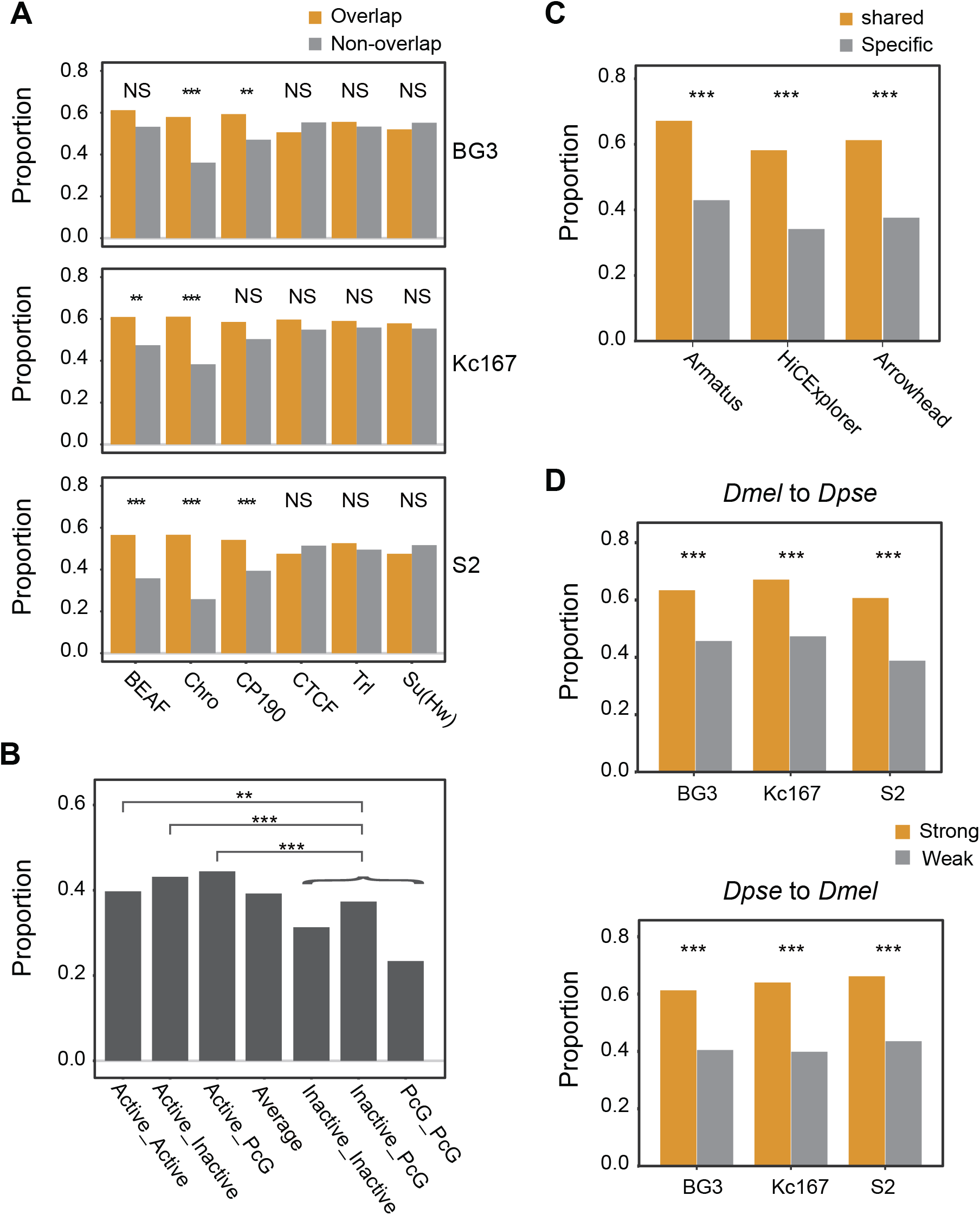
Conservation of distinct classes of TAD boundaries between *D. melanogaster* and *D. pseudoobscura*. (*A*) Boundaries that overlap with binding sites of architectural proteins versus those lacking the corresponding binding sites. (*B*) Boundaries linked with active TADs versus inactive TADs. (*C*) Boundaries that share across cell lines versus cell-line-specific boundaries. (*D*) Strong versus weak boundaries. Statistical significance were calculated using Chi-square test (\*\*\**P* < 0.001; **<0.01; NS: no significance) (for details, see Supplemental Table S14).

We also investigated the degree of conservation of the boundaries of TADs associated with different chromatin marks. Using the data from (Ramírez et al. 2018), we found that boundaries associated with active TADs are more conserved than those of TADs enriched for inactive and PcG marks or those lacking chromatin marks altogether (Fig. 3B), which is consistent with the fact that active chromatin may play crucial roles in TAD formation (Ulianov et al. 2016).

Next, we partitioned TAD boundaries into cell-specific boundaries and boundaries observed in more than one cell line. HiCExplorer identified 921 D. melanogaster TAD boundaries shared by at least two cell lines, whereas 735 were cell-specific (Supplemental Table S14). Of the 672/921 boundaries that we successfully lifted over to *D. pseudoobscura*, 58% (391/672) are conserved with *D. pseudoobscura*, which is significantly higher than cell-specific boundaries (*P* < 0.001, Chi-square test) in which only 34% (165/483) are conserved with *D. pseudoobscura* (Fig. 3C). Similar results were also obtained for Arrowhead and Armatus calls (Fig. 3C). However, this pattern can also be caused by the potential underrepresentation of cell-specific boundaries in the *D. pseudoobscura* whole body TAD set.

Finally, we compared stronger and weaker boundaries, wherein boundary strengths were categorized based on HiCExplorer TAD-separation score. Both types of boundaries tend to be observed in multiple cell lines, but weaker boundaries overlap more cell-specific boundaries (Supplemental Table S15). Hence, as indicated above, we expect stronger boundaries to be more conserved than weaker ones. We indeed observed this in the comparisons between *D. pseudoobscura* and each of the three *D. melanogaster* cell lines, and also in the reciprocal comparisons (Fig. 3D).

### The potential roles of TADs in *Drosophila* gene regulation

To investigate the potential link between TADs and gene regulation, we compared expression profiles across 7 tissues from both sexes for 10,921 one-to-one orthologs between *D. melanogaster* and *D. pseudoobscura* (Yang et al. 2018). The orthologs were classified by their locations with respect to TADs or TAD boundaries (Juicer call), covering the following categories: 1) genes inside versus outside TADs; 2) genes inside conserved versus nonconserved TADs; 3) genes inside cell-specific versus those shared by multiple cell types TADs; 4) genes within 20 kbp of conserved versus nonconserved TAD boundaries; and 5) genes within 20 kbp of cell-specific and those shared by multiple cell types TAD boundaries. To measure expression divergence, we calculated both Euclidean distance and Pearson’s correlation coefficient distance (Pereira et al. 2009). Both distances show that comparisons made using partitions from 1-3 above were statistically significant, whereas those made using 4-5 above (those involving proximity to TAD boundaries) were not (Fig. 4A; Supplemental Fig. S9A). These results show that TADs correlate with evolutionary stability of gene expression because genes inside TADs, conserved TADs, and TADs that are shared by multiple cell types, all show significantly lower expression divergence than their corresponding counterparts, respectively.

**Figure 4.**
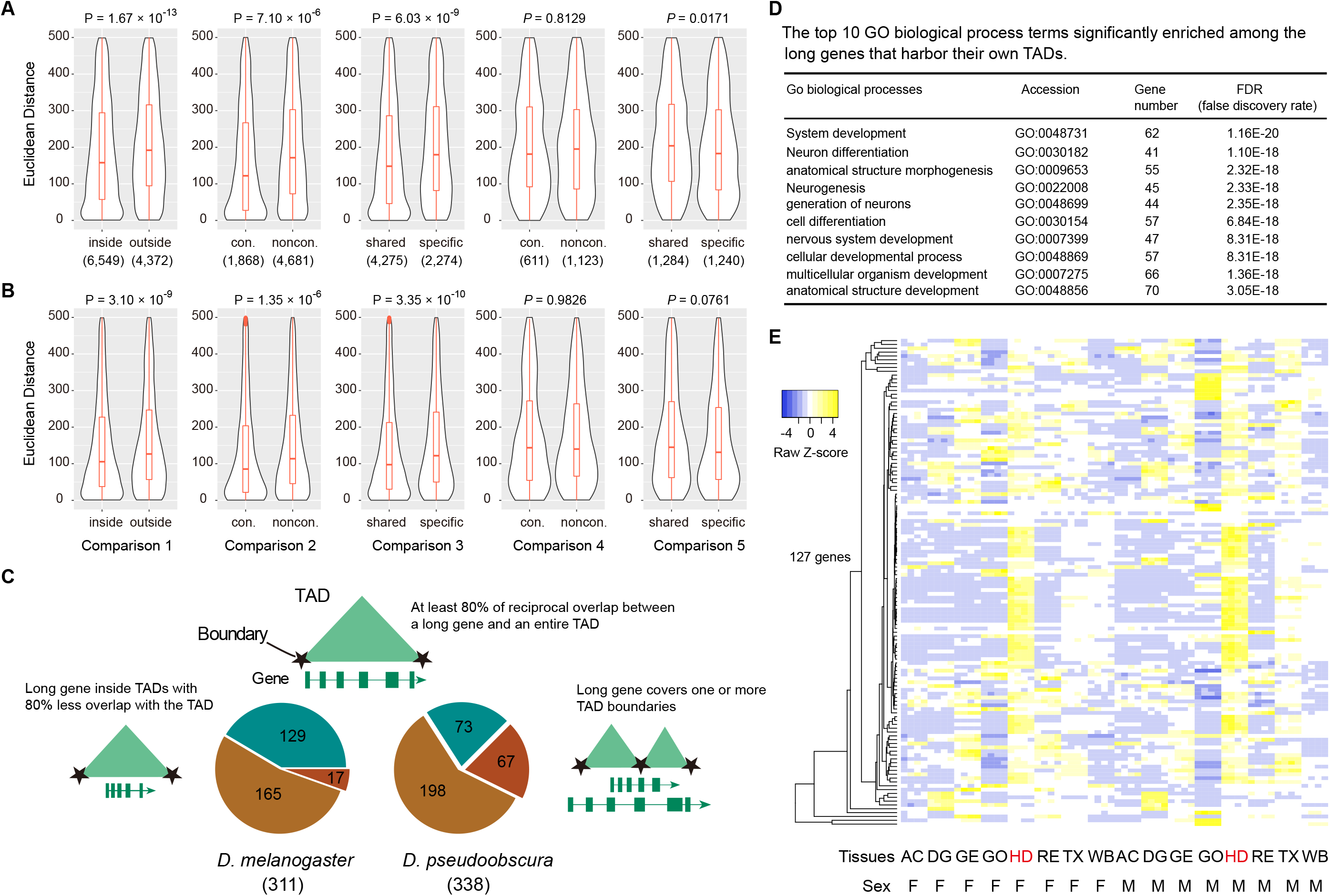
The potential roles of TADs in gene regulation in *Drosophila*. (*A*) Expression divergence measured by Euclidean distance for one-to-one orthologs between *D. melanogaster* and *D. pseudoobscura*. (*B*) Expression variation measured by Euclidean distance for the same gene sets used in the above interspecific comparison between two *D. melanogaster* strains, OreR and w1118. (*C*) Physical overlap of long genes and TADs in *D. melanogaster* and *D. pseudoobscura*. (*D*) Representative GO biological process terms significantly enriched among the 127 long genes that constitute their own TADs in *D. melanogaster*. (*E*) Expression profile of the 127 long genes across eight tissues in both male and female in *D. melanogaster*. AC, abdomen without digestive or reproductive system; DG, digestive plus excretory system; GE, genitalia; GO, gonad; HD, head; RE, reproductive system without gonad; TX, thorax without digestive system; WB, whole body. F, female; M, male.

We also considered the possibility that the stability of gene expression in annotated or conserved TAD regions is driven by gene content rather than TADs per se. To test this, we repeated the above analyses using the same gene set (10,921) between two *D. melanogaster* strains, OreR and w1118, with expression data across 8 tissues from both sexes (Yang et al. 2018). This intraspecies analysis (assuming intraspecific TAD differences can be largely neglected) revealed similar patterns as those of cross-species comparisons (Fig. 4B; Supplemental Fig. S9B), suggesting that genes constrained in the annotated or conserved TAD regions indeed tend to be more stable.

We further investigated the link between the gene structures and TADs by focusing on protein-coding genes larger than 50 kbp. Of 311 those long genes (> 50 kbp) in *D. melanogaster*, which span 30.35 Mb of the genome, we found that TAD boundaries are significantly depleted inside genes compared to the genome-wide background (143/1,113 S2 boundaries; *P* = 4.86 × 10^−15^, proportion test against 30.35/134 Mb, or 0.226). Only 17 long genes contained TAD boundaries inside them in all three cell types (Fig. 4C). 129 long genes (Supplementary Table S16), comprising 15 Mb of the *D. melanogaster* genome, individually occupied an intact TAD predicted in at least one of the three cell lines or tools (Supplementary Fig. S10). In *D. pseudoobscura*, we identified 338 long genes (>50 kbp), but we found relatively fewer long genes (73) that spanned full TADs (Supplemental Table S17). This may be due to the fact that *D. pseudoobscura* TADs were annotated only in the whole body, whereas TADs in *D. melanogaster* come from three cell lines. This observation raises the possibility that some TADs emerge only in certain situations (e.g. cell types or developmental stages) to regulate their corresponding genes. This prediction is supported by Gene Ontology analysis (Supplemental Methods), which shows that functions of the long genes are enriched for development processes, particularly those affecting the nervous system (Fig. 4D), consistent with the observation that they are preferentially expressed in head (Fig. 4E, Supplemental Fig. S11).

### Evolutionary genome rearrangement breakpoints coincide with *Drosophila* TAD boundaries

Conservation of a considerable fraction of TADs between two distantly related *Drosophila* species prompted us to investigate whether TADs are evolutionarily constrained. To this end, we identified 1,061 genome rearrangement breakpoints between *D. melanogaster* and *D. pseudoobscura* (Fig. 5A; Supplemental Fig. S12) and found that they are enriched at TAD boundaries (257/1,061; *P* < 2.2 × 10^−16^, proportion test against 11.13 Mb S2 HiCExplorer boundary regions as a proportion of total 134 Mb genome size, or 0.083), consistent with previous results (Fishman et al. 2019; Lazar et al. 2018).

**Figure 5.**
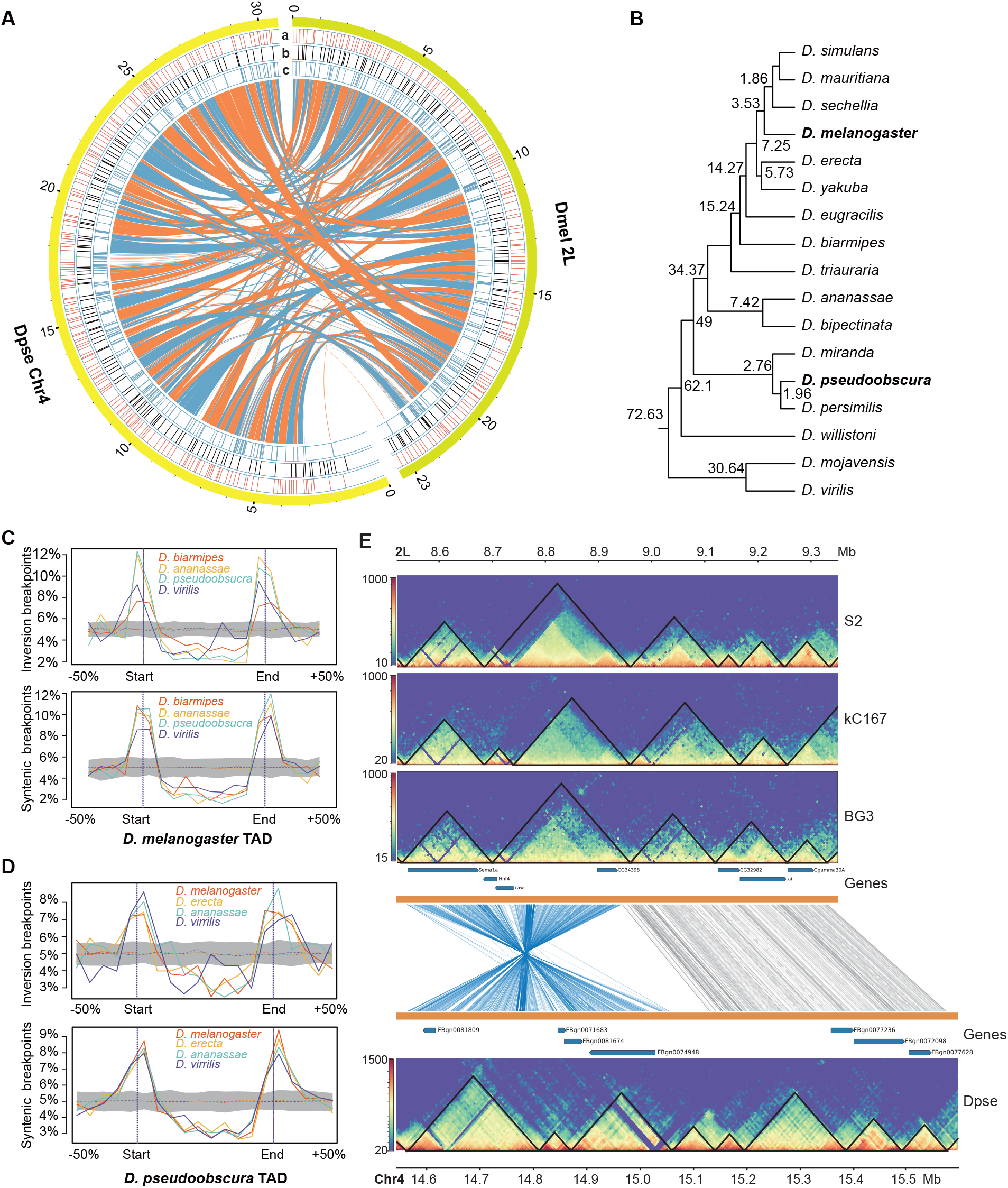
Evolutionary genome rearrangement breakpoints are enriched at *Drosophila* TAD boundaries. (*A*) Synteny map between *D. melanogaster* 2L and *D. pseudoobscura* Chr4. Tracks a: HiCExperor TAD boundaries annotated at restriction fragment resolution; b: 10 kbp resolution; c: synteny breakpoints. (*B*) Phylogeny of 17 *Drosophila* species. Estimated divergence times are obtained from (Thomas and Hahn 2017) except for *D. triauraria*. (*C*) Distribution of genome rearrangement breakpoints between *D. melanogaster* and four other *Drosophila* species along TAD regions. (*D*) Distribution of genome rearrangement breakpoints between *D. pseudoobscura* and four species along TAD regions. (*E*) Conservation of TADs in an inverted genomic segment (*D. melanogaster* 2L: 8.55 - 8.95 Mb) between *D. melanogaster* and *D. pseudoobscura*.

We extended the analysis to 17 *Drosophila* species, spanning 72 million years of evolution (Fig. 5B) (Thomas and Hahn 2017). All these species possess highly contiguous genome assemblies with contig N50 larger than 4Mb (Miller et al. 2018; Mahajan et al. 2018), permitting reliable identification of genome rearrangement breakpoints. Across comparisons between *D. melanogaster* and the other 16 species, we identified from 108 to 1,180 synteny and from 10 to 314 inversion breakpoints (Supplemental Table S18). For most comparisons, we observed these breakpoints were enriched at TAD boundaries, whereas the frequency of breakpoints was depleted within TADs (Fig. 5C; Supplemental Fig. S13A). However, in comparisons between *D. melanogaster* and its closest relatives (*D. sechellia, D. mauritiana*, and *D. simulans*), the small number of events likely did not offer enough power to observe this pattern.

We repeated this analysis using *D. pseudoobscura* as the reference, identifying from 259 to 1,242 synteny and from 60 to 359 inversion breakpoints in the 16 analogous comparisons (Supplemental Table S18). We observed the same pattern as above (Fig. 5D; Supplemental Fig. S13B). Such enrichment of rearrangement breakpoints at TAD boundaries suggests that a fraction of TADs are evolutionarily constrained even among genomes as extensively rearranged as those in the genus *Drosophila*. For example, three TADs are well preserved on a ∼450kb inverted genomic segment between *D. melanogaster* and *D. pseudoobscura* across 49 million years of evolution (Fig. 5E).

### TADs shape structural genomic variants at their boundaries in *Drosophila*

To investigate the potential role of TADs in shaping patterns of structural genomic variants, we obtained two comprehensive SV datasets (comprising deletions, TE insertions, non-TE insertions, and tandem duplications) both based on reference-quality genome assemblies (Fig. 6A), which include: (1) a polymorphic SV dataset (Fig. 6B) from 14 *D. melanogaster* strains (Chakraborty et al. 2019); and (2) a divergence SV dataset spanning ∼3 million years of evolution between *D. melanogaster* and the three members of the *D*. simulans species complex (Fig. 6C) (Chakraborty et al. 2020). Deletions and insertions were polarized using *D. erecta* and *D. yakuba* as outgroups. The resulting unfolded allele frequency spectrum (Fig. 6D) shows that, in the polymorphic dataset, most SVs are observed in only a single strain, with TE insertions exhibiting the greatest proportion (∼92%) of rare variants than other types. However, a considerable proportion (35-63%) of interspecific SVs are present in at least two species (Fig. 6E), suggesting that these SVs occurred prior to at least one species-splitting event in the *D. simulans* clade and therefore have existed for a relatively longer period of time than intraspecies SVs.

**Figure 6.**
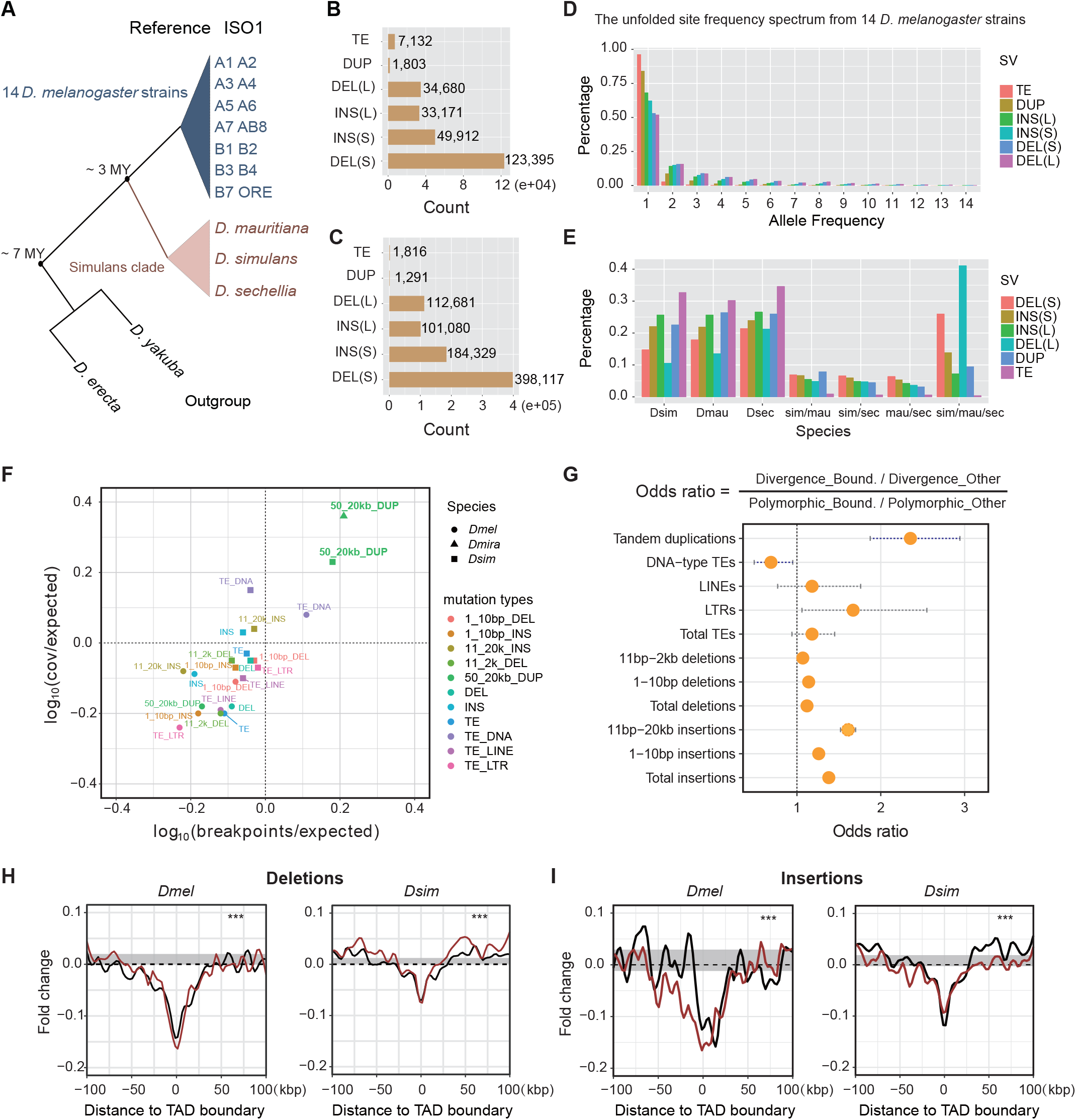
Patterns of structural variants at *Drosophila* TAD boundaries. (*A*) Highly contiguous genome assemblies from 14 *D. melanogaster* strains and three *D. simulans* clade species, together with two outgroup species, *D. erecta* and *D. yakuba*. (*B*) Nonredundant SVs, including TE insertions, tandem duplications (DUP), Non-TE insertions (INS, “S” represents insertions size range from 1-10 bp and “L” represents insertions size range from 11 bp to 20 kbp), and deletions (DEL, “S” for 1-10 bp and “L” for 11 bp to 2 kbp) identified from the 14 *D. melanogaster* strains. (*C*) Nonredundant SVs identified in the three *D. simulans* clade species. (*D*) The unfolded site frequency spectrum of SVs from 14 *D. melanogaster* strains. (*E*) Phylogenetic profiling of SVs among the three *D. simulans* clade species. (*F*) Tests of purifying selection on SVs at TAD boundaries using Fudenberg and Pollard’s method. (*G*) Odds ratios of 2×2 contingency tables with margins categorizing polymorphism/divergence and boundary/non-boundary mutations. Confidence intervals are calculated from the Fisher exact test results. (*H*) Deletions from both datasets are depleted at the TAD boundaries. (*I*) Non-TE insertions from both datasets are depleted at the TAD boundaries. Red and black lines represent larger and shorter variants, respectively. (\*\*\**P* < 1×10^−4^, permutation test) (Supplemental Fig. S15).

Using these two datasets, we sought to assess the mode of selection on SVs at *Drosophila* TAD boundaries by exploiting the method described previously (Fudenberg and Pollard 2019). We used a conservative set of 2,156 TAD boundaries in the euchromatin regions (Supplemental Table S19) that are shared between two independent studies (Ramírez et al. 2018; Wang et al. 2018). We find that both deletions and small non-TE insertions are depleted at TAD boundaries in both polymorphic and divergence SV datasets (Fig. 6F; Supplemental Fig. S14), consistent with that purifying selection may act to remove them to prevent disruption of TAD boundaries. A contrast was observed for tandem duplications (TDs). TDs from 14 *D. melanogaster* strains are depleted at TAD boundaries as compared to genome-wide background (181/3,606 breakpoints; *P* = 2.83 × 10^−8^, proportion test against 8.6 Mb boundary regions as a proportion of total 117 Mb euchromatin regions, or 0,074), whereas TDs from the three *D. simulans* clade species are significantly enriched at TAD boundaries (287/2,582 breakpoints; *P* = 3.63 × 10^−13^, proportion test against 0.074). The enrichment of TDs at TAD boundaries are also observed in the comparison between *D. pseudoobscura* and *D. miranda* (293/2,728 breakpoints, *P* < 2.2 × 10^−16^, permutation test against 8.86 Mb/133.62 Mb) (Fig. 6F; Supplemental Table S20). This excess divergence relative to polymorphism may indicate the action of adaptive selection (Andolfatto 2005). To quantify this excess for all other genomic variants studied, we calculated the odds ratios by dividing the ratio of genomic variants occurring at TAD boundary regions versus the rest of euchromatin regions in the divergence dataset by the same ratio computed from polymorphism (Fig. 6G). The highest excess value of TDs (287/2,295 in divergence TDs versus 181/3,425 in polymorphic TDs; *P* < 2.2 × 10^−16^, Fisher exact test) suggests that TDs at TAD boundaries might have evolved under adaptive natural selection.

Enrichment of TDs at TAD boundaries is consistent with previous observations that TAD boundaries tend to act as relatively frequent targets of duplications (Sadowski et al. 2019) and co-duplicate with super enhancers (Gong et al. 2018). To determine what kinds of sequences are associated with duplicated TAD boundaries, we inspected 181 boundary TDs in the three *D*. simulans clade species, finding that they rarely overlap large-scale genome rearrangement breakpoints, instead about 88% (160/181) of them overlap genes. These genes are weakly enriched for constitutive genes (i.e. genes that are expressed in most or all cells of an organism) (70/169; *P* = 0.018, proportion test against the expected proportion of 5,854/17,473) (Supplemental Methods), consistent with the previous observation that TAD boundaries are enriched in housekeeping genes (Hug et al. 2017).

Finally, with the large number of SVs we identified in our datasets, we also investigated SVs in the surrounding TAD boundary regions. Confirming the observations above, deletions and non-TE insertions are broadly depleted around TAD boundaries with peaks at the boundaries (Fig. 6H, I). Larger deletions in the polymorphic dataset are more depleted at TAD boundaries, implying larger deletions may be more deleterious to TAD boundaries (Fig. 6H). It is worth noting that TE insertions exhibit complex patterns at TAD boundaries. For example, long terminal repeat retrotransposons (LTRs) and long interspersed nuclear elements (LINEs) are strongly depleted at TAD boundaries (*P* < 1.0 × 10^−4^, permutation test) in the polymorphic dataset, but such pattern is not shown in the divergence dataset or for DNA-type TEs (Supplemental Fig S15).

## Discussion

Our knowledge of 3D genome evolution remains limited (Yang et al. 2019). To interrogate the evolutionary patterns of genome topology and its potential association with genome structure and function, we generated a reference-quality genome assembly and high resolution Hi-C data for *D. pseudoobscura*.

Although our *D. pseudoobscura* Hi-C data was obtained from whole body samples, we observed high consistency with several biological features known to be associated with TADs, regardless of the specific approaches employed in their annotation. For example, our TAD annotations are correlated with epigenetic states (e.g. H3K4me3 and H3K27me3), and their boundaries are enriched for insulator binding sites (e.g. CTCF and BEAF-32) and open and active chromatin marks (Fig. 1). These observations are consistent with the fact that TADs are a largely invariant feature across tissues in a given organism (Ulianov et al. 2016). While CTCF appears not to be a major TAD boundary definition protein in *Drosophila* (Szabo et al. 2019), we nevertheless observed enrichment of CTCF at our TAD boundaries, likely because it connects TAD borders in a cell-specific manner in *Drosophila* (Chathoth and Zabet 2019).

Our analysis revealed that TADs are conserved across at least 30-40 percent of the genomes between D. melanogaster and *D. pseudoobscura*. This rate is comparable to that observed between the much more closely-related comparison between humans and their closest sister species, chimpanzees (∼43%) (Eres et al. 2019). The conservation we observe is substantially higher than that among three distantly related *Drosophila* species-*D. melanogaster, D. busckii* and *D. virilis* (∼10%) (Renschler et al. 2019). Such incongruity is perhaps explained by differences in the quality of genome assemblies, depth of Hi-C data, or the evolutionary distance between species comparisons. Despite the difference, both results showed that a substantial proportion of TADs persist for long periods of time during evolution, suggesting they are functionally relevant. It is worth noting that our study likely still underestimates conservation. Our estimates were derived from pairwise comparisons between three distinct cell lines (Kc167, S2, and BG3) in *D. melanogaster* but whole body in *D. pseudoobscura*. Given extensive cell and allele-specific variability of TADs observed using single-cell Hi-C (Nagano et al. 2013) and super-resolution fluorescence in situ hybridization (FISH) imaging approaches (Bintu et al. 2018), TADs identified in whole body samples might represent topologies averaged across multiple tissues and millions of cells. Thus, many cell-, tissue-, or developmental stage-specific TADs may be underrepresented. Future experiments that carefully match samples (e.g. the same cell types or tissues) may provide a path to address this problem.

The role of TADs in gene regulation remains a matter of active research. Recent thinking suggests a reciprocal interplay between spatial genome organization and transcription, in which each is able to modulate or reinforce the activity of the other (van Steensel and Furlong 2019). Our results and others (Krefting et al. 2018) have revealed that the evolutionary stability of TADs correlates to constraint on gene expression, suggesting TADs may play roles in gene regulation. This effect is potentially confounded by the properties of the gene content in conserved TADs, though these two possibilities are by no means mutually exclusive. For example, the pattern of constraint we see in gene regulation is already established on intraspecific timescales in *D. melanogaster*, potentially before enough time has elapsed to establish variation between constraint in TADs.

Our finding that a large proportion of long genes coincides with entire TADs implies that transcription may be one of the deterministic factors for the establishment and maintenance of spatial genomic organization, or, conversely, that TADs are important in the regulation of long genes. Such gene-level chromatin domains are reminiscent of self-loop structures of genes found in *Arabidopsis thaliana* (Liu et al. 2016) and gene crumples in *S. cerevisiae* (Hsieh et al. 2015). Moreover, such gene-level domains are more likely to be cell-, tissue-, or developmental stage-specific since we detected substantially more (129 or 46%) genes in individual cell lines in *D. melanogaster* than in whole body samples (73 or 26%) in *D. pseudoobscura*.

Our analyses, in combination with previous works (Krefting et al. 2018; Lazar et al. 2018; Fishman et al. 2019; Renschler et al. 2019) show that genome rearrangement breakpoints acquired during evolution preferentially occur at TAD borders, suggesting that rearrangements resulting in disruption of TAD integrity are subjected to negative selection. Moreover, it suggests that TADs tend to evolve as intact structural units in genome shuffling, probably due to their putative functional constraint. Nevertheless, the non-random distribution of chromosomal breaks can also be explained by the “fragile regions” model that breaks of chromosome occur at a higher frequency at TAD borders than genome background (Berthelot et al. 2015). Further experiments designed to characterize recurrent breakpoints across the genome would be necessary to clarify the above hypotheses.

We also found evidence for selection acting on structural genomic variants at TAD boundaries. Recent studies have found that deletions are depleted at TAD boundaries in human populations (Sadowski et al. 2019) and in humans’ close relatives, apes (Fudenberg and Pollard 2019), as well as in cancer genomes (Akdemir et al. 2020). In *Drosophila*, we observed the same pattern for deletions and non-TE insertions, suggesting that these common SVs are subject to purifying selection. Unlike the above two types of SVs, patterns of TE insertions are not only different in their classes, but differ in evolutionary timescales as well. In polymorphic SV datasets, LTRs and LINEs are strongly depleted at TAD boundaries, whereas DNA-type TE is slightly enriched at TAD boundaries (Supplemental Table S15), suggesting they are under different selective pressure. Furthermore, such patterns were not observed in the divergence SV dataset, possibly because most of the older deleterious TE insertions have already been eliminated uniformly across the genomes of the *D. simulans* complex species. TAD boundaries appear to appear largely in gene-dense, chromatin-accessible, and transcribed regions where enriched in active chromatin marks (Szabo et al. 2019). For example, ∼77% of TAD boundaries annotated in *D. melanogaster* overlap promoters (Ramírez et al. 2018). Thus, it remains unclear which functional aspects are principal factors governing constraints of SVs at TAD boundaries.

The finding that divergence of tandem duplications is elevated at TAD boundaries relative to elsewhere when both are normalized by levels of polymorphism (Fig. 6G), suggesting that tandem duplicates at TAD boundaries are fixed at higher rates, though this imbalance in the odds ratio could also stem from a deviation in any of the four terms. The absence of enrichment of tandem duplications at TAD boundaries in the polymorphism data suggests that this finding is unlikely to be mutationally driven. Therefore, we propose that adaptation could be driving up the divergence of tandem duplicates at TAD boundaries. One intuitive reason for this suggestion may be that duplicated boundary sequences, such as insulator binding sites, may strengthen topological domain borders, thereby reinforcing the stability of chromatin domains. This is consistent with the hypothesis that duplications may be an important evolutionary mechanism of spatial genome organization (Sadowski et al. 2019). Similarly, the fact that TAD boundary duplications largely overlap with functional regulatory elements and genes argues for further examining the forces shaping enrichment of tandem duplications at TAD boundaries. Collectively, our findings offer novel insight into the evolutionary significance of spatial genome organization in shaping patterns of large-scale chromosomal rearrangements, common structural variants, and gene expression.

## Methods

### Fly strain and genome sequencing

The sequenced *D. pseudoobscura* strain (MV-25-SWS-2005) was initially collected at Mesa Verde, Colorado (Lat 37d 18’ 0” N, Long. 108d 24’ 58” W) in July 2005 by Stephen W. Schaeffer. The strain was subsequently inbred. DNA was extracted from adult females following a previously published protocol (Chakraborty et al. 2016). DNA was sheared using 21 gauge needles and size selected using the 30-80 kbp cutoff on Blue Pippin (Sage Science). Size selected DNA was sequenced on 10 SMRT cells using the Pacific Biosciences Sequel Platform. Illumina paired end (2 × 150 bp) reads were generated on HiSeq 4000 using the same DNA that was used for PacBio sequencing. PacBio long reads were assembled with Canu v1.7(Koren et al. 2017). After removal of redundant contigs and gap filling using raw reads with finisherSC (Lam et al. 2015), the assembly was polished twice with Arrow (Smrtanalysis 5.1.0) and three times with Pilon (Walker et al. 2014). Transposable elements were annotated using the EDTA pipeline (Ou et al. 2019). Gene models were annotated using MAKER (version 2.31.8) (Campbell et al. 2014). More details described in the Supplemental Methods.

### Hi-C experiments

Hi-C experiments were performed by Arima Genomics (https://arimagenomics.com/) with adult female flies according to the Arima-HiC protocol described in the Arima-HiC kit (P/N : A510008) with minor modifications to the crosslinking protocol (for details, see Supplemental Methods).

### Hi-C data processing and TAD annotation

Juicer (Durand et al. 2016) and HiCExplorer (Ramírez et al, 2018) were used to process Hi-C data from raw reads to interaction maps. Arrowhead from Juicer package, Armatus (Filippova et al. 2014), and HiCExplorer were used to annotate TADs each with different combinations of parameters. The output was compared and inspected visually based on chromatin interaction maps (Supplemental Fig. S16) using HiCPlotter (Akdemir and Chin 2015) to determine the optimal parameters (Supplemental Table S21).

### ChIP-seq and ATAC-seq data analysis

All short read alignments were performed against our *D. pseudoobscura* genome using Bowtie2 v2.2.7 (Langmead and Salzberg 2012). ChIP-seq peak calling was performed using MACS2 (version 2.0.10) (Zhang et al. 2008) with the default parameters. ChIP-seq normalization was performed using *bamCompare* from the deepTools suite (version 3.2.1) (Ramírez et al. 2016) with the following setting: ‘--binSize 10 --operation log_2_ --minMappingQuality 30 --skipNonCoveredRegions --ignoreDuplicates’. Read coverage of ATAC-seq was computed using deepTools *bamCoverage* for a bin size of 10 bp. To generate metaregions plots (Fig1. D-F) of ChIP-seq/ATAC-seq signals or frequency of insulator binding sites surrounding TAD boundaries, a matrix A_ij_ was generated for each dataset using deepTools *computeMatrix* and Perl scripts, in which each row represents a boundary and each column (j ∈ [-40,40]) represents the signal value in a 1 kbp non-overlapping bin within 40 kbp of the downstream and upstream flanking regions of that boundary. For CTCF and BEAF-32 binding sites, we summed values from columns of *A*_*ij*_ into a vector in which each element represents the signal value for the corresponding 1-kbp bin. For ChIP-seq and ATAC-seq data, we averaged values from columns of *A*_*ij*_ into a vector. To assess the significance of each signal at TAD boundaries, we generated 10,000 random samples of simulated TAD boundaries with the number and chromosome distribution confined by the observed dataset using BEDTools *shuffle* (version 2.25.0) (Quinlan and Hall 2010). We then computed the sampling distribution of each signal value around TAD boundaries in the same way as described above for actual boundaries and determined the *p*-values.

The preprocessed ChIP data for *D. melanogaster* were obtained from the modENCODE Consortium (http://www.modencode.org/)(modENCODE Consortium et al. 2010).

### Identification of conserved TAD features and significance tests

To identify conserved TAD features (i.e. body and boundary), genomic coordinates were converted between species using the UCSC liftOver tool and the *Dmel*-*Dpse* chain file generated in this study. To be successfully lifted over, features in one species require a 25% minimum ratio of bases (-minMatch=0.25) for body and one third for boundary (-minMatch=0.33) to be remapped in the other species and the size difference should not exceed 50% for body and 100% for boundary. Conserved TAD bodies were determined using BEDTools *intersect* with the parameters: -F 0.8 -f 0.8, which requires at least 80% reciprocal overlap in the corresponding intervals in both species. For boundaries, we considered any overlap as indicative of conservation.

To determine if the observed conservation of TAD features is statistically significant, we tested two null hypotheses. First, we assumed that the locations of TAD features across the genome are completely independent between separate species. To test this, we simulated 10,000 random samples of TAD features in one species and computed the sampling distribution of conservation with the other species. The *p*-values were then determined by the permutation distributions or Fisher Exact tests based on the observed and expected (mean of the 10,000 simulations) number of lifted and conserved TAD features. As an alternative null hypothesis for the TAD body, we assumed that TADs are completely conserved across the genome between species only when chromosomal rearrangements can disrupt them. To test this, we simulated 10,000 sets of genome shuffling by random fragmentation in each species. The size distribution of each sample of genome fragments requires to match the actual synteny blocks between *D. melanogaster* and *D. pseudoobscura*. A TAD from the first species was considered to be conserved if it is successfully lifted and at least 80% of the converted genomic coordinate overlap with any of the simulated genome fragments in the second species. Then, the sampling distribution of the conservation was used to determine the *p*-values.

### Gene expression data analysis

The preprocessed expression data were obtained from the Gene Expression Omnibus (https://www.ncbi.nlm.nih.gov/geo) database under accession ID GSE99574. Orthologs were obtained from FlyBase (https://flybase.org/) Orthologs gene sets. After filtering, 10,921 of the 13,638 *Dmel*-*Dpse* orthologs we retrieved are in a one-to-one relationship and have expression data. To measure expression divergence, we computed both Euclidean distance and Pearson’s correlation coefficient distance following the formulas as previously described (Pereira et al. 2009).

### Assembly-based structural variants detection

SV calling was performed following our custom pipeline (Kou et al. 2020; Liao et al. 2018) based on the LASTZ/CHAIN/NET workflow (Schwartz et al. 2003; Harris 2007) (see Supplemental Methods for a more detailed description of the pipeline). The pipeline is available on GitHub (https://github.com/yiliao1022/LASTZ_SV_pipeline).

### Identification and analysis of evolutionary chromosomal rearrangement breakpoints

Pairwise genome alignments were performed against *D. melanogaster* and *D. pseudoobscura* genome, respectively, using LASTZ (Version 1.04). The resulting alignments were then processed with axtChain/chainNet/netSyntenic tools to get the *netSyntenic* files which were used as input in our custom Perl script *Synbreaks*.*pl* to identify chromosomal rearrangement breakpoints. Breakpoints were classified into two categories: (1) synteny breaks if they were obtained from ‘top’, ‘syn’ or ‘NonSyn’ fills, and (2) inversion breaks if they were obtained from ‘inv’ fills. We excluded breaks which were identified from synteny blocks of size less than 10 kbp and near the terminal regions (<10 kbp) of long contigs in our analysis, because they are more likely introduced by assembling artifacts.

To quantify the distribution of rearrangement breakpoints along the TADs, we followed a previously described method (Krefting et al, 2018). Briefly, each TAD domain was extended by 50% of its size on each side and the resulting interval was subdivided into 20 equal-sized bins. The occurrence of breakpoints was then summed over bins for all TADs to generate a vector in which each element represents one of the 20 bins. Additionally, we generated 100 sets of random breakpoints as background control.

### Selection of structural variants at TAD boundaries

We measured the relative abundance of structural variants at TAD boundaries following a previously described method (Fudenberg and Pollard, 2019; Supplemental Methods). We also permuted 10,000 sets of TAD boundaries across the genome, excluding heterochromatic regions, to generate the background distribution of the relative abundance of SVs at TAD boundaries for statistical tests.

### Code availability

The code that reproduces analyses from the manuscript is available at GitHub (https://github.com/yiliao1022/TADEvoDrosophila).

### Data access

All raw and processed sequencing data generated in this study have been submitted to the NCBI BioProject database (https://www.ncbi.nlm.nih.gov/bioproject/) under accession number PRJNA596268.

### Competing interest statement

The authors declare no competing interests.

## Acknowledgements

This work was funded by the National Institutes of Health (R01GM123303), National Science Foundation (IOS-1656260), and startup funding from the University of California, Irvine to J.J.E. We thank the Genomics High Throughput Facility at University of California, Irvine (UCI) and Arima Genomics, Inc. San Diego for expert service. We would also like to thank Luna Thanh Ngo for her help with data collection and management. We thank Stephen W. Schaeffer for supplying the strain sequenced in the study.

## Author contributions

Y.L. and J.J.E. conceived the study and designed the research. M.C. contributed to the genomic data collection. M.C. and Y.L. generated the genome assembly. Y.L., X.Z. and J.J.E. analyzed the data and interpreted the results. Y.L. wrote the original draft. All authors reviewed and edited the manuscript.

## Notes

### Competing Interest Statement

The authors have declared no competing interest.

### Summary of Updates

Newly submitted revision

