## Supplemental Material for "Topologically associating domains and their role in the evolution of genome structure and function in *Drosophila*"

#### Supplemental Methods

##### QV estimation and BUSCO validation for the *D. pseudoobscura* assembly

We applied the method of Koren et al. (Koren et al. 2018) to the polished, pre-scaffolded assemblies to estimate base level error rates from the concordance between Illumina reads and the assembly of the same strain (i.e. QV).

Freebayes version v0.9.21 was run with the command:

```
freebayes -C 2 -0 -O -q 20 -z 0.10 -E 0 -X -u -p 2 -F 0.75 -b Dpseu.sorted.bam -v dpse.bayes.vcf -f Dpseudo_PacBioV2_genomic.fna
```

We calculated BUSCOs in our assembly with BUSCO V3 against the Diptera and Arthropoda database (Waterhouse et al. 2017).

##### Genome Annotation

***Simple Tandem repeats, centromeric and telomeric repeats identification*** Tandem repeats were identified using Tandem repeat finder (Benson 1999) with parameters “2 7 7 80 10 50 2000 -f -d -m”. The top 5 most abundant consensus sequences of tandemly repeated units were used as queries for further searching against the genome assembly to check their chromosomal distributions and the abundant frequency in the genome using RepeatMasker (http://www.repeatmasker.org/). Based on this information, we identified two centromere-specific tandem repeats in *D. pseudoobscura*, with the motif size of 170 bp and 21 bp, respectively. The 170-bp tandem repeat is only detected on the X and dot chromosome and is absent from its closely related species, *D. miranda*, suggesting the rapid turnover of centromere-specific repeats between species and even across chromosomes within species in *Drosophila*.

***Transposable elements annotation*** TE annotation was mainly performed using the EDTA pipeline. We also ran the following additional utilities to identify additional transposable elements in *D. pseudoobscura* genome. First, we applied RepeatModeler version 1.0.11 (Smit et al. 2015) for *de novo* repeat sequences identification with the default parameters. The resulting transposable elements library was filtered to exclude elements that overlapping with gene features based on Blastn. In total, RepeatModeler generated 924 candidate TE families, covering 1,149,638 bp. Next, we employed LTR_retriever pipeline (Ou and Jiang 2018) to specifically annotate long terminal repeat (LTR) retrotransposons in the *D. pseudoobscura* and *D. miranda* genomes. This pipeline outline is as follows: LTR retrotransposons were first identified using LTRharvest (Ellinghaus, Kurtz, and Willhoeft 2008) with parameters: -similar 90 -vic 10 -seed 20 -seqids yes -minlenltr 100 -maxlenltr 7000 -mintsd 4 -maxtsd 6 -motif TGCA -motifmis 1; then LTR_FINDER (Xu and Wang 2007) was run with parameters: -D 15000 -d 1000 -L 7000 -l 100 -p 20 -C -M 0. The output from the utilities above was combined and processed using LTR_retriever to obtain intact LTR-RT elements and construct a non-redundant LTRs library. Using LTR_retriever, we identified a total of 470 and 1,927 (of them, 1,406 located on the new Y chromosome) intact LTR-RT elements for *D. pseudoobscura* and *D. miranda*, respectively. We also employed MITE_Hunter (Han and Wessler 2010) and GRF https://github.com/bioinfolabmu/GenericRepeatFinder to identify Miniature Inverted repeat Transposable Elements (MITE). These two programs generated a total of 149 families of MITEs.

By searching the Repbase *Drosophila* TE library, the custom LTR retrotransposon library from LTR_retriever and the custom MITE sequences from MITE_Hunter and GRF, we were able to assign the TE type for 797/924 of RepeatModeler non-redundant repeat families, including 454 LTR retrotransposable elements and 343 DNA-type elements. The final custom TE library we generated for *D. pseudoobscura* integrates results from Repbase, LTR_retriver, the MITE annotation, and the RepeatModeler output.

***Gene annotation*** In this study, we performed *de novo* gene annotation for *D. miranda* (Mahajan et al. 2018) and *D. pseudoobscura* using Maker (version 2.31.8)(Campbell et al. 2014), following https://gist.github.com/darencard/bb1001ac1532dd4225b030cf0cd61ce2. For *D. miranda*, we assembled a transcriptome with data from using two publicly available RNA-seq datasets (Nozawa et al. 2016; Mahajan et al. 2018), composed of 12 and 21 samples, respectively. These samples cover a wide range of tissues and developmental stages for both male and female. RNA-seq reads from each sample were individually aligned to the genome assembly using HiSat2 (Kim et al. 2015) and assembled by StringTie (Pertea et al. 2015) with the default parameters. The resulting gtf files for each sample were then merged into a single file using the Stringtie --merge function. To prepare transcript evidence for

MAKER, the merged gtf files were converted to gff files using gffread. The transcriptome sequences were extracted accordingly using ReSeqTools (He et al. 2013) based on the gff file and reference genome. For protein evidence, the amino acid sequences from *D. melanogaster* (Flybase r6.26) and *D. pseudoobscura* (r3.04) were used.

For *D. pseudoobscura*, we used transcripts obtained from Flybase (r3.04) and a recent re-annotation project (Yang et al. 2018), each containing 23,456 and 39,527 transcripts, respectively. In addition, we generated the transcriptome using RNA-seq data from a previous study (Nozawa et al. 2016). Moreover, in order to achieve more accurate gene models and identify high-confidential alternative splice transcripts, a total of 15,372 and 22,237 Full-length Isoforms obtained from ISO-seq sequencing in male and female, respectively, were also integrated to guide gene annotation in the MAKER pipeline. Protein evidence used for *D. pseudoobscura* gene annotation includes translated sequences from *D. melanogaster* (Flybase r6.26) and protein sequences annotated for *D. miranda* in this study. The MAKER pipeline was run three times for each species.

##### Single-molecule RNA sequencing (Iso-seq) experiment and data analysis

Total RNA was extracted from the adult full bodies for males and females, respectively, using RNeasy Plus Mini Kit (Cat No./ID: 74134). cDNA synthesis and library preparation was performed at UC Irvine GHTF. One SMRT cell for each sex sample was sequenced using PacBio Sequel I. ISO-seq data was processed following the *IsoSeq* v3 pipeline which is available at https://github.com/PacificBiosciences/IsoSeq.

##### HiC Experiments

The Arima-HiC experiment was performed as follows: First, adult female flies were crosslinked as whole animals using 2% formaldehyde. After crosslinking, flies were pulverized on dry ice with mortar and pestle and then subject to the Arima-HiC protocol described in the Arima-HiC kit. Briefly, pulverized crosslinked fly tissue was digested using a cocktail or restriction enzymes recognizing the GATC and GANTC motifs. Next, digested ends were labeled, proximally ligated, and then proximally-ligated DNA was purified. After the Arima-HiC protocol, Illumina-compatible sequencing libraries were prepared by first shearing purified Arima-HiC proximally-ligated DNA and then size-selecting ~400bp DNA fragments using SPRI beads. The size-selected fragments containing ligation junctions were enriched using Enrichment Beads provided in the Arima-HiC kit, and converted into Illumina-compatible sequencing libraries using the KAPA Hyper Prep kit (P/N: KK8504) reagents. After unique dual index adapter ligation, DNA was PCR amplified and purified using SPRI beads. The purified DNA underwent standard QC (qPCR and Bioanalyzer) and sequenced on the HiSeq X following the manufacturer's protocols.

##### Gene Ontology analysis and identification of constitutive genes

Gene Ontology analysis was performed using the online tool available at the GO project website http://geneontology.org/. To identify constitutive genes in *D. melanogaster*, we used the expression data obtained from 8 tissues for both sex in two strains (w1118 and oreR) for *D. melanogaster* (Yang et al. 2018). We defined constitutive genes as those genes have expression level larger than 5 (gene-level DESeq2 normalized read counts) in all tissues (8), repilcations (4), sex (2), and strains (2). In total, 5854 genes meet this criteria.

##### Assembly-based structural variants detection pipeline

Our assembly-based structural variant detection pipeline includes four key steps: (1) Soft masked query genomes are aligned against the reference genome using LASTZ (Version 1.04). (2) The resulting Axt alignment files from step 1 are used to build long genome alignment chains (i.e. connect alignments if they are close enough) using axtChain. The chain files were sorted and merged into a single file using chainMergeSort if necessary. Next, a filtering procedure was applied to remove the low-quality chains using chainPreNet and the keeping chains were then used to get alignment nets by running chainNet. Finally, synteny information was added using netSyntenic. (3) SV calling for each pairwise genome comparison was performed using our custom Perl scripts based on the final syntenic format file from step 2. The SV output includes insertion, deletion, tandem duplication, inversion and complex SVs. (4) Population-scale genotyping of SVs was performed using custom Perl scripts. The custom Perl scripts for this pipeline are available at https://github.com/yiliao1022/LASTZ_SV_pipeline and https://github.com/yiliao1022/TADEvoDrosophila/tree/master/5_SVs_Calling.

#### Evaluation of the relative abundance of SVs at TAD boundaries

We measured the relative abundance of structural variants at TAD boundaries following a previously described method (Fudenberg and Pollard 2019). The method compares the observed number of SVs and the amount of sequences these SVs affected within the genomic regions where annotated as TAD boundaries with a uniform genome-wide expectation, which imposes the simplifying assumption that the SV mutation rate is fairly similar across the genome. In total, 8.62 Mb (4 kbp plus 2,156) was annotated as TAD boundary regions in *D. melanogaster*. For analysis in *D. pseudoobscura*, we used the 5-kbp HiCExplorer boundaries**.** The observed/expected count and base coverage of SVs were calculated according to the formula: $(\sum_{i \in k} N_i)/(N_{total} \sum_{i \in k} \frac{S_i}{S_{total}})$ , where i indexes genomic regions annotated as TAD boundaries, $S_{total}$ is the genome size, and $N_{total}$ is the total number and base coverage of each type of SV genomewide. SVs that fall into the heterochromatin and centromeric regions were excluded from the analysis, as these regions may be more prone to variants artifacts.

### Supplemental Tables

**Supplemental Table S1:** Sequencing data for *D. pseudoobscura* genome assembling.

| PacBio | | | Illumina paired-end | | | Hi-C | | |
|---|---|---|---|---|---|---|---|---|
| Reads | Mean Length | Depth | Pairs | Length | Depth | Pairs | Length | Depth |
| 3,170,985 | 14,493 bp | ~280 | 154,932,139 | 150bp | ~283 | 397,020,516 | 150bp | ~726 |

Depth is calculated based on the genome size of 164 MB.

**Supplemental Table S2:** Assembly statistics for the *D. pseudoobscura* genome.

| Features | Scaffold | Contig |
|---|---|---|
| Number | 70 | 72 |
| Total Bases | 163,274,969 | 163,282,969 |
| N10 | 68,158,638 | 35,530,770 |
| N30 | 68,158,638 | 32,422,566 |
| N50 | 32,422,566 | 30,706,867 |
| N70 | 30,706,867 | 23,510,042 |
| N80 | 30,706,867 | 22,902,128 |
| N90 | 23,510,042 | 9,717,740 |
| Min | 1,117 | 1,117 |
| Max | 68,158,638 | 35,530,770 |

| | Length | Contig Number | NCBI accession |
|---|---|---|---|
| Chr2 | 32,422,566 | 1 | CM020868.1 |
| Chr3 | 23,510,042 | 1 | CM020869.1 |
| Chr4 | 30,706,867 | 1 | CM020870.1 |
| Chr5 | 1,881,070 | 1 | CM020871.1 |
| ChrX | 68,158,638 | 3 | CM020872.1 |
| Mitochondrion | 16,118 | 1 | CM020873.2 |
| Unplace | 6,587,668 | 64 | - |

**Supplemental Table S3:** Quality evaluation of the current *D. pseudoobscura* genome assembly.

| | | | |
|---|---|---|---|
| **Quality Value (QV) = -10log$_{10}$(Probability of error)** | | 52 | |
| | | **Diptera_odb9** | **Arthropoda_odb9** |
| **BUSCO score** | Complete BUSCOs (C) | 2731 (97.6%) | 1061 (99.5%) |
| | Complete and single-copy BUSCOs (S) | 2700 (96.5%) | 1047 (98.2%) |
| | Fragmented BUSCOs (F) | 31 (1.1%) | 14 (1.3%) |
| | Missing BUSCOs (M) | 42 (1.5%) | 1 (0.1%) |
| | | 26 (0.9%) | 4 (0.4%) |
| | Total BUSCO groups searched | 2799 | 1066 |

**Supplemental Table S4:** Sequence composition of the 64 unplaced contigs.

| | | | |
|---|---|---|---|
| **TE occupancy** | Centromeric repeats:<br>Simple repeats:<br>Small RNA:<br>Retroelements:<br>DNA transposon:<br>Unclassified: | 3,639,793 bp<br>71,045 bp<br>263,447 bp<br>2,211,064 bp<br>26,375 bp<br>453,477 bp | 54 %<br>1 %<br>4 %<br>33 %<br>0.4 %<br>6.8 % |
| **Contigs contains centromere-specific tandem repeats** | 170bp<br><br>WVEN01000009.1<br>WVEN01000011.1<br>WVEN01000012.1<br>WVEN01000013.1<br>WVEN01000014.1<br>WVEN01000015.1<br>WVEN01000016.1<br>WVEN01000018.1<br>WVEN01000020.1<br>WVEN01000022.1<br>WVEN01000028.1<br>WVEN01000041.1<br>WVEN01000042.1<br>WVEN01000044.1<br>WVEN01000045.1<br>WVEN01000046.1<br>WVEN01000047.1<br>WVEN01000048.1<br>WVEN01000049.1<br>WVEN01000050.1<br>WVEN01000054.1<br>WVEN01000055.1<br>WVEN01000056.1<br>WVEN01000057.1<br>WVEN01000058.1<br>WVEN01000065.1<br>WVEN01000066.1 | 21bp<br><br>WVEN01000008.1<br>WVEN01000019.1<br>WVEN01000021.1<br>WVEN01000023.1<br>WVEN01000029.1<br>WVEN01000060.1<br>WVEN01000061.1 | Both<br><br>WVEN01000026.1<br>WVEN01000043.1<br>WVEN01000059.1 |

Contig Names refer to NCBI GenBank assembly accession GCF_009870275.2

**Supplemental Table S5:** Comparison of assembly continuity for existing *D. pseudoobscura* genome assemblies.

| Continuity | | Flybase r3.04 | Dpse_4.0 (English et al. 2012) | (Bracewell et al. 2019) | (Miller et al. 2018) | This study |
|---|---|---|---|---|---|---|
| | (Strain) | (MV2-25) | (MV2-25) | - | (MV2-25) | (MV-25-SWS-2005) |
| N10 | (bp) | 568,313 | 31,355,180 | 20,319,488 | 7,334,445 | 35,530,770 |
| N30 | (bp) | 304,883 | 30,679,157 | 9,603,404 | 4,302,182 | 32,422,566 |
| N50 | (bp) | 156,571 | 26,005,469 | 5,757,666 | 2,983,193 | 30,706,867 |
| N70 | (bp) | 74,128 | 22,861,174 | 1,827,199 | 1,033,991 | 23,510,042 |

**Supplemental Table S6:** Gene and TE annotation for the *D. pseudoobscura* genome.

| Gene Models | | BUSCO score (Diptera_odb9) | |
|---|---|---|---|
| **Chromosome** | **Number** | | |
| Chr2 | 3,287 | C:95.0%[S:62.8%,D:32.2%],F:2.5%,M:2.5%,n:2799 | |
| Chr3 | 2,545 | | |
| Chr4 | 2,564 | 2658 | Complete BUSCOs (C) |
| Chr5 | 74 | 1757 | Complete and single-copy BUSCOs (S) |
| X | 4,292 | 901 | Complete and duplicated BUSCOs (D) |
| Mitochondrion | 13 | 70 | Fragmented BUSCOs (F) |
| Unplaced | 638 | 71 | Missing BUSCOs (M) |
| Total | 13,413 | 2799 | Total BUSCO groups searched |

**Repetitive sequences**

| | | |
|---|---|---|
| Centromeric repeats: | 4,851,739 bp | 2.97 % |
| Simple repeats: | 7,656,035 bp | 4.69 % |
| Retro-elements: | 22,773,268 bp | 13.95 % |
| DNA transposons: | 1,985,898 bp | 1.22 % |
| Unclassified | 6,660,145 bp | 4.08% |

**Supplemental Table S7:** RNA sequencing data used for gene annotation in the MAKER pipeline.

| Data | NCBI accession | Tissue | Reference |
|---|---|---|---|
| **Published RNA-seq** | From DRR055234 to DRR055275 | **Tissues including:** abdomens; Male accessory glands; abdomens without gonads; imaginal discs; heads; 3rd instar larvae; 3rd instar larvae without imaginal discs; Female ovaries; pupae; thoraxes; Male testes; adult; | (Nozawa et al. 2016) |
| **Published transcripts** | | | |
| 39,527 transcripts | GSE99574 | Details see GSE99574 | (Yang et al. 2018) |
| 23,456 transcripts | NA | NA | FlyBase r3.04 |
| **ISO-seq** | | | |
| 15,372 transcripts | PRJNA596268 | Male adult full body | This study |
| 22,237 transcripts | PRJNA596268 | Female adult full body | This study |

**Supplemental Table S8:** Hi-C data preprocessing.

| | HiCExplorer | | 3dDNA |
|---|---|---|---|
| **Pairs considered** | 397,020,516 | **Sequenced Read Pairs** | 397,020,516 |
| Pairs mappable, unique and high quality | 304,513,515 (76.70%) | Alignable (Normal+Chimeric Paired) | 337,748,332 (85.1%) |
| Pairs used (Hi-C Contacts) | 194,589,768 (**49.01%**) | Paired used (Hi-C Contacts) | 204,080,597 (**51.4%**) |
| Inter chromosomal | 9,275,009 (2.3%) | Inter chromosomal | 7,746,309 (2.0%) |
| Intra chromosomal | | Intra chromosomal | |
| Short range (<20 kbp) | 74,173,026 (18.7%) | Short range (<20 kbp) | 99,160,171 (25.0%) |
| Long range | 111,141,733 (28.0%) | Long range | 97,172,702 (24.5%) |

**Supplemental Table S9**: Synteny blocks between *D. melanogaster* (BDGP Release 6) and *D. pseudoobscura* (this study).

***Supplemental Table S9A:*** *D. melanogaster to D. pseudoobscura*

| | Intrachromosomal | | Interchromosomal | | |
|---|---|---|---|---|---|
| | Number of Syntenic blocks | Syntenic coverage | Number of Syntenic blocks | Syntenic coverage | Base Coverage |
| 2L | 163 | 19,489,042 | 1 | 81,462 | 16,062,093 |
| 2R | 157 | 17,105,335 | 1 | 16,682 | 14,630,878 |
| 3L | 152 | 18,854,583 | 5 | 69,854 | 16,580,574 |
| 3R | 210 | 24,793,351 | 0 | 0 | 21,277,942 |
| 4 | 29 | 895,323 | 0 | 0 | 406,947 |
| X | 217 | 15,483,353 | 50 | 3,243,056 | 12,146,805 |
| Total | 928 | 96,620,987 | 57 | 3,411,054 | 81,105,239 |

***Supplemental Table S9B:*** *D. pseudoobscura to D. melanogaster*

| | Intrachromosomal | | Interchromosomal | | |
|---|---|---|---|---|---|
| | Number of Syntenic blocks | Syntenic coverage | Number of Syntenic blocks | Syntenic coverage | Base Coverage |
| Chr2 | 210 | 26,818,840 | 6 | 129,583 | 22,275,738 |
| Chr3 | 157 | 17,119,684 | 0 | 0 | 15,090,175 |
| Chr4 | 163 | 23,133,097 | 1 | 4,662 | 17,083,083 |
| Chr5 | 29 | 846379 | 0 | 0 | 456,074 |
| XR | 152 | 20,096,070 | 50 | 4,026,659 | 19,857,440 |
| XL | 217 | 17,063,934 | 0 | 0 | 10,747,531 |
| Total | **928** | **105,078,004** | **57** | **4,160,904** | **85,510,041** |

**Supplemental Table S10**: TADs identified in different cell lines/tissues and tools.

| **Species/ cell lines** | | **Armatus** | | **HiCExporer** | | **Arrowhead** | **Ref.** |
|---|---|---|---|---|---|---|---|
| ***Dmel*** | Num. | Cov. | Num. | Cov. | Num. | Cov. | |
| Kc167 | 914 | 79 Mb | 933 | 131 Mb | 698 | 76 Mb | (Chathoth and Zabet 2019) |
| BG3 | 979 | 76 Mb | 956 | 131 Mb | 609 | 63 Mb | |
| S2 | 1,005 | 72 Mb | 1,107 | 132 Mb | 624 | 63 Mb | (Wang et al. 2018) |
| ***Dpse*** | | | | | | | |
| Full body | 858 | 95.8 Mb | 1013 | 146 Mb | 795 | 87.4 Mb | This study |

Only considered TAD size > 30 kbp for Armatus in *Dps*e and > 20 kbp in *Dmel*;
Num., Number of TADs;
Cov., genome coverage;

**Supplemental Table S11:** Permutation tests (n=10,000) for determining if the observed conservation rates were significant than expected.

| Features | Samples | Observation | Expectation[d] | 95% confidence interval | *P*-values |
|---|---|---|---|---|---|
| Body[a] | *Dmel* (Kc167) | 291/640 | 68 | 55 - 86 | $1 \times 10^{-4}$ |
| | *Dmel* (BG3) | 308/668 | 71 | 58 - 89 | $1 \times 10^{-4}$ |
| | *Dmel* (S2) | 325/792 | 74 | 60 - 92 | $1 \times 10^{-4}$ |
| | *Dpse* (WB) | 419/678 | 101 | 86 - 116 | $1 \times 10^{-4}$ |
| Body[b] | *Dmel* (Kc167) | 465 | 210 | 190 - 229 | $1 \times 10^{-4}$ |
| | *Dmel* (BG3) | 496 | 221 | 201 - 241 | $1 \times 10^{-4}$ |
| | *Dmel* (S2) | 601 | 280 | 257 - 302 | $1 \times 10^{-4}$ |
| | *Dpse* (WB) | 534 | 252 | 232 - 274 | $1 \times 10^{-4}$ |
| | | [C/L] | [C/L] | [C/L] | |
| Boundary[c] | *Dmel* (Kc167) | 331/683 | 87/774 | 70 - 105 / 752 - 796 | $2.2 \times 10^{-16}$ |
| | *Dmel* (BG3) | 339/722 | 90/801 | 73 - 108 / 779 - 822 | $2.2 \times 10^{-16}$ |
| | *Dmel* (S2) | 351/807 | 103/924 | 86 - 123 / 899 - 947 | $2.2 \times 10^{-16}$ |
| | *Dpse* (WB) | 494/768 | 135/757 | 114 -157 / 730 - 783 | $2.2 \times 10^{-16}$ |

[a] hypothesis first;

[b] alternative hypothesis;

[c]5 kbp boundary for *Dmel* and 10 kbp boundary for *Dpse*;

[d]Expectation was determined by the mean of the 10,000 simulated samples;

(*Dmel*) *D. melanogaster*;

(*Dpse*) *D. pseudoobscura*;

(WB) whole body;

(C/L) Conserved/lifted over TADs.

**Supplemental Table S12:** Summary of TAD annotation (HiCExplorer), lift and conservation between *D. melanogaster* (BDGP Release 6) and *D. pseudoobscura* (this study).

| TAD Features | Species (sample) | Total | Liftover success | Conserved[a] | Genome Cov. [C/L/T] (Mb) | *P* values |
|---|---|---|---|---|---|---|
| Body | *Dmel* (Kc167) | 933 | 640 (68.6%) | 182 (19.5%) | 27.9/88.2/131.4 | $1\times10^{-4}$ |
| | *Dmel* (BG3) | 964 | 668 (69.3%) | 179 (18.6%) | 28.5/90.0/131.3 | $1\times10^{-4}$ |
| | *Dmel* (S2) | 1,107 | 792 (71.5%) | 173 (15.6%) | 24.4/91.5/132.4 | $1\times10^{-4}$ |
| | *Dpse* (WB) | 1,013 | 678 (66.9%) | 284 (28.0%) | 43.6/90.5/145.6 | $1\times10^{-4}$ |

[a]Conservation was determined by **90%** reciprocal overlap cutoff;
(*Dmel*) *D. melanogaster*;
(*Dpse*) *D. pseudoobscura*;
(Genome Cov.) Genome coverage, which was calculated for total annotated TADs (T), successfully lifted TADs (L) and conserved TADs based on the genome of the original species;
*P* values were calculated using permutation tests (n=10,000).

**Supplemental Table S13:** Published ChIP-seq or ChIP-chip data sources used in this study.

| Species | Cell /tissue | Insulator proteins | Accession | Reference |
|---|---|---|---|---|
| *D. melanogaster* | S2 | BEAF32 | GSE20760 | (Schwartz et al. 2012) |
| *D. melanogaster* | S2 | CP190 | GSE20766 | Schwartz et al. 2012 |
| *D. melanogaster* | S2 | CTCF | GSE32818 | Riddle et al. 2011) |
| *D. melanogaster* | S2 | Chromator | GSE20765 | modENCODE |
| *D. melanogaster* | S2 | Su(Hw) | GSE32813 | (Riddle et al. 2011) |
| *D. melanogaster* | S2 | Trl | GSE32822 | Riddle et al. 2011 |
| *D. melanogaster* | Kc167 | BEAF32 | GSM762845 | (Wood et al. 2011) |
| *D. melanogaster* | Kc167 | CP190 | GSM762836 | Wood et al. 2011 |
| *D. melanogaster* | Kc167 | CTCF | GSM1535983 | Wood et al. 2011 |
| *D. melanogaster* | Kc167 | Chromator | GSM1318357/GSE20763 | Wood et al. 2011 |
| *D. melanogaster* | Kc167 | Su(Hw) | GSM762839/GSE51964 | Wood et al. 2011 |
| *D. melanogaster* | Kc167 | Trl | modEncode_3245 | modENCODE |
| *D. melanogaster* | BG3 | BEAF32 | GSE20811 | Schwartz et al. 2012 |
| *D. melanogaster* | BG3 | CP190 | GSE20814 | Schwartz et al. 2012 |
| *D. melanogaster* | BG3 | CTCF | GSE20767 | Schwartz et al. 2012 |
| *D. melanogaster* | BG3 | Chromator | GSE20761 | modENCODE |
| *D. melanogaster* | BG3 | Su(Hw) | GSE20833 | Schwartz et al. 2012 |
| *D. melanogaster* | BG3 | Trl | GSE23466 | modENCODE |
| *D. pseudoobscura* | Embryos | BEAF-32 | - | (Yang et al. 2012) |
| *D. pseudoobscura* | WP | CTCF | - | (Ni et al. 2012) |

**Supplemental Table S14**: Evolutionary conservation of different TAD boundary classes between *D. melanogaster* and *D. pseudoobscura* (Supplemental to Figure 3).

***Supplemental Table S14A:*** *Genomic background expectation obtained from simulating 10,000 samples of randomized 10-kbp TAD boundaries.*

| Total number | Successfully lifted over | Expected conservation rate |
|---|---|---|
| 1673 | 1324 | 198/1324 = 15% |

***Supplemental Table S14B:*** *TAD boundaries that overlapped with distinct insulator binding sites.*

| Cell lines /Methods | Insulator | Conserved overlapped | Total | Percent | Conserved without overlapped | Total | Percent | *P*-values |
|---|---|---|---|---|---|---|---|---|
| **Kc167** | | | | | | | | |
| HiCExplorer | BEAF | 278 | 456 | 0.61 | 93 | 196 | 0.47 | 0.001875 |
| | CTCF | 163 | 273 | 0.60 | 208 | 379 | 0.55 | 0.2512 |
| | Chro | 325 | 532 | 0.61 | 46 | 120 | 0.38 | 8.783e-06 |
| | SuHw | 227 | 392 | 0.58 | 144 | 260 | 0.55 | 0.578 |
| | GAF | 124 | 210 | 0.59 | 247 | 442 | 0.56 | 0.4978 |
| | CP190 | 304 | 519 | 0.59 | 67 | 133 | 0.50 | 0.1084 |
| Juicer | BEAF | 252 | 401 | 0.63 | 367 | 662 | 0.55 | 0.02097 |
| | CTCF | 167 | 258 | 0.65 | 452 | 805 | 0.56 | 0.01832 |
| | Chro | 335 | 520 | 0.64 | 284 | 534 | 0.53 | 0.00027 |
| | SuHw | 272 | 452 | 0.60 | 347 | 611 | 0.57 | 0.2968 |
| | GAF | 119 | 221 | 0.54 | 500 | 842 | 0.59 | 0.1589 |
| | CP190 | 329 | 525 | 0.63 | 290 | 538 | 0.54 | 0.004593 |
| Armatus | BEAF | 534 | 747 | 0.71 | 315 | 635 | 0.50 | 2.2e-16 |
| | CTCF | 320 | 438 | 0.73 | 529 | 944 | 0.56 | 2.109e-09 |
| | Chro | 654 | 918 | 0.71 | 195 | 464 | 0.42 | 2.2e-16 |
| | SuHw | 481 | 727 | 0.66 | 368 | 655 | 0.56 | 0.000177 |
| | GAF | 270 | 404 | 0.67 | 579 | 978 | 0.59 | 0.009613 |
| | CP190 | 622 | 901 | 0.69 | 227 | 481 | 0.47 | 3.072e-15 |
| **BG3** | | | | | | | | |
| HiCExplorer | BEAF | 60 | 98 | 0.61 | 332 | 623 | 0.54 | 0.1749 |
| | CTCF | 78 | 154 | 0.51 | 314 | 567 | 0.55 | 0.3402 |
| | Chro | 349 | 602 | 0.58 | 43 | 119 | 0.36 | 1.957e-0 |
| | SuHw | 101 | 194 | 0.53 | 291 | 527 | 0.55 | 0.5027 |
| | GAF | 178 | 320 | 0.56 | 214 | 401 | 0.53 | 0.5964 |
| | CP190 | 254 | 428 | 0.59 | 138 | 293 | 0.47 | 0.001542 |
| Juicer | BEAF | 36 | 68 | 0.53 | 514 | 933 | 0.55 | 0.8276 |
| | CTCF | 102 | 169 | 0.60 | 448 | 832 | 0.54 | 0.1427 |
| | Chro | 320 | 546 | 0.59 | 230 | 455 | 0.51 | 0.01285 |
| | SuHw | 158 | 290 | 0.54 | 392 | 711 | 0.55 | 0.9063 |
| | GAF | 170 | 313 | 0.54 | 380 | 688 | 0.55 | 0.8395 |

| | | | | | | | | |
|---|---|---|---|---|---|---|---|---|
| | CP190 | 228 | 389 | 0.59 | 322 | 612 | 0.53 | 0.07285 |
| Armatus | BEAF | 102 | 149 | 0.68 | 772 | 1,352 | 0.57 | 0.009881 |
| | CTCF | 162 | 280 | 0.58 | 712 | 1,221 | 0.58 | 0.9424 |
| | Chro | 704 | 1061 | 0.66 | 170 | 440 | 0.39 | 2.2e-16 |
| | SuHw | 240 | 475 | 0.51 | 634 | 1,026 | 0.62 | 4.902e-05 |
| | GAF | 366 | 609 | 0.60 | 508 | 892 | 0.57 | 0.2457 |
| | CP190 | 497 | 778 | 0.64 | 377 | 723 | 0.52 | 5.239e-06 |
| **S2** | | | | | | | | |
| HiCExplorer | BEAF | 328 | 580 | 0.57 | 77 | 215 | 0.36 | 3.128e-07 |
| | CTCF | 49 | 103 | 0.48 | 356 | 692 | 0.51 | 0.5301 |
| | Chro | 367 | 648 | 0.57 | 38 | 147 | 0.26 | 2.941e-11 |
| | SuHw | 68 | 143 | 0.48 | 337 | 652 | 0.52 | 0.4218 |
| | GAF | 190 | 361 | 0.53 | 215 | 434 | 0.50 | 0.4254 |
| | CP190 | 336 | 620 | 0.54 | 69 | 175 | 0.39 | 0.000766 |
| Juicer | BEAF | 262 | 459 | 0.57 | 234 | 525 | 0.46 | 0.000118 |
| | CTCF | 51 | 101 | 0.50 | 445 | 883 | 0.50 | 1 |
| | Chro | 325 | 566 | 0.57 | 171 | 418 | 0.41 | 4.277e-07 |
| | SuHw | 90 | 183 | 0.49 | 406 | 801 | 0.51 | 0.7751 |
| | GAF | 137 | 279 | 0.49 | 359 | 705 | 0.51 | 0.6575 |
| | CP190 | 294 | 516 | 0.57 | 202 | 468 | 0.43 | 2.003e-05 |
| Armatus | BEAF | 594 | 904 | 0.66 | 215 | 563 | 0.38 | 2.2e-16 |
| | CTCF | 100 | 175 | 0.57 | 709 | 1,292 | 0.55 | 0.6278 |
| | Chro | 662 | 1021 | 0.65 | 147 | 446 | 0.33 | 2.2e-16 |
| | SuHw | 159 | 345 | 0.46 | 650 | 1,122 | 0.58 | 0.000141 |
| | GAF | 301 | 516 | 0.58 | 508 | 951 | 0.53 | 0.07964 |
| | CP190 | 629 | 1036 | 0.611 | 180 | 431 | 0.42 | 4.393e-11 |

**Supplemental Table S14C:** Stronger versus weaker TAD boundaries.

| | Total Num. of stronger borders | Num. of conserved strong borders | Percent | Total Num. of of weaker borders | Num. of conserved weaker borders | Percent | *P*-values |
|---|---|---|---|---|---|---|---|
| *Dmel* to *Dpse* | | | | | | | |
| Kc167 | 292 | 196 | 0.67 | 393 | 186 | 0.47 | 3.757e-07 |
| BG3 | 347 | 220 | 0.63 | 396 | 181 | 0.46 | 1.995e-06 |
| S2 | 430 | 261 | 0.61 | 394 | 153 | 0.39 | 5.617e-10 |
| *Dpse* to *Dmel* | | | | | | | |
| Kc167 | 367 | 235 | 0.64 | 326 | 130 | 0.40 | 3.373e-10 |
| BG3 | 367 | 225 | 0.61 | 326 | 132 | 0.40 | 6.782e-08 |
| S2 | 367 | 243 | 0.66 | 326 | 142 | 0.44 | 3.344e-09 |

***Supplemental Table S14D:*** *TAD boundaries that are shared across cell lines versus cell-specific boundaries.*

| Methods | Total Num.of shared borders | Conserved shared borders | Percent | Total Num. of cell-specific borders | Conserved cell-specific borders | Percent | *P*-values |
|---|---|---|---|---|---|---|---|
| Armatus | 1757 | 1180 | 0.67 | 943 | 405 | 0.43 | 2.2e-16 |
| Hicexplorer | 672 | 392 | 0.58 | 483 | 165 | 0.34 | 8.309e-16 |
| Juicer | 1229 | 753 | 0.61 | 795 | 299 | 0.38 | 2.2e-16 |

***Supplemental Table S14E:*** *TAD boundary classes with different flanking chromatin modifications.*

| Chromatin combinations | Total Num. of boundary | Num. of conserved boundary | Percent | *P*-values |
|---|---|---|---|---|
| Active_Active | 659 | 262 | 0.40 | 0.002325[a] |
| Active_PcG | 342 | 152 | 0.44 | 8.119e-05[b] |
| Active_Inactive | 665 | 287 | 0.43 | 2.336e-05[c] |
| Inactive_Inactive | 233 | 73 | 0.31 | - |
| Inactive_PcG | 166 | 62 | 0.37 | - |
| PcG_PcG | 141 | 33 | 0.23 | - |
| Inactive_total | 540 | 168 | 0.31 | - |
| Total | 2245 | 881 | 0.39 | - |
| Randomization test | 2245 | 476 | 0.21 | - |

Inactive_total = sum (Inactive_Inactive + Inactive_PcG + PcG_PcG);

[a]*P*-value was calculated for Active_Active versus Inactive_total;

[b]*P*-value was calculated for Active_PcG versus Inactive_total;

[c]*P*-value was calculated for Active_Inactive versus Inactive_total;

Note: *P*-values from Supplemental Table S14 A,B,C,D are determined by Chi-squared test.

**Supplemental Table S15:** Both strong and weak TAD boundaries are more frequently shared with boundaries that were identified across cell types, whereas weak TAD boundaries are more frequently overlapped with cell-specific boundaries than strong ones.

| | **Cell-specific** (752) | **Across-cell-types** (921) | ***P*-values** |
|---|---|---|---|
| **Kc167** | | | |
| Strong (413) | 59 | 354 | |
| Weak (576) | 151 | 392 | $2.802 \times 10^{-7}$ |
| **BG3** | | | |
| Strong (441) | 53 | 388 | |
| Weak (568) | 160 | 384 | $1.37 \times 10^{-11}$ |
| **S2** | | | |
| Strong (584) | 135 | 449 | |
| Weak (569) | 221 | 318 | $8.332 \times 10^{-11}$ |

*P*-values are determined by Fisher exact test.

**Supplementary Table S16:** 129 long (>50 kbp) coding genes coincided with full TADs in *D. melanogaster*.

| ***Dmel* geneID** | **Gene name** | **Chr** | **Start** | **End** | ***Dpse* ortholog** | ***Dpse* geneID** | **Chr** | **Start** | **End** |
|---|---|---|---|---|---|---|---|---|---|
| FBgn0000479 | | | | | | | | | |
| FBgn0003256 | | | | | | | | | |
| FBgn0004198 | | | | | | | | | |
| FBgn0052580 | | | | | | | | | |
| FBgn0085638 | | | | | | | | | |
| FBgn0266199 | | | | | | | | | |
| FBgn0267033 | | | | | | | | | |
| FBgn0267336 | | | | | | | | | |
| FBgn0284236 | | | | | | | | | |
| FBgn0003963 | ush | 2L | 476220 | 540560 | FBgn0075472 | LOC4817316 | NC_046681.1 | 18886568 | 18962765 |
| FBgn0041111 | lilli | 2L | 2885929 | 2954406 | FBgn0081331 | LOC4816631 | NC_046681.1 | 5205103 | 5303786 |
| FBgn0015600 | toc | 2L | 3068345 | 3144786 | FBgn0246841 | LOC6903567 | NC_046681.1 | 10160973 | 10211507 |
| FBgn0051774 | fred | 2L | 3902928 | 3991713 | FBgn0076481 | LOC4816819 | NC_046681.1 | 19828341 | 19951473 |
| FBgn0000547 | ed | 2L | 4031377 | 4115749 | FBgn0071799 | LOC4817240 | NC_046681.1 | 19639096 | 19758672 |
| FBgn0003716 | tkv | 2L | 5219000 | 5271384 | FBgn0072755 | LOC4817227 | NC_046681.1 | 24852925 | 24919964 |
| FBgn0085450 | Snoo | 2L | 7891187 | 7984196 | FBgn0080091 | LOC4817376 | NC_046681.1 | 23141739 | 23255965 |
| FBgn0011259 | Sema1a | 2L | 8542147 | 8672041 | FBgn0074948 | LOC4816208 | NC_046681.1 | 14892768 | 15012903 |
| FBgn0041092 | tai | 2L | 9166778 | 9250211 | FBgn0072098 | LOC4816259 | NC_046681.1 | 15388231 | 15481384 |
| FBgn0032151 | nAChRalpha6 | 2L | 9793317 | 9886250 | FBgn0077986 | LOC4817310 | NC_046681.1 | 18110288 | 18208112 |
| FBgn0051721 | Trim9 | 2L | 10544883 | 10628054 | FBgn0079380 | LOC4816752 | NC_046681.1 | 6696732 | 6769563 |
| FBgn0259176 | bun | 2L | 12455540 | 12546611 | FBgn0247162 | LOC6902896 | NC_046681.1 | 20992639 | 21111231 |
| FBgn0263354 | CG42784 | 2L | 13415431 | 13500427 | FBgn0246640 | LOC6903438 | NC_046681.1 | 12668130 | 12693940 |
| FBgn0259984 | kuz | 2L | 13550139 | 13639411 | FBgn0080133 | LOC4818148 | NC_046681.1 | 12755253 | 12871144 |
| FBgn0028509 | CenG1A | 2L | 13835564 | 13898712 | FBgn0076510 | LOC4816443 | NC_046681.1 | 3648729 | 3710205 |
| FBgn0261563 | wb | 2L | 14263027 | 14328262 | FBgn0073665 | LOC4816591 | NC_046681.1 | 4185195 | 4261530 |
| FBgn0003016 | osp | 2L | 14599196 | 14689340 | FBgn0077483 | LOC4818069 | NC_046681.1 | 11488589 | 11613073 |
| FBgn0028863 | CG4587 | 2L | 15644183 | 15718272 | FBgn0078284 | LOC4816419 | NC_046681.1 | 5020033 | 5066453 |
| FBgn0259735 | mtgo | 2L | 16545016 | 16663218 | FBgn0247420 | LOC6903125 | NC_046681.1 | 26136170 | 26267233 |
| FBgn0261804 | CG42750 | 2L | 18159811 | 18275770 | FBgn0080155 | LOC4817320 | NC_046681.1 | 22488603 | 22544365 |
| FBgn0058006 | CG40006 | 2L | 22622417 | 22756349 | FBgn0265242 | LOC6902757 | NC_046681.1 | 2952011 | 2974803 |
| FBgn0263780 | CG17684 | 2R | 1866080 | 2262115 | FBgn0246910 | LOC6903318 | NC_046681.1 | 17564336 | 17570127 |
| FBgn0260995 | dpr21 | 2R | 3066499 | 3191011 | FBgn0272884 | LOC26533897 | NC_046681.1 | 17771326 | 17783592 |
| FBgn0040849 | Ir41a | 2R | 4917519 | 5021955 | FBgn0245983 | LOC6898024 | NC_046680.1 | 3767355 | 3769951 |
| FBgn0053554 | Nipped-A | 2R | 5177965 | 5251022 | FBgn0245413 | LOC26533571 | NC_046680.1 | 1398559 | 1548443 |
| FBgn0000546 | EcR | 2R | 6087873 | 6169087 | FBgn0243551 | LOC6899016 | NC_046680.1 | 21585943 | 21671815 |
| FBgn0086655 | jing | 2R | 6502006 | 6621792 | FBgn0263814 | LOC6899052 | NC_046680.1 | 21772166 | 21902528 |
| FBgn0003090 | pk | 2R | 7150703 | 7223378 | FBgn0070804 | LOC4803477 | NC_046680.1 | 4966792 | 5020419 |

| | | | | | | | | | |
|---|---|---|---|---|---|---|---|---|---|
| FBgn0033438 | Mmp2 | 2R | 9611139 | 9683858 | FBgn0245614 | LOC6898791 | NC_046680.1 | 17885883 | 17960741 |
| FBgn0261698 | SLO2 | 2R | 10322195 | 10416996 | FBgn0071943 | LOC4804232 | NC_046680.1 | 10892329 | 10984572 |
| FBgn0263102 | psq | 2R | 10557888 | 10617280 | FBgn0264580 | LOC6898873 | NC_046680.1 | 19855085 | 19902889 |
| FBgn0053144 | CG33144 | 2R | 10740054 | 10798709 | FBgn0077331 | LOC4805321 | NC_046680.1 | 19716315 | 19755708 |
| FBgn0000633 | CG17716 | 2R | 13623044 | 13749367 | FBgn0074649 | LOC4805164 | NC_046680.1 | 18229569 | 18358516 |
| FBgn0265974 | ttv | 2R | 14526267 | 14587917 | FBgn0070143 | LOC4803961 | NC_046680.1 | 8788651 | 8853115 |
| FBgn0264089 | sli | 2R | 15869495 | 15922118 | FBgn0263810 | LOC4805561 | NC_046680.1 | 21507934 | 21562752 |
| FBgn0285917 | sbb | 2R | 18278198 | 18357296 | FBgn0245672 | LOC6898650 | NC_046680.1 | 15255540 | 15329626 |
| FBgn0003435 | sm | 2R | 19517864 | 19631508 | FBgn0264769 | LOC4804945 | NC_046680.1 | 16339804 | 16441091 |
| FBgn0010415 | Sdc | 2R | 21392402 | 21481176 | FBgn0070413 | LOC4805278 | NC_046680.1 | 19298122 | 19391643 |
| FBgn0266129 | lov | 2R | 25085780 | 25139627 | FBgn0265027 | LOC4804618 | NC_046680.1 | 13839121 | 13849504 |
| FBgn0035094 | CG9380 | 2R | 25183628 | 25255318 | FBgn0081730 | LOC4804620 | NC_046680.1 | 13921798 | 13927626 |
| FBgn0001316 | klar | 3L | 434083 | 540614 | FBgn0074321 | LOC117184056 | NC_046679.1 | 17232815 | 17233267 |
| FBgn0267487 | Ptp61F | 3L | 1342493 | 1475257 | FBgn0081581 | LOC4812118 | NC_046683.1 | 64577786 | 64585987 |
| FBgn0266696 | Svil | 3L | 2377655 | 2499226 | FBgn0076834 | LOC4812804 | NC_046683.1 | 55075034 | 55211476 |
| FBgn0263392 | Tet | 3L | 2786207 | 2879158 | FBgn0249681 | LOC6900088 | NC_046683.1 | 65557238 | 65659194 |
| FBgn0262593 | Shab | 3L | 2895017 | 2959579 | FBgn0249911 | LOC6900092 | NC_046683.1 | 65471782 | 65527489 |
| FBgn0262870 | axo | 3L | 4629604 | 4687317 | FBgn0074906 | LOC4812589 | NC_046683.1 | 59089541 | 59153919 |
| FBgn0035574 | RhoGEF64C | 3L | 4692783 | 4796253 | FBgn0249899 | LOC6900063 | NC_046683.1 | 59158565 | 59284682 |
| FBgn0085447 | sif | 3L | 5657913 | 5749599 | FBgn0078858 | LOC4813607 | NC_046683.1 | 50614964 | 50710357 |
| FBgn0020251 | sfl | 3L | 6495680 | 6550104 | FBgn0080991 | LOC4814130 | NC_046683.1 | 66608453 | 66672946 |
| FBgn0261788 | Ank2 | 3L | 7655389 | 7718395 | FBgn0245004 | LOC6900405 | NC_046683.1 | 44181432 | 44250541 |
| FBgn0016694 | Pdp1 | 3L | 7811458 | 7867369 | FBgn0249858 | LOC4812101 | NC_046683.1 | 57972383 | 58017080 |
| FBgn0011817 | nmo | 3L | 7979046 | 8054147 | FBgn0249744 | LOC6899982 | NC_046683.1 | 57785259 | 57853064 |
| FBgn0085385 | bma | 3L | 9908120 | 9967298 | FBgn0244878 | LOC6900645 | NC_046683.1 | 50839836 | 50897100 |
| FBgn0041096 | rols | 3L | 12008225 | 12064859 | FBgn0076690 | LOC4812739 | NC_046683.1 | 60074493 | 60128691 |
| FBgn0260941 | app | 3L | 12206238 | 12266841 | FBgn0079009 | LOC4813857 | NC_046683.1 | 45062249 | 45128414 |
| FBgn0264001 | bru3 | 3L | 13521907 | 13803400 | FBgn0262083 | LOC6900229 | NC_046683.1 | 62237844 | 62554451 |
| FBgn0087007 | bbg | 3L | 14412828 | 14536276 | FBgn0081889 | LOC4813512 | NC_046683.1 | 49738548 | 49857900 |
| FBgn0259175 | ome | 3L | 14672839 | 14747868 | FBgn0076726 | LOC4813899 | NC_046683.1 | 43509178 | 43577286 |
| FBgn0261090 | Sytbeta | 3L | 15014695 | 15064977 | FBgn0075740 | LOC4813079 | NC_046683.1 | 43181391 | 43225404 |
| FBgn0260943 | Rbp6 | 3L | 17058774 | 17238212 | FBgn0244751 | LOC6900901 | NC_046683.1 | 55683847 | 55875214 |
| FBgn0016797 | fz2 | 3L | 19140975 | 19235373 | FBgn0081984 | LOC4812483 | NC_046683.1 | 61926741 | 61929000 |
| FBgn0262737 | mub | 3L | 21844788 | 21932702 | FBgn0080346 | LOC4813190 | NC_046683.1 | 47711136 | 47763777 |
| FBgn0004449 | Ten-m | 3L | 22293044 | 22407863 | FBgn0246553 | LOC6903600 | NC_046683.1 | 56595766 | 56737063 |
| FBgn0266347 | nAChRalpha4 | 3L | 23206669 | 23287448 | FBgn0071670 | LOC4811728 | NC_046679.1 | 18978998 | 19036653 |
| FBgn0287185 | nvd | 3L | 24324866 | 24401655 | FBgn0272278 | LOC26533291 | NC_046679.1 | 19042190 | 19046707 |
| FBgn0267429 | lovit | 3L | 25236940 | 25670980 | FBgn0263149 | LOC6897476 | NC_046679.1 | 19205188 | 19215147 |
| FBgn0287183 | l(3)80Fg | 3L | 25798441 | 26014112 | FBgn0270973 | LOC26531986 | NC_046679.1 | 18926361 | 18949519 |

| | | | | | | | | | |
|---|---|---|---|---|---|---|---|---|---|
| FBgn0267430 | Pzl | 3R | 2554162 | 3263582 | FBgn0246990 | LOC6899548 | NC_046679.1 | 26030363 | 26054844 |
| FBgn0263346 | smash | 3R | 4659579 | 4712193 | FBgn0263296 | LOC6897751 | NC_046679.1 | 25020967 | 25036713 |
| FBgn0013576 | mtd | 3R | 5270116 | 5350443 | FBgn0248772 | LOC6897619 | NC_046679.1 | 22529751 | 22600472 |
| FBgn0083949 | side-III | 3R | 5862865 | 5959555 | FBgn0248644 | LOC6897387 | NC_046679.1 | 16350190 | 16436589 |
| FBgn0083963 | Nlg3 | 3R | 7570281 | 7639434 | FBgn0247584 | LOC6897868 | NC_046679.1 | 27586270 | 27676356 |
| FBgn0266801 | CG45263 | 3R | 8364328 | 8477617 | FBgn0076231 | LOC4802637 | NC_046679.1 | 24064667 | 24177705 |
| FBgn0261929 | 5-HT2B | 3R | 8578205 | 8629996 | FBgn0080752 | LOC4802031 | NC_046679.1 | 17586939 | 17634157 |
| FBgn0262614 | pyd | 3R | 8827068 | 8931896 | FBgn0263173 | LOC4802625 | NC_046679.1 | 23926197 | 24041582 |
| FBgn0003165 | pum | 3R | 9066343 | 9237682 | FBgn0263302 | LOC6899331 | NC_046679.1 | 25150068 | 25361033 |
| FBgn0001235 | hth | 3R | 10507561 | 10639568 | FBgn0074358 | LOC4801015 | NC_046679.1 | 7419554 | 7526298 |
| FBgn0037963 | Cad87A | 3R | 11893512 | 11949129 | FBgn0248851 | LOC6897810 | NC_046679.1 | 26173815 | 26206576 |
| FBgn0264493 | rdx | 3R | 13967061 | 14032057 | FBgn0082107 | LOC4801646 | NC_046679.1 | 13899676 | 13970122 |
| FBgn0285955 | cv-c | 3R | 14391683 | 14481713 | FBgn0076192 | LOC4801949 | NC_046679.1 | 16632194 | 16724764 |
| FBgn0263929 | jvl | 3R | 14741582 | 14795868 | FBgn0077532 | LOC4801060 | NC_046679.1 | 7994919 | 8070854 |
| FBgn0266756 | btsz | 3R | 14804092 | 14875488 | FBgn0248064 | LOC6897024 | NC_046679.1 | 8084537 | 8136265 |
| FBgn0259244 | CG42342 | 3R | 16502770 | 16571363 | FBgn0262050 | LOC6896920 | NC_046679.1 | 5650062 | 5712218 |
| FBgn0263995 | cpo | 3R | 17919832 | 18018892 | FBgn0076133 | LOC4803162 | NC_046679.1 | 29609529 | 29684944 |
| FBgn0004652 | fru | 3R | 18414273 | 18545586 | FBgn0262676 | LOC4801442 | NC_046679.1 | 12163419 | 12258033 |
| FBgn0263974 | qin | 3R | 18591027 | 18650137 | FBgn0072931 | LOC4801434 | NC_046679.1 | 11990290 | 12061241 |
| FBgn0003118 | pnt | 3R | 23290231 | 23346167 | FBgn0262662 | LOC6897112 | NC_046679.1 | 9813046 | 9884417 |
| FBgn0051145 | CG31145 | 3R | 23605751 | 23669659 | FBgn0248188 | LOC6896811 | NC_046679.1 | 3196002 | 3278400 |
| FBgn0004509 | Fur1 | 3R | 25347431 | 25473058 | FBgn0263364 | LOC6897901 | NC_046679.1 | 28168848 | 28337718 |
| FBgn0011666 | msi | 3R | 25516118 | 25608423 | FBgn0248760 | LOC6897597 | NC_046679.1 | 21943710 | 22050910 |
| FBgn0039431 | plum | 3R | 26470078 | 26535217 | FBgn0079629 | LOC4801081 | NC_046679.1 | 8225629 | 8289170 |
| FBgn0004842 | RYa-R | 3R | 26996980 | 27057935 | FBgn0262494 | LOC13036374 | NC_046679.1 | 8807721 | 8868130 |
| FBgn0004369 | Ptp99A | 3R | 29377646 | 29487131 | FBgn0263238 | LOC4800475 | NC_046679.1 | 2268638 | 2383280 |
| FBgn0266411 | sima | 3R | 30058311 | 30121798 | FBgn0080705 | LOC4800293 | NC_046679.1 | 729199 | 793552 |
| FBgn0010113 | hdc | 3R | 30277932 | 30372382 | FBgn0262041 | LOC6896726 | NC_046679.1 | 361603 | 444621 |
| FBgn0261988 | Gprk2 | 3R | 31405245 | 31457873 | FBgn0074786 | LOC4800727 | NC_046679.1 | 4750576 | 4790127 |
| FBgn0284435 | tyn | X | 142208 | 200663 | FBgn0244270 | LOC6901851 | NC_046683.1 | 22322415 | 22367122 |
| FBgn0052816 | CG32816 | X | 316576 | 470726 | FBgn0077175 | LOC4815282 | NC_046683.1 | 14038292 | 14061888 |
| FBgn0264449 | CG43867 | X | 808217 | 927975 | FBgn0243670 | LOC6901139 | NC_046683.1 | 39399697 | 39543032 |
| FBgn0025390 | Mur2B | X | 1523554 | 1668788 | FBgn0244397 | LOC6902079 | NC_046683.1 | 40512867 | 40519529 |
| FBgn0283741 | prage | X | 1773719 | 1853667 | FBgn0073294 | LOC4815930 | NC_046683.1 | 4167796 | 4202646 |
| FBgn0052791 | DIP-alpha | X | 3507245 | 3571649 | FBgn0244620 | LOC6901640 | NC_046683.1 | 9241195 | 9268292 |
| FBgn0266429 | AstA-R1 | X | 3574536 | 3666724 | FBgn0075512 | LOC4814982 | NC_046683.1 | 9393504 | 9418311 |
| FBgn0283657 | Tlk | X | 3720007 | 3789770 | FBgn0247156 | LOC6901632 | NC_046683.1 | 9488755 | 9586869 |
| FBgn0000635 | Fas2 | X | 4132887 | 4206093 | FBgn0077610 | LOC4814416 | NC_046683.1 | 39288156 | 39368197 |
| FBgn0029881 | pigs | X | 6600062 | 6653015 | FBgn0077823 | LOC4815731 | NC_046683.1 | 22102264 | 22159609 |

| | | | | | | | | | |
|---|---|---|---|---|---|---|---|---|---|
| FBgn0261873 | sdt | X | 8178549 | 8240474 | FBgn0077114 | LOC4814721 | NC_046683.1 | 14168679 | 14235710 |
| FBgn0261260 | mgl | X | 9358496 | 9500129 | FBgn0071481 | LOC4816002 | NC_046683.1 | 2983510 | 3010620 |
| FBgn0052698 | CARPB | X | 9783223 | 9855449 | FBgn0243764 | LOC6901062 | NC_046683.1 | 1115681 | 1163504 |
| FBgn0259170 | alpha-Man-Ia | X | 10266792 | 10325958 | FBgn0077085 | LOC4811663 | NC_046683.1 | 9116064 | 9182650 |
| FBgn0085443 | spri | X | 10490069 | 10585422 | FBgn0247809 | LOC6901478 | NC_046683.1 | 12339033 | 12451658 |
| FBgn0267001 | Ten-a | X | 12043698 | 12335214 | FBgn0077061 | LOC4815803 | NC_046683.1 | 42182177 | 42397715 |
| FBgn0259171 | Pde9 | X | 12785563 | 12897327 | FBgn0077055 | LOC4814257 | NC_046683.1 | 38534763 | 38676799 |
| FBgn0264078 | Flo2 | X | 14839376 | 14933946 | FBgn0244429 | LOC6902140 | NC_046683.1 | 4923878 | 5025517 |
| FBgn0000535 | eag | X | 14955477 | 15009989 | FBgn0244430 | LOC6902148 | NC_046683.1 | 4848589 | 4889314 |
| FBgn0285944 | para | X | 16455027 | 16533096 | FBgn0082095 | LOC4814623 | NC_046683.1 | 15180603 | 15252555 |
| FBgn0266354 | CG45002 | X | 16969069 | 17066732 | FBgn0248789 | LOC6901200 | NC_046683.1 | 17251805 | 17361961 |
| FBgn0003380 | Sh | X | 17924307 | 18063247 | FBgn0247929 | LOC6901448 | NC_046683.1 | 12898779 | 13029681 |
| FBgn0265598 | Bx | X | 18515428 | 18572809 | FBgn0247290 | LOC6902855 | NC_046681.1 | 20210853 | 20212694 |
| FBgn0264090 | CG43759 | X | 18930335 | 19028629 | FBgn0076989 | LOC4814891 | NC_046683.1 | 7320033 | 7439102 |

Note: Coordinates in *D. melanogaster* are based on reference genome version BDGP Release 6; Coordinates and gene IDs in *D. pseudoobscura* are based on the GenBand assembly accession GCA_009870125.2.

**Supplementary Table S17:** 73 long (>50 kbp) coding genes coincided with full TADs in *D. pseudoobscura*.

| ***Dpse*** | **Chr** | **Start** | **End** | **Flybase geneID** | ***Dmel* ortholog** | **Gene name** | **Chr** | **Start** | **End** |
|---|---|---|---|---|---|---|---|---|---|
| LOC6896811 | NC_046679.1 | 3196002 | 3278400 | FBgn0248188 | FBgn0051145 | CG31145 | 3R | 23605751 | 23669659 |
| LOC4800781 | NC_046679.1 | 5386959 | 5438514 | FBgn0077383 | FBgn0053208 | Mical | 3R | 10001554 | 10042895 |
| LOC6896916 | NC_046679.1 | 5461116 | 5578166 | | | | | | |
| LOC6896920 | NC_046679.1 | 5650062 | 5712218 | FBgn0262050 | FBgn0259244 | CG42342 | 3R | 16502770 | 16571363 |
| LOC4800948 | NC_046679.1 | 6763020 | 6834216 | FBgn0248101 | FBgn0037546 | mAChR-B | 3R | 8059284 | 8087049 |
| LOC4801003 | NC_046679.1 | 7189395 | 7246437 | FBgn0079305 | FBgn0004876 | cdi | 3R | 19044882 | 19094270 |
| LOC6897007 | NC_046679.1 | 7732765 | 7790727 | FBgn0248077 | FBgn0262617 | Nuak1 | 3R | 10264306 | 10308467 |
| LOC4801081 | NC_046679.1 | 8225629 | 8289170 | FBgn0079629 | FBgn0039431 | plum | 3R | 26470078 | 26535217 |
| LOC6897112 | NC_046679.1 | 9813046 | 9884417 | FBgn0262662 | FBgn0003118 | pnt | 3R | 23290231 | 23346167 |
| LOC4801646 | NC_046679.1 | 13899676 | 13970122 | FBgn0082107 | FBgn0264493 | rdx | 3R | 13967061 | 14032057 |
| LOC4801764 | NC_046679.1 | 14784567 | 14835704 | FBgn0247893 | FBgn0053100 | eIF4EHP | 3R | 24063077 | 24108960 |
| LOC4801843 | NC_046679.1 | 15359555 | 15533548 | FBgn0075943 | FBgn0011224 | heph | 3R | 31846987 | 32015520 |
| LOC6897387 | NC_046679.1 | 16350190 | 16436589 | FBgn0248644 | FBgn0083949 | side-III | 3R | 5862865 | 5959555 |
| LOC4801949 | NC_046679.1 | 16632194 | 16724764 | FBgn0076192 | FBgn0285955 | cv-c | 3R | 14391683 | 14481713 |
| LOC4802074 | NC_046679.1 | 17817447 | 17894364 | FBgn0243536 | FBgn0003944 | Ubx | 3R | 16656623 | 16734426 |
| LOC4802304 | NC_046679.1 | 21082351 | 21136455 | FBgn0261650 | FBgn0085386 | CG34357 | 3R | 4491943 | 4558067 |
| LOC4802625 | NC_046679.1 | 23926197 | 24041582 | FBgn0263173 | FBgn0262614 | pyd | 3R | 8827068 | 8931896 |
| LOC6899331 | NC_046679.1 | 25150068 | 25361033 | FBgn0263302 | FBgn0003165 | pum | 3R | 9066343 | 9237682 |
| LOC26534216 | NC_046679.1 | 26295215 | 26353745 | FBgn0247625 | FBgn0264357 | SNF4Agamma | 3R | 21140739 | 21214269 |
| LOC6897868 | NC_046679.1 | 27586270 | 27676356 | FBgn0247584 | FBgn0083963 | Nlg3 | 3R | 7570281 | 7639434 |
| LOC6897901 | NC_046679.1 | 28168848 | 28337718 | FBgn0263364 | FBgn0004509 | Fur1 | 3R | 25347431 | 25473058 |
| LOC4803056 | NC_046679.1 | 28349791 | 28404616 | FBgn0263363 | FBgn0262582 | cic | 3R | 20252770 | 20303942 |
| LOC6899355 | NC_046679.1 | 28625913 | 28734691 | FBgn0249366 | FBgn0038755 | Hs6st | 3R | 19929277 | 20008512 |
| LOC4803264 | NC_046679.1 | 30602446 | 30780262 | FBgn0263373 | FBgn0260003 | Dys | 3R | 19461085 | 19597288 |
| LOC4803646 | NC_046680.1 | 6343075 | 6410100 | FBgn0075887 | FBgn0040752 | Prosap | 2R | 14060456 | 14140902 |
| LOC4804232 | NC_046680.1 | 10892329 | 10984572 | FBgn0071943 | FBgn0261698 | SLO2 | 2R | 10322195 | 10416996 |
| LOC6898462 | NC_046680.1 | 11786375 | 11861598 | FBgn0245753 | FBgn0013733 | shot | 2R | 13864237 | 13942110 |
| LOC4804516 | NC_046680.1 | 12998411 | 13067112 | FBgn0072699 | FBgn0033405 | CG13954 | 2R | 9309296 | 9389467 |
| LOC6898564 | NC_046680.1 | 13825579 | 13905008 | FBgn0246243 | FBgn0054038 | CG34038 | 2R | 25167659 | 25169855 |
| LOC6898642 | NC_046680.1 | 15005173 | 15069436 | FBgn0245675 | FBgn0002643 | mam | 2R | 13991203 | 14060356 |
| LOC6898647 | NC_046680.1 | 15163374 | 15249936 | FBgn0245674 | FBgn0053958 | CG33958 | 2R | 18184382 | 18192851 |
| LOC4804945 | NC_046680.1 | 16339804 | 16441091 | FBgn0264769 | FBgn0003435 | sm | 2R | 19517864 | 19631508 |
| LOC6898791 | NC_046680.1 | 17885883 | 17960741 | FBgn0245614 | FBgn0033438 | Mmp2 | 2R | 9611139 | 9683858 |
| LOC4805263 | NC_046680.1 | 19176666 | 19251051 | FBgn0075833 | FBgn0041239 | Gr58a | 2R | 21979860 | 21981099 |
| LOC4805278 | NC_046680.1 | 19298122 | 19391643 | FBgn0070413 | FBgn0010415 | Sdc | 2R | 21392402 | 21481176 |
| LOC6898874 | NC_046680.1 | 19914625 | 19976308 | FBgn0264579 | FBgn0283521 | lola | 2R | 10481894 | 10543291 |
| LOC4805422 | NC_046680.1 | 20762914 | 20815098 | FBgn0072148 | FBgn0033667 | reb | 2R | 11904323 | 11912779 |
| LOC4805561 | NC_046680.1 | 21507934 | 21562752 | FBgn0263810 | FBgn0264089 | sli | 2R | 15869495 | 15922118 |
| LOC6899052 | NC_046680.1 | 21772166 | 21902528 | FBgn0263814 | FBgn0086655 | jing | 2R | 6502006 | 6621792 |
| LOC4805739 | NC_046680.1 | 23319395 | 23407484 | FBgn0080886 | FBgn0023441 | fus | 2R | 15657318 | 15676669 |
| LOC4816443 | NC_046681.1 | 3648729 | 3710205 | FBgn0076510 | FBgn0028509 | CenG1A | 2L | 13835564 | 13898712 |
| LOC4816591 | NC_046681.1 | 4185195 | 4261530 | FBgn0073665 | FBgn0261563 | wb | 2L | 14263027 | 14328262 |
| LOC6903561 | NC_046681.1 | 10275293 | 10366559 | FBgn0246567 | FBgn0016977 | spen | 2L | 159032 | 203397 |
| LOC4818069 | NC_046681.1 | 11488589 | 11613073 | FBgn0077483 | FBgn0003016 | osp | 2L | 14599196 | 14689340 |

| | | | | | | | | | |
|---|---|---|---|---|---|---|---|---|---|
| LOC4817896 | NC_046681.1 | 12699950 | 12749976 | FBgn0081619 | FBgn0023407 | B4 | 2L | 13500624 | 13549328 |
| LOC4818148 | NC_046681.1 | 12755253 | 12871144 | FBgn0080133 | FBgn0259984 | kuz | 2L | 13550139 | 13639411 |
| LOC26533542 | NC_046681.1 | 14293071 | 14349942 | | | | | | |
| LOC4816208 | NC_046681.1 | 14892768 | 15012903 | FBgn0074948 | FBgn0011259 | Sema1a | 2L | 8542147 | 8672041 |
| LOC4816259 | NC_046681.1 | 15388231 | 15481384 | FBgn0072098 | FBgn0041092 | tai | 2L | 9166778 | 9250211 |
| LOC6902405 | NC_046681.1 | 15579131 | 15732776 | FBgn0249459 | FBgn0000464 | Lar | 2L | 19586623 | 19732069 |
| LOC4817865 | NC_046681.1 | 16393823 | 16474646 | FBgn0076373 | FBgn0266521 | stai | 2L | 6100377 | 6124653 |
| LOC4817310 | NC_046681.1 | 18110288 | 18208112 | FBgn0077986 | FBgn0032151 | nAChRalpha6 | 2L | 9793317 | 9886250 |
| LOC4816903 | NC_046681.1 | 18235141 | 18286585 | FBgn0075064 | FBgn0028704 | Nckx30C | 2L | 9711512 | 9746495 |
| LOC4817316 | NC_046681.1 | 18886568 | 18962765 | FBgn0075472 | FBgn0003963 | ush | 2L | 476220 | 540560 |
| LOC4817453 | NC_046681.1 | 19120954 | 19177483 | FBgn0078274 | FBgn0004611 | Plc21C | 2L | 305935 | 355566 |
| LOC6902868 | NC_046681.1 | 20531988 | 20656719 | FBgn0272756 | | | | | |
| LOC6902896 | NC_046681.1 | 20992639 | 21111231 | FBgn0247162 | FBgn0259176 | bun | 2L | 12455540 | 12546611 |
| LOC4817376 | NC_046681.1 | 23141739 | 23255965 | FBgn0080091 | FBgn0085450 | Snoo | 2L | 7891187 | 7984196 |
| LOC6903125 | NC_046681.1 | 26136170 | 26267233 | FBgn0247420 | FBgn0259735 | mtgo | 2L | 16545016 | 16663218 |
| LOC4817542 | NC_046681.1 | 26739952 | 26812700 | FBgn0073723 | FBgn0053516 | dpr3 | 2L | 2058790 | 2109878 |
| LOC6903295 | NC_046681.1 | 29801894 | 29908866 | FBgn0249104 | FBgn0034106 | CG9068 | 2R | 16283435 | 16287450 |
| LOC4811795 | NC_046682.1 | 784082 | 839864 | FBgn0074773 | FBgn0085432 | pan | 4 | 69326 | 114270 |
| LOC4815782 | NC_046683.1 | 3474993 | 3530448 | FBgn0074848 | FBgn0004370 | Ptp10D | X | 11622015 | 11677338 |
| LOC6901756 | NC_046683.1 | 6654611 | 6736259 | FBgn0249613 | FBgn0029946 | CG15034 | X | 7346259 | 7347506 |
| LOC6901632 | NC_046683.1 | 9488755 | 9586869 | FBgn0247156 | FBgn0283657 | Tlk | X | 3720007 | 3789770 |
| LOC6901448 | NC_046683.1 | 12898779 | 13029681 | FBgn0247929 | FBgn0003380 | Sh | X | 17924307 | 18063247 |
| LOC26532449 | NC_046683.1 | 25938122 | 26018820 | | | | | | |
| LOC6901093 | NC_046683.1 | 38284146 | 38417666 | FBgn0243698 | FBgn0265597 | rad | X | 12988615 | 13079070 |
| LOC6901139 | NC_046683.1 | 39399697 | 39543032 | FBgn0243670 | FBgn0264449 | CG43867 | X | 808217 | 927975 |
| LOC6902018 | NC_046683.1 | 41866256 | 41951315 | FBgn0244155 | FBgn0264542 | hwt | X | 12516634 | 12568452 |
| LOC4813557 | NC_046683.1 | 45301101 | 45446946 | FBgn0076932 | FBgn0262509 | nrm | 3L | 22998372 | 23040300 |
| LOC6900681 | NC_046683.1 | 51484002 | 51610780 | FBgn0245227 | FBgn0052062 | Rbfox1 | 3L | 10481412 | 10594012 |
| LOC4812408 | NC_046683.1 | 61629954 | 61730871 | FBgn0076771 | FBgn0052206 | CG32206 | 3L | 19340321 | 19412912 |

Note: Coordinates in *D. melanogaster* are based on reference genome version BDGP Release 6; Coordinates and gene IDs in *D. pseudoobscura* are based on the GenBank assembly accession GCA_009870125.2.

**Supplemental Table S18**: Genome synteny breakpoints (within synteny blocks larger than 10 kbp) identified in the other 16 *Drosophila* genome assemblies relative to *D. melanogaster* and *D. pseudoobscura*, respectively.

| | Divergence Time (Mys) | Top fills | Nonsyn fills | Syntenic fills | Inv fills | Total syntenic breaks | Total inversion breaks |
|---|---|---|---|---|---|---|---|
| ***D. melanogaster*** **versus** | | | | | | | |
| *D. simulans* | 3.53 | 101 | 9 | 3 | 13 | 121 | 13 |
| *D. mauritiana* | 3.53 | 144 | 12 | 3 | 14 | 167 | 14 |
| *D. sechellia* | 3.53 | 123 | 8 | 4 | 10 | 143 | 10 |
| *D. erecta* | 7.25 | 90 | 6 | 4 | 22 | 108 | 22 |
| *D. yakuba* | 7.25 | 112 | 10 | 5 | 16 | 135 | 16 |
| *D. eugracilis* | 14.27 | 178 | 55 | 4 | 19 | 245 | 19 |
| *D. biarmipes* | 15.24 | 85 | 158 | 18 | 38 | 269 | 38 |
| *D. triauraria* | Unknown | 627 | 115 | 9 | 36 | 759 | 36 |
| *D. ananassae* | 34.37 | 189 | 237 | 59 | 106 | 493 | 106 |
| *D. bipectinata* | 34.37 | 540 | 219 | 7 | 22 | 774 | 22 |
| *D. pseudoobscura* | 49 | 169 | 9 | 246 | 314 | 432 | 314 |
| *D. persimilis* | 49 | 385 | 281 | 35 | 101 | 709 | 101 |
| *D. miranada* | 49 | 187 | 30 | 223 | 308 | 448 | 308 |
| *D. willistoni* | 62.1 | 1001 | 163 | 8 | 63 | 1180 | 63 |
| *D. mojavensis* | 72.63 | 879 | 123 | 16 | 58 | 1026 | 58 |
| *D. virilis* | 72.63 | 729 | 184 | 41 | 91 | 962 | 91 |
| ***D. pseudoobscura*** **versus** | | | | | | | |
| *D. simulans* | 49 | 232 | 324 | 94 | 144 | 678 | 172 |
| *D. mauritiana* | 49 | 265 | 286 | 101 | 164 | 681 | 193 |
| *D. sechellia* | 49 | 197 | 323 | 112 | 159 | 661 | 188 |
| *D. melanogaster* | 49 | 181 | 4 | 256 | 331 | 469 | 359 |
| *D. erecta* | 49 | 260 | 221 | 113 | 190 | 624 | 220 |
| *D. yakuba* | 49 | 352 | 317 | 49 | 115 | 748 | 145 |
| *D. eugracilis* | 49 | 555 | 290 | 16 | 54 | 889 | 82 |
| *D. biarmipes* | 49 | 353 | 392 | 44 | 105 | 819 | 135 |
| *D. triauraria* | 49 | 722 | 265 | 15 | 36 | 1032 | 66 |
| *D. ananassae* | 49 | 302 | 307 | 80 | 162 | 717 | 190 |
| *D. bipectinata* | 49 | 769 | 234 | 3 | 33 | 1033 | 60 |
| *D. persimilis* | 1.96 | 292 | 80 | 24 | 10 | 446 | 60 |
| *D. miranada* | 2.76 | 60 | 120 | 29 | 54 | 259 | 104 |
| *D. willistoni* | 62.1 | 1072 | 135 | 10 | 66 | 1242 | 91 |
| *D. mojavensis* | 72.63 | 941 | 95 | 21 | 50 | 1085 | 78 |
| *D. virilis* | 72.63 | 723 | 217 | 43 | 77 | 1010 | 104 |

The genome assemblies are obtained from (Miller et al. 2018; Mahajan et al. 2018).

**Supplementary Table S19:** Euchromatic regions defined in this study for *D. melanogaster* (BDGP Release 6).

| Chromosome | Coordinates |
|---|---|
| 2L | 1..22200000 |
| 2R | 5000000..25479258 |
| 3L | 1..23400000 |
| 3R | 4000000..31815305 |
| X | 1..21900000 |

**Supplemental Table S20:** Structural genomic variants in the polymorphic and divergence datasets used for testing the mode of selection operating upon them at TAD boundaries using Fudenberg and Pollard's method (Fudenberg and Pollard 2019) (Supplemental to Figure 6F).

| SV Types | Num. | Cov. (Mb) | Exp. breaks | Exp. Cov. (kb) | Obv. breaks | Exp. Cov. (kb) | Breakpoints: Log10(O/E) | Coverage: Log10(O/E) |
|---|---|---|---|---|---|---|---|---|
| ***Dmel*** | | | | | | | | |
| Total INS | 83,083 | 6.06 | 6,121 | 446.13 | 3,914 | 364.00 | -0.19 | -0.09 |
| 1-10 bp INS | 49,912 | 0.22 | 3,677 | 16.00 | 2,450 | 10.10 | -0.18 | -0.20 |
| 11bp - 20kb INS | 33,171 | 5.84 | 2,444 | 430.13 | 1,464 | 353.90 | -0.22 | -0.08 |
| Toal DEL | 158,075 | 1.93 | 23,290 | 142.38 | 19,044 | 94.034 | -0.09 | -0.18 |
| 1-10 bp DEL | 123,395 | 0.41 | 18181 | 30.08 | 15,135 | 23.51 | -0.08 | -0.11 |
| 11bp - 2kb DEL | 34,680 | 1.52 | 5109 | 112.30 | 3,909 | 70.53 | -0.12 | -0.20 |
| Toal TE | 7,132 | 32.75 | 525 | 2,412.43 | 405 | 1,514.60 | -0.11 | -0.20 |
| LTR | 3,536 | 23.80 | 260 | 1753.00 | 154 | 1005,38 | -0.23 | -0.24 |
| LINE | 2,242 | 6.95 | 165 | 511.73 | 124 | 333.25 | -0.12 | -0.19 |
| DNA-type | 1,352 | 2.00 | 100 | 147.61 | 127 | 175.98 | 0.11 | 0.08 |
| DUP (TD) | 1,803 | 3.16 | 266 | 232.80 | 181 | 152.43 | -0.17 | -0.18 |
| ***Dsim*** | | | | | | | | |
| Total INS | 285,409 | 4.76 | 21,026 | 350.82 | 18,269 | 372.90 | -0.06 | 0.03 |
| 1-10 bp INS | 184,329 | 0.76 | 13,579 | 56.24 | 11,287 | 48.09 | -0.08 | -0.07 |
| 11bp - 20kb INS | 101,080 | 4.00 | 7446 | 194.58 | 6,982 | 324.81 | -0.03 | 0.04 |
| Toal DEL | 510,798 | 4.14 | 75,259 | 305.17 | 68,093 | 272.629 | -0.04 | -0.05 |
| 1-10 bp DEL | 398,117 | 1.40 | 58,657 | 102.95 | 54,599 | 0.92 | -0.03 | -0.05 |
| 11bp - 2kb DEL | 112,681 | 2.74 | 16,602 | 202.22 | 13,494 | 181.70 | -0.09 | -0.05 |
| Toal TE | 1,816 | 3.73 | 134 | 274.71 | 120 | 254.72 | -0.05 | -0.03 |
| LTR | 396 | 2.07 | 29 | 152.61 | 28 | 130.47 | -0.02 | -0.07 |
| LINE | 527 | 1.06 | 39 | 77.70 | 34 | 62.00 | -0.06 | -0.10 |
| DNA-type | 902 | 0.60 | 66 | 44.45 | 60 | 62.49 | -0.04 | 0.15 |
| DUP (TD) | 1,291 | 1.40 | 190 | 103.38 | 287 | 177.56 | 0.18 | 0.23 |
| ***Dmira*** | | | | | | | | |
| DUP (TD) | 1,364 | 2.38 | 181 | 157.84 | 293 | 363.66 | 0.21 | 0.36 |

Note: We assigned two breakpoints for each deletion (DELs) or tandem duplication (DUP), and one breakpoint for each insertion (INS) or TE insertion.

**Supplemental Table 21:** Combinations of parameters tested for each TAD annotation tool.

| | |
|---|---|
| **HiCExplorer** | hicFindTADs --correctForMultipleTesting fdr<br><br>--m hic_corrected.h5 \| hic_corrected_5K.h5 \| hic_corrected_10K.h5 \| hic_corrected_20K.h5<br>--delta 0.01 \| 0.04<br>--minBoundaryDistance 5000 \| 20000<br>--thresholdComparisons 0.05 \| 0.01 \| 0.005 \| 0.001<br><br>**hicFindTADs -m hic_corrected_5K.h5 --correctForMultipleTesting fdr --delta 0.04 --minBoundaryDistance 20000 --thresholdComparisons 0.001** |
| **Juicer Arrowhead** | java -Xmx20g -jar /data/apps/juicer/1.5.6/scripts/juicer_tools.jar arrowhead -k NONE --ignore_sparsity inter_30.hic dpse_arrowhead<br><br>-r 5000 \| 10000<br><br>**java -Xmx20g -jar /data/apps/juicer/1.5.6/scripts/juicer_tools.jar arrowhead -k NONE -r 5000 --ignore_sparsity inter_30.hic dpse_arrowhead** |
| **Armatus** | Armatus -m -S<br>-r 5000 \| 10000 \| 20000 \| 25000 \| 40000 \| 50000<br>-s 0.1 \| 0.05<br>-g 1.0 \| 2.0 \| 0.9<br><br>**armatus -S -g 0.9 -r 5000 -s 0.1 -i 5K.matrix -o 5K.out** |

The optimal parameters for each tool are highlighted below and used in this study.

### Supplemental Figures

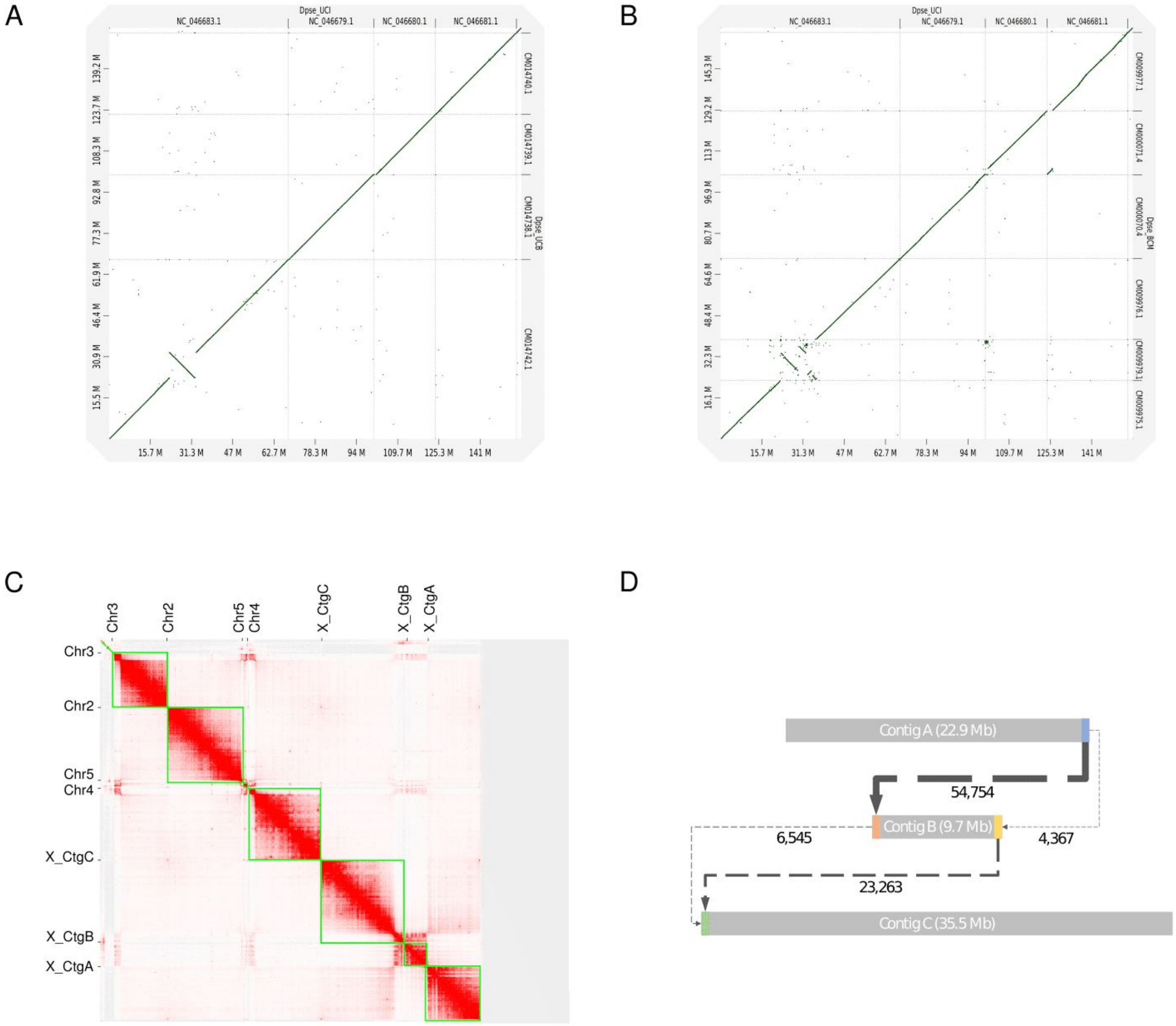


**Supplemental Figure S1:** A large pericentromeric inversion (9.7Mb) on the X-chromosome of *D. pseudoobscura* between the current assembly and the previous assemblies of this species. (A) Whole-genome alignment plot between the current assembly and the UCB assembly (Mahajan et al. 2018). (B) Whole-genome alignment plot between the current assembly and the FlyBase Dpse_4.0 assembly (English et al. 2012). (C) Hi-C contact map verified the highly continuous contigs and supported the scaffolding of the three contigs of the X-chromosome. (D) Local Hi-C contact frequency data further supported the scaffolding of the three contigs for the X-chromosome.

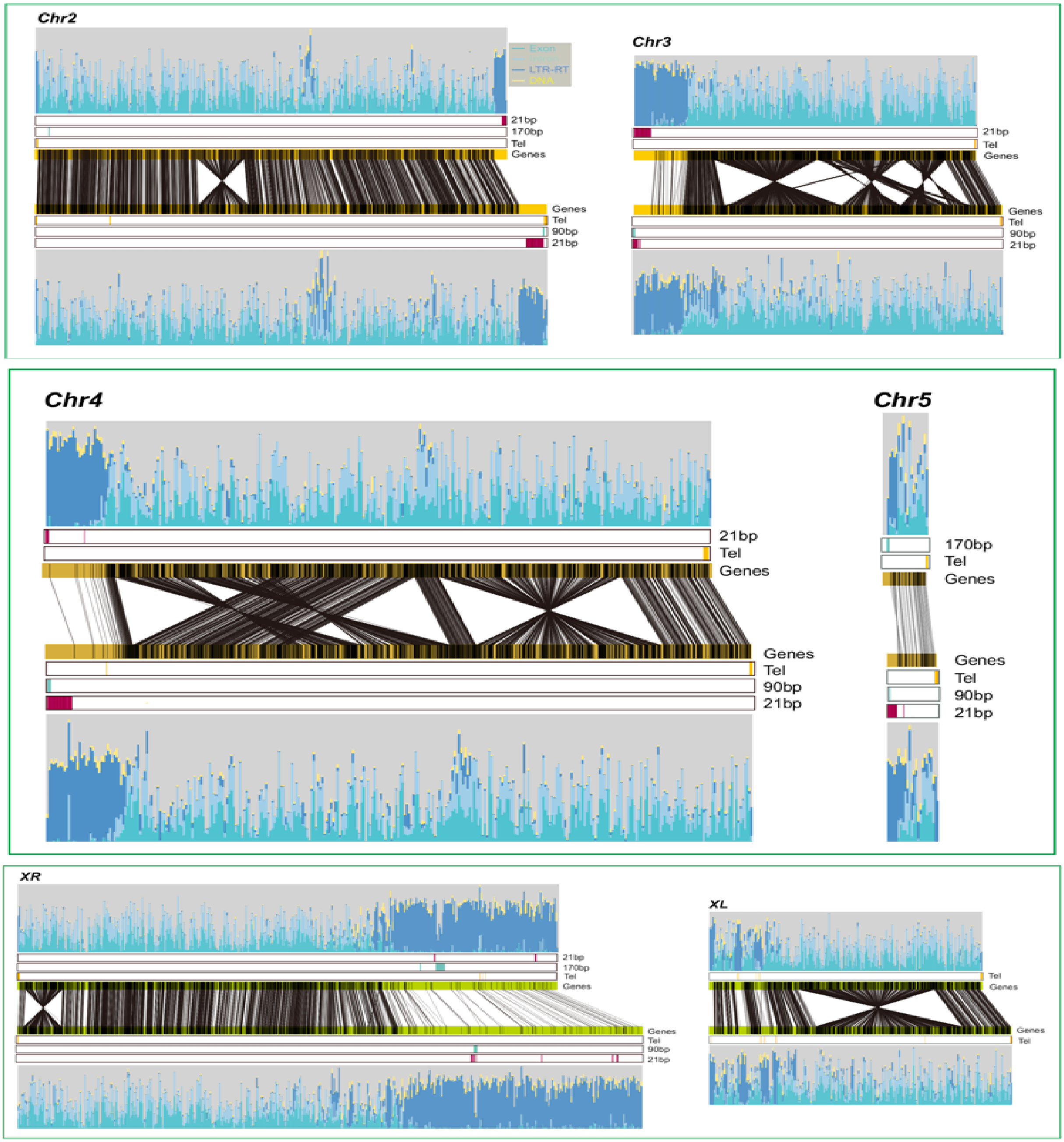


**Supplemental Figure S2:** Comparative genomic analysis of genome features between *D. pseudoobscura* (this study) and *D. miranda (Mahajan et al. 2018)*. Although their genomes were shuffled by several large rearrangements, the distribution of genomic features (i.e. repeat and gene content) is generally preserved.

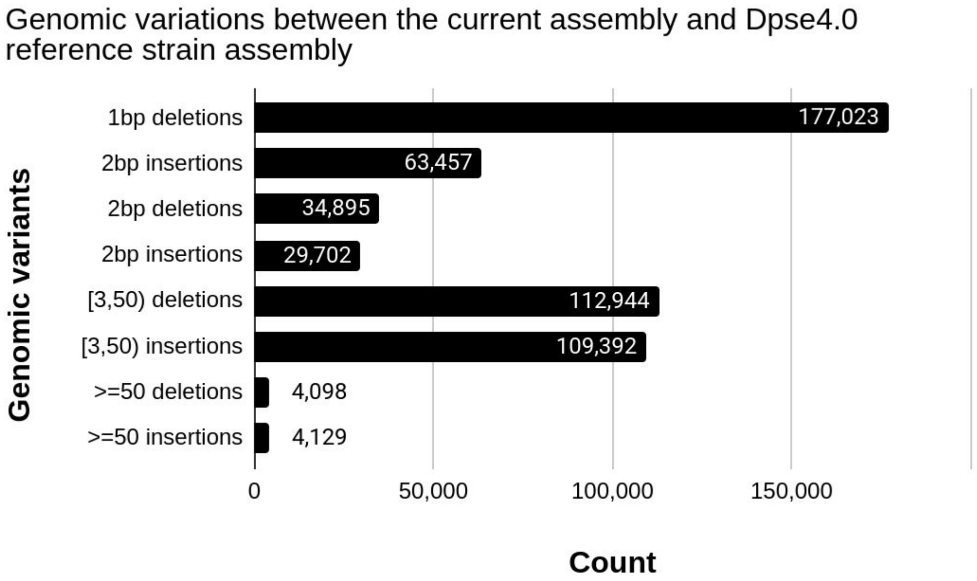


**Supplemental Figure S3:** Genomic variations identified between the current assembly and the *Dpse4.0* assembly from the reference strain MV2-25. The genomic variants were identified using Paftools (https://github.com/lh3/minimap2/tree/master/misc) within ~140 Mb alignable regions both with single coverage between assemblies. Additionally, there are a total of 1,434,692 SNPs were called between these two assemblies.

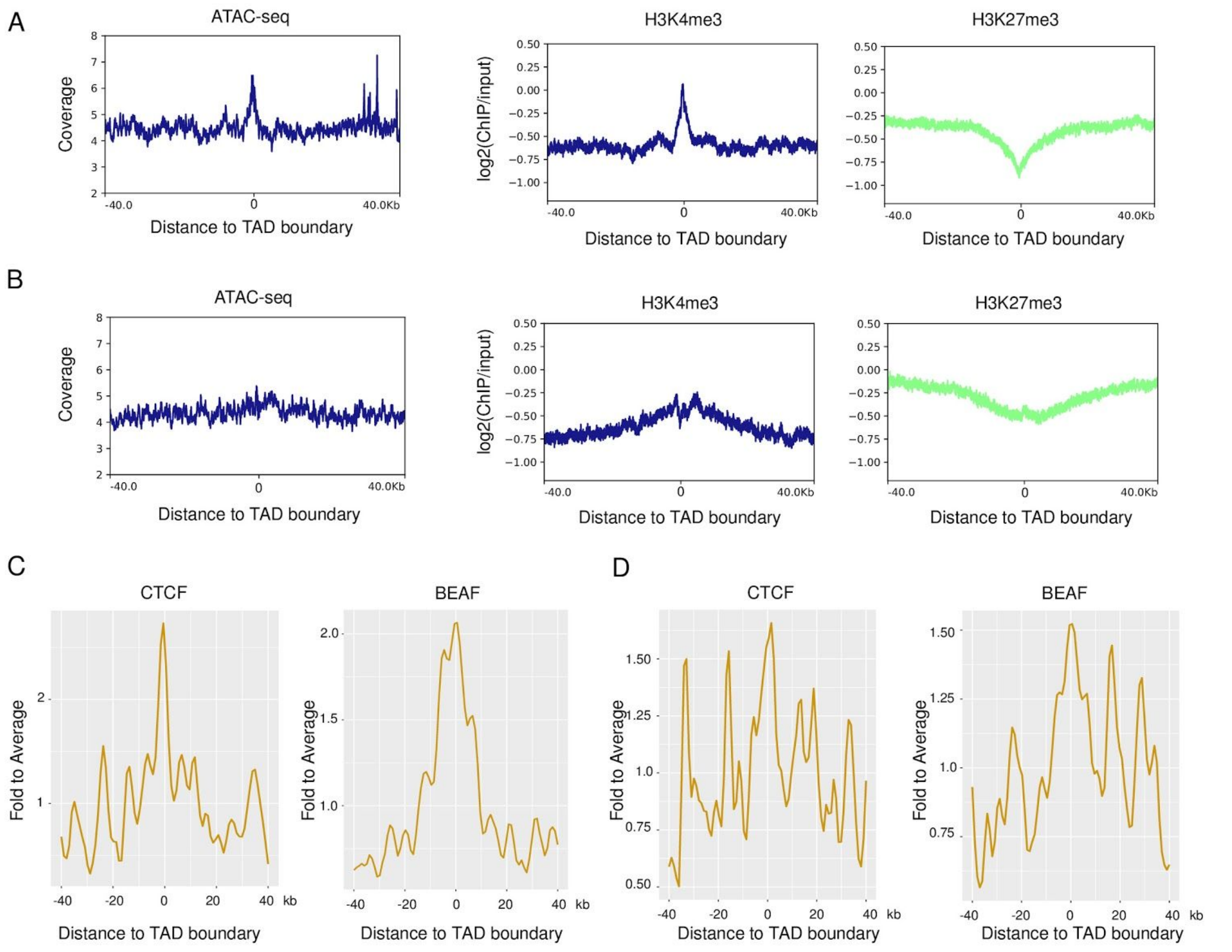


**Supplemental Figure S4:** Enrichment analysis of insulator binding sites (BEAF-32 and CTCF), epigenetic marks (H3K4me3 and H3K27me3) and ATAC-seq at TAD boundaries. TAD boundaries in (*A*) and (*C*) were identified using Armatus, TAD boundaries in (*B*) and (*D*) were identified using Arrowhead from the Juicer package (Supplemental to Figure 1).

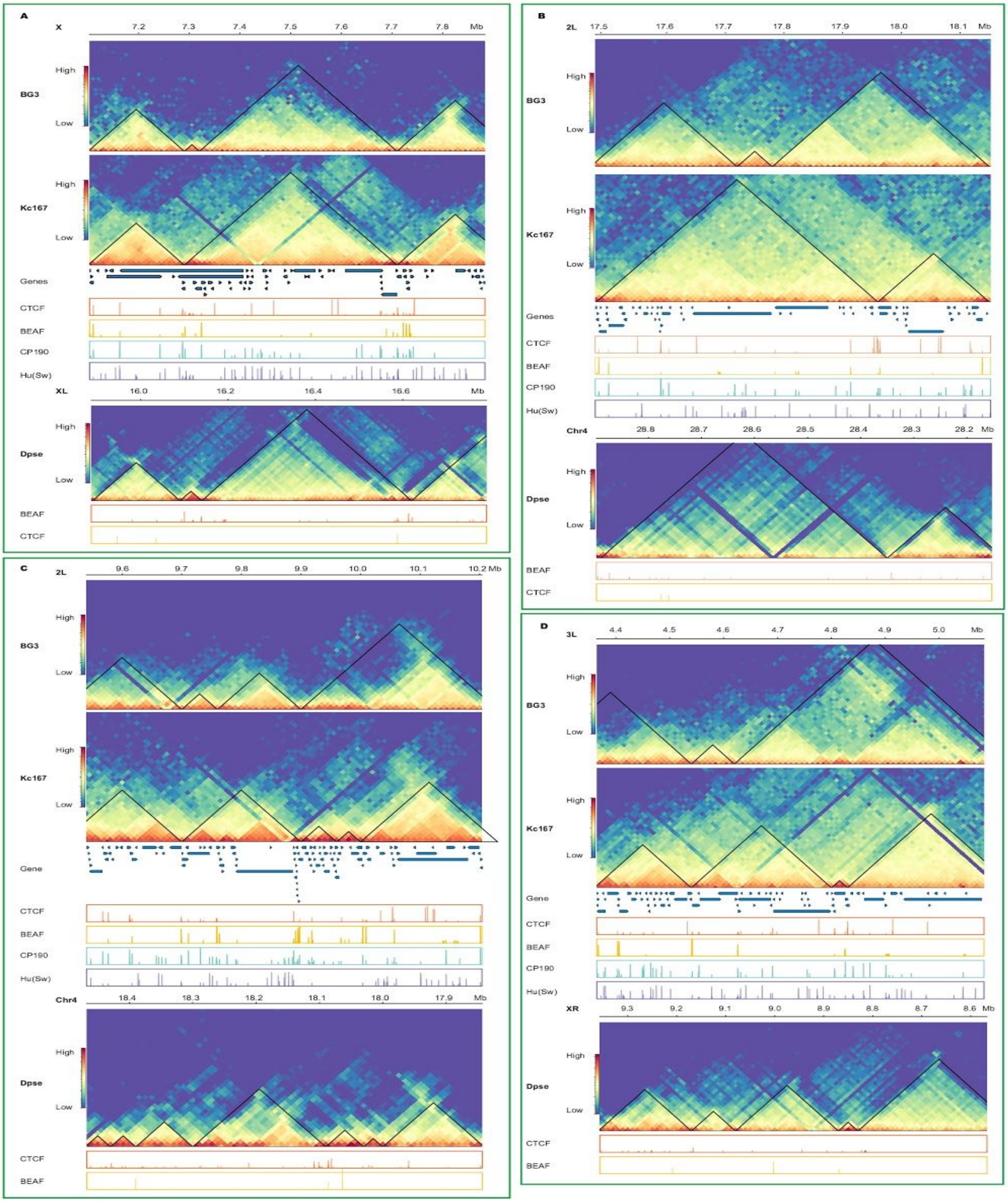


Continued in the next page

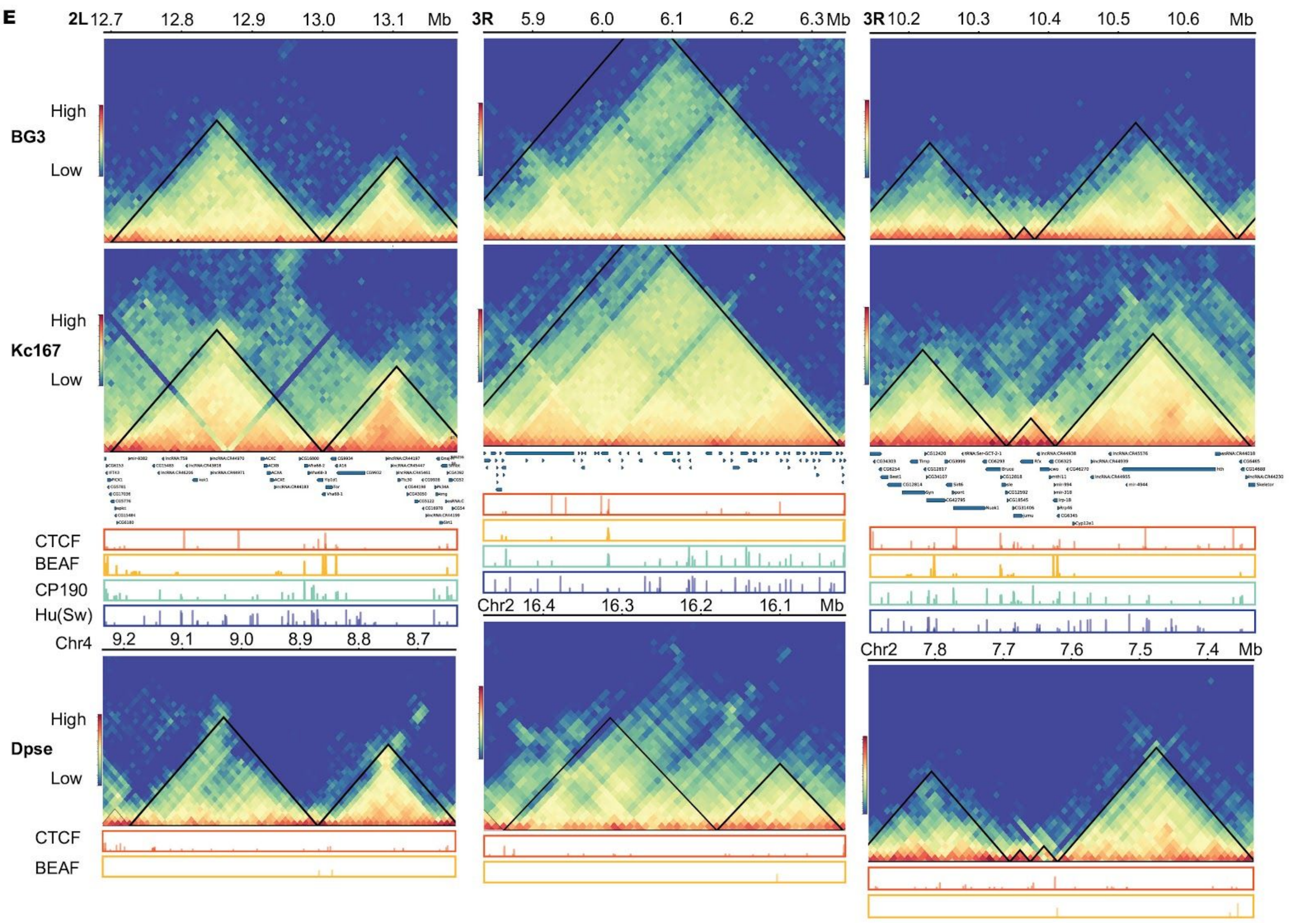


**Supplemental Figure S5**: Conservation of TADs at the syntenic regions between *D. melanogaster* and *D. pseudoobscura*.

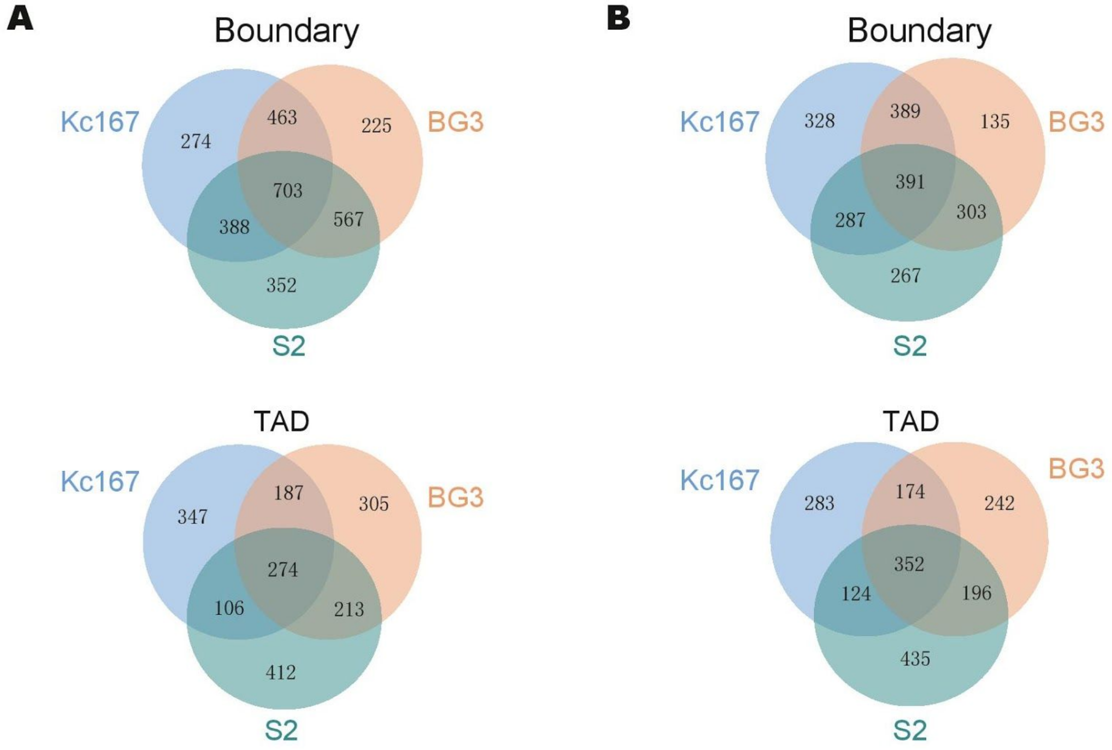


**Supplemental Figure S6:** TADs are largely shared across cell lines in *D. melanogaster*. (*A*) TADs from three different cell lines (Kc167, BG3, and S2) identified using Armatus. (*B*) TADs identified using Arrowhead.

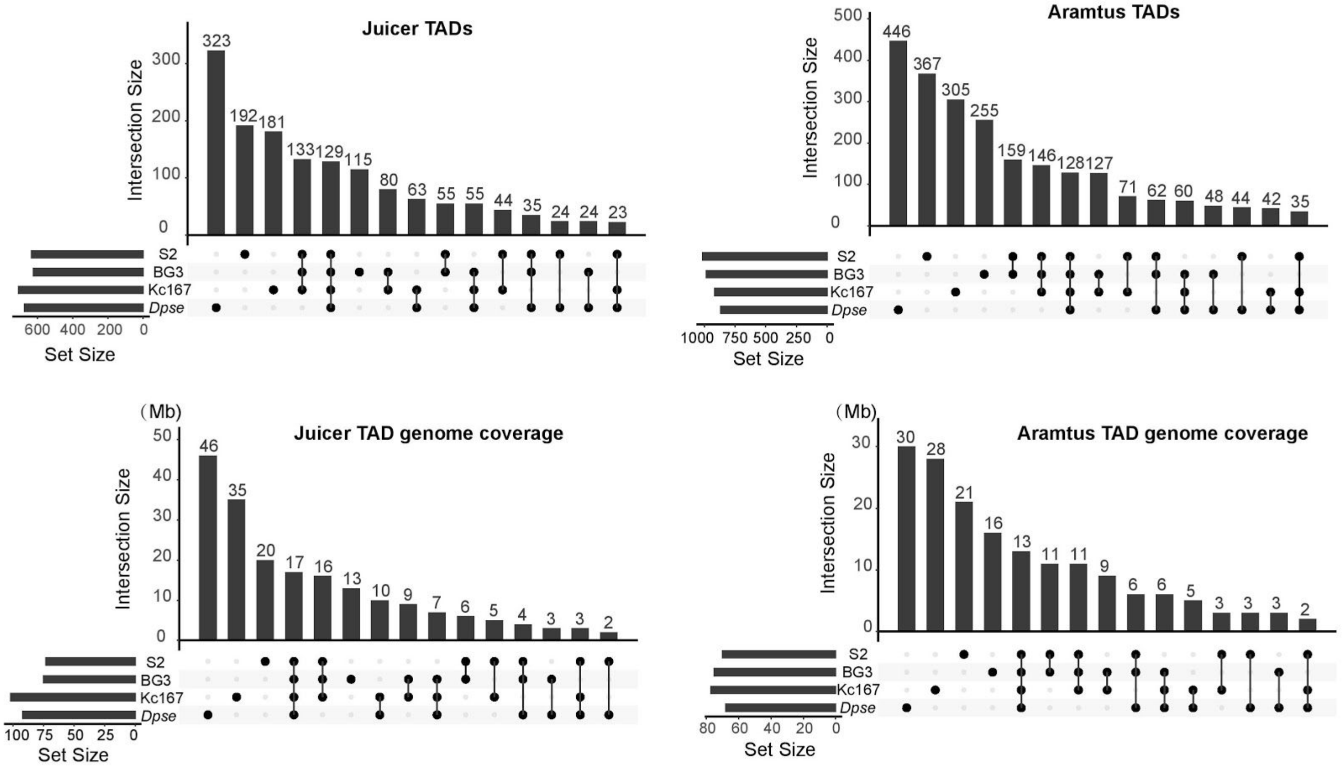


**Supplemental Figure S7:** Upset plot showing the overlap of TAD structures among three cell lines in *D. melanogaster,* as well as *D. pseudoobscura* whole body (Supplemental to Figure 2). (Left) TADs were identified using Arrowhead from the Juicer package; (Right) TADs were identified using Armatus.

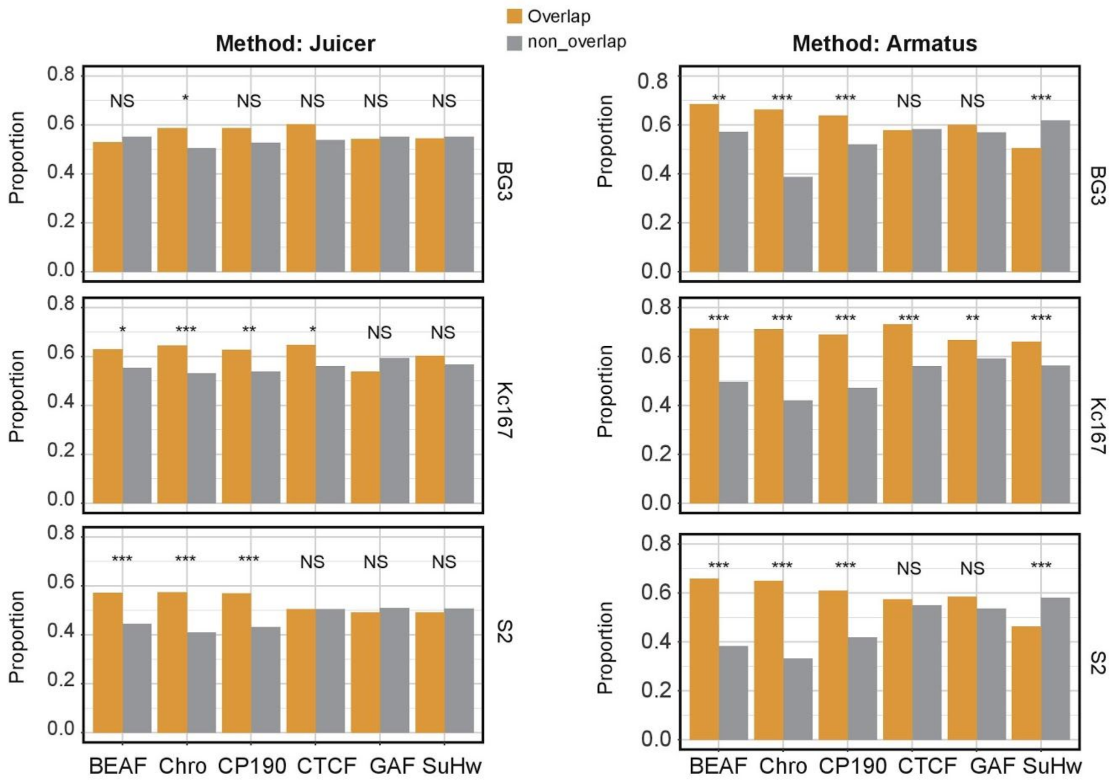


**Supplemental Figure S8**: Conservation of TAD boundaries that overlapped with different insulator binding sites including BEAF-32, Chro, CP190, CTCF, GAF (Trl) and Su(Hw) between *D. melanogaster* (Kc167, BG3, and S2) and *D. pseudoobscura* (full body) (Supplemental to Figure 3). (Left), TAD boundaries were identified using Arrowhead from the Juicer package; (Right), TAD boundaries were identified using Armatus.

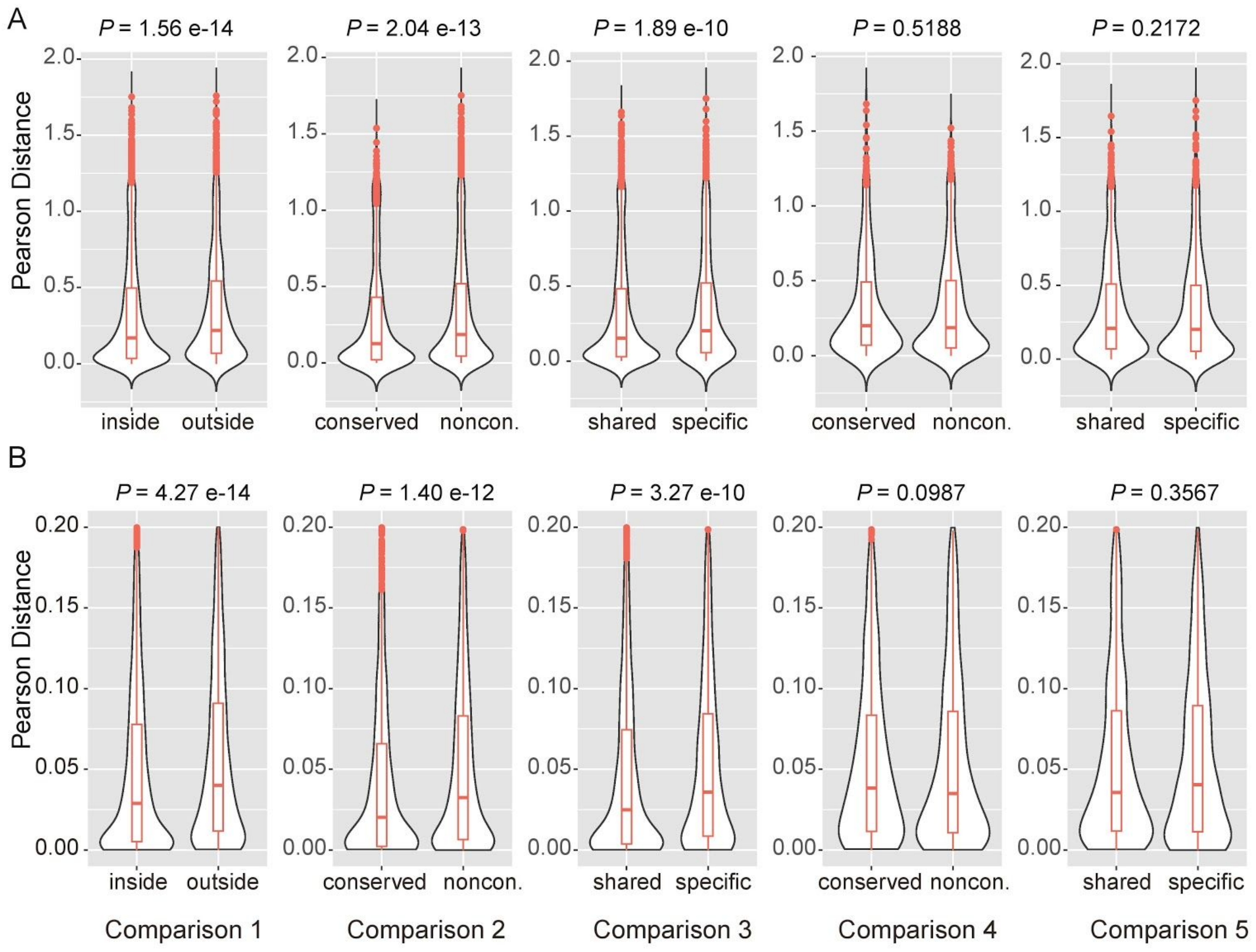


**Supplemental Figure S9:** The potential roles of TADs in gene regulation in *Drosophila* (Supplemental to Figure 4). (*A*) Expression divergence measured by Pearson's correlation coefficient distance for one-to-one orthologs between *D. melanogaster* and *D. pseudoobscura*. (*B*) Expression variation measured by Pearson's correlation coefficient distance for the same gene sets used in the above interspecific comparison between two *D. melanogaster* strains, OreR and w1118.

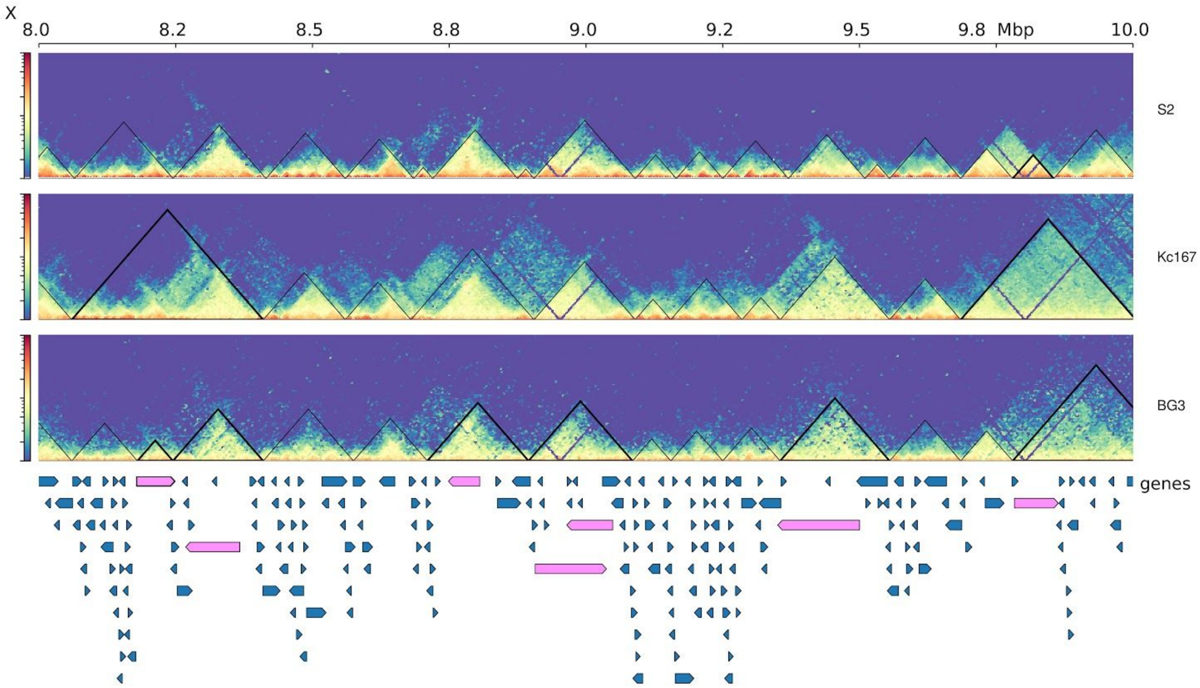


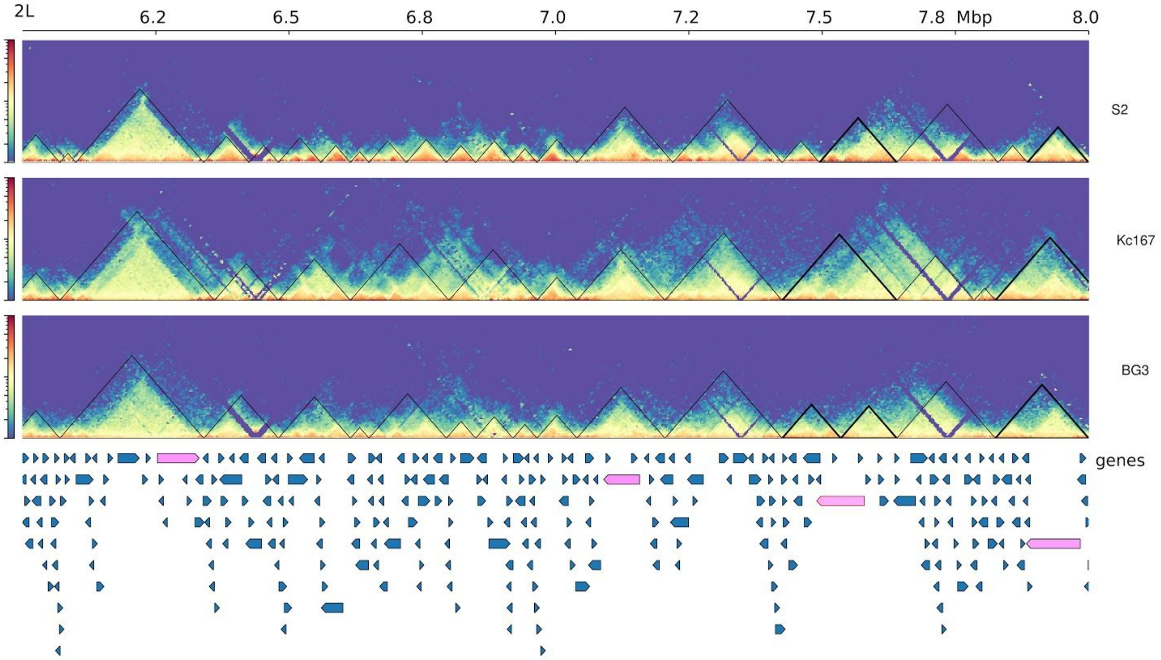


**Supplemental Figure S10:** Long coding genes (>50 kbp) tend to span full TADs observed in *D. melanogaster*. Top panels represent TADs that were annotated in three *D. melanogaster* cell lines. Below represent gene structures with long genes highlighted in pink color.

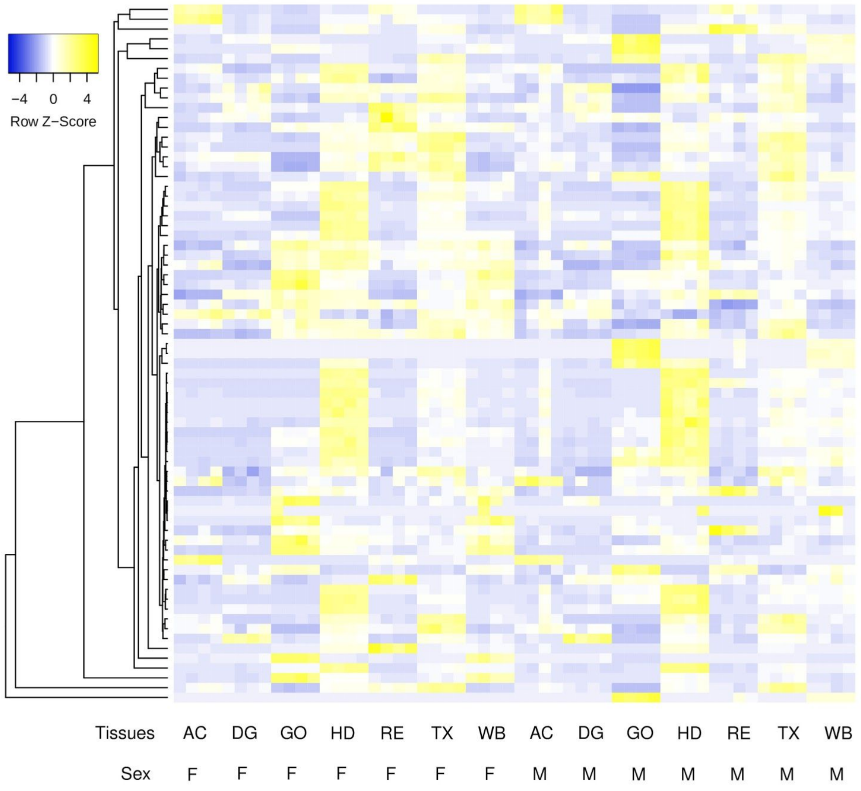


**Supplemental Figure S11:** Expression profile of 71 long genes (> 50 kbp) that overlapped with complete TADs across 7 tissues from both sex in *D. pseudoobscura* (Supplemental to Figure 4). Long genes are preferentially expressed in the head. AC, abdomen without digestive or reproductive system; DG, digestive plus excretory system; GO, gonad; HD, head; RE, reproductive system without gonad; TX, thorax without digestive system; WB, whole body. F, female; M, male.

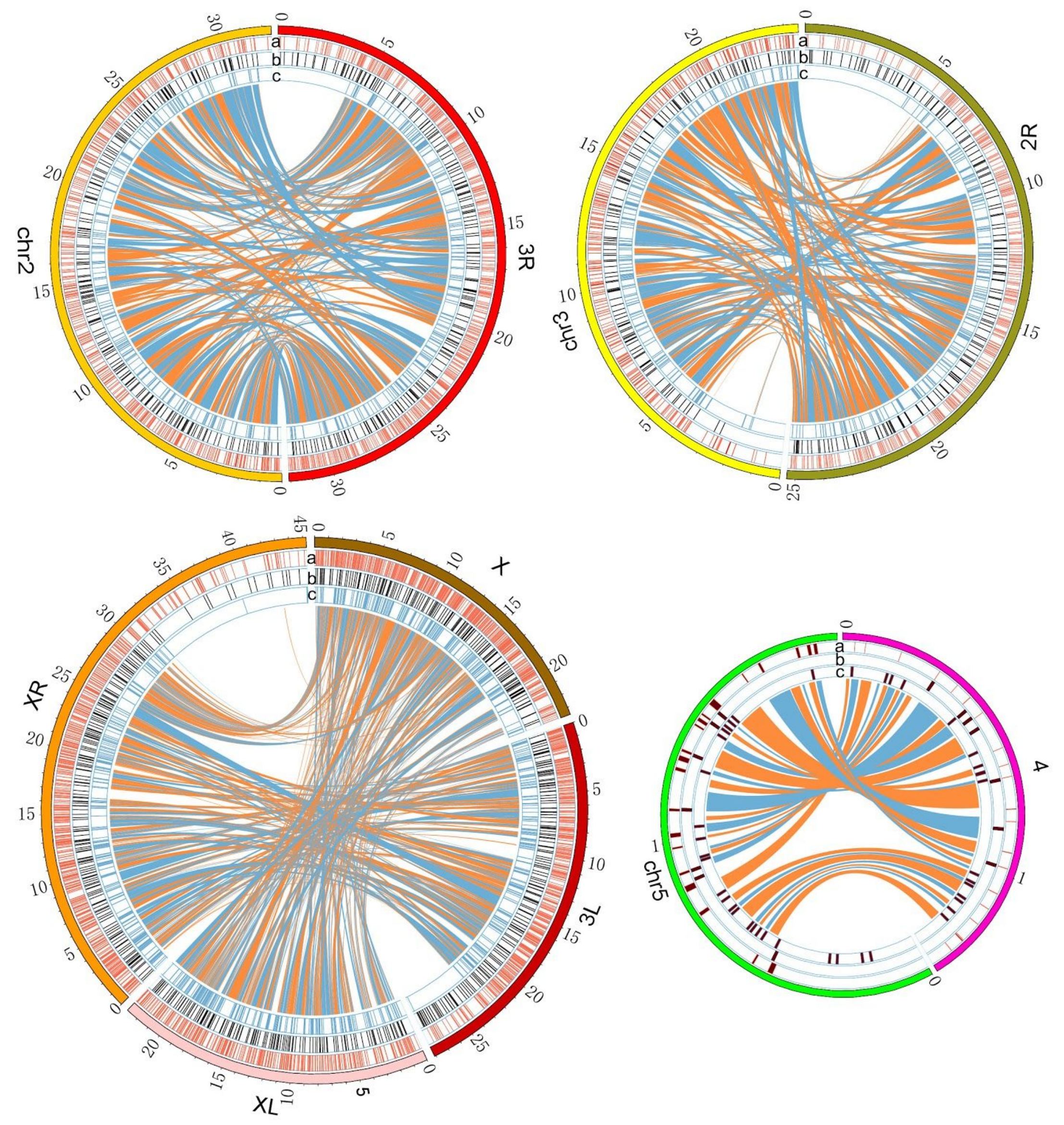


**Supplemental Figure S12:** Synteny map between *D. melanogaster* and *D. pseudoobscura* (Supplemental to Figure 5A). Tracks a: TAD boundaries annotated by HiCExperor at restriction fragment resolution; b: 10 kbp resolution; c: Synteny breakpoints.

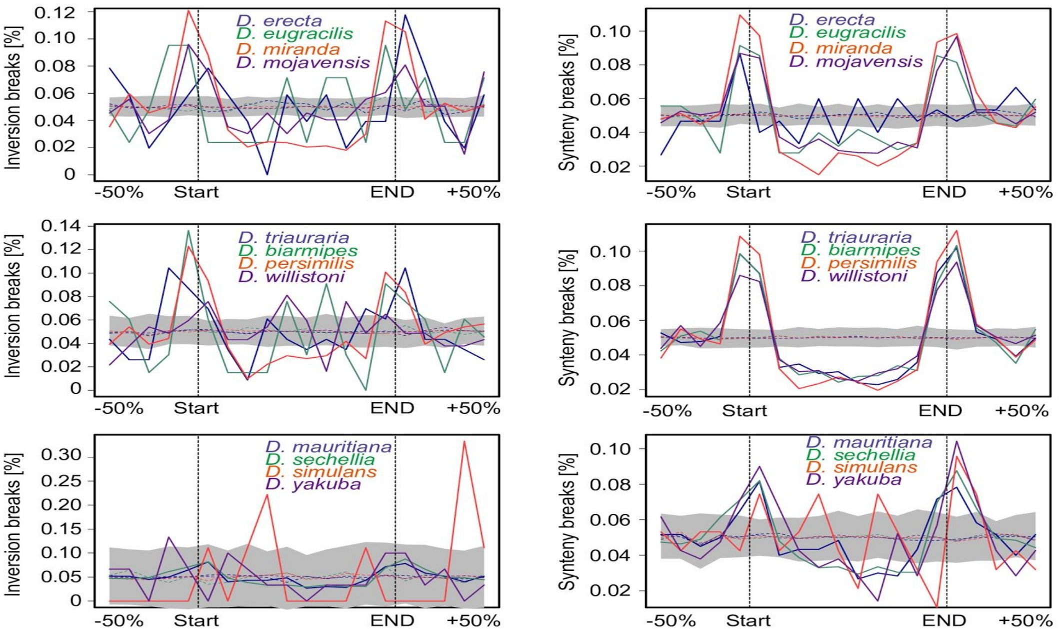


***D. melanogaster* TAD**

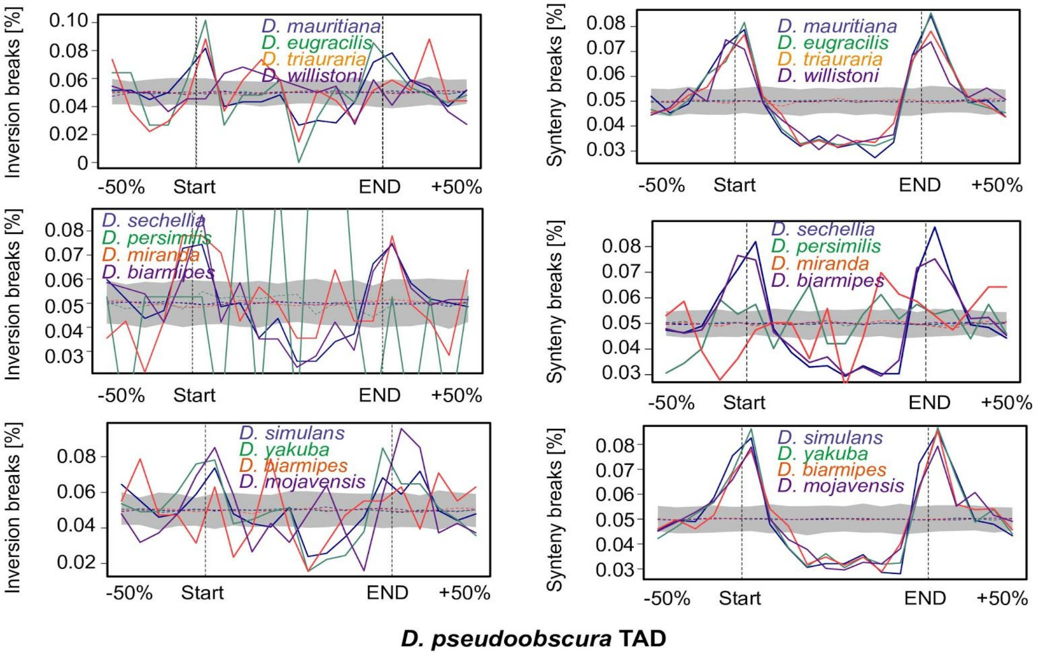


***D. pseudoobscura* TAD**

**Supplemental Figure S13**: Distribution of evolutionary synteny/inversion breakpoints around TAD boundaries (Supplemental to Figure 5*C*,*D*). (Top) Distribution of genome rearrangement breakpoints between *D. melanogaster* and twelve other *Drosophila* species along TAD regions. (Bottom) Comparisons in *D. pseudoobscura*.

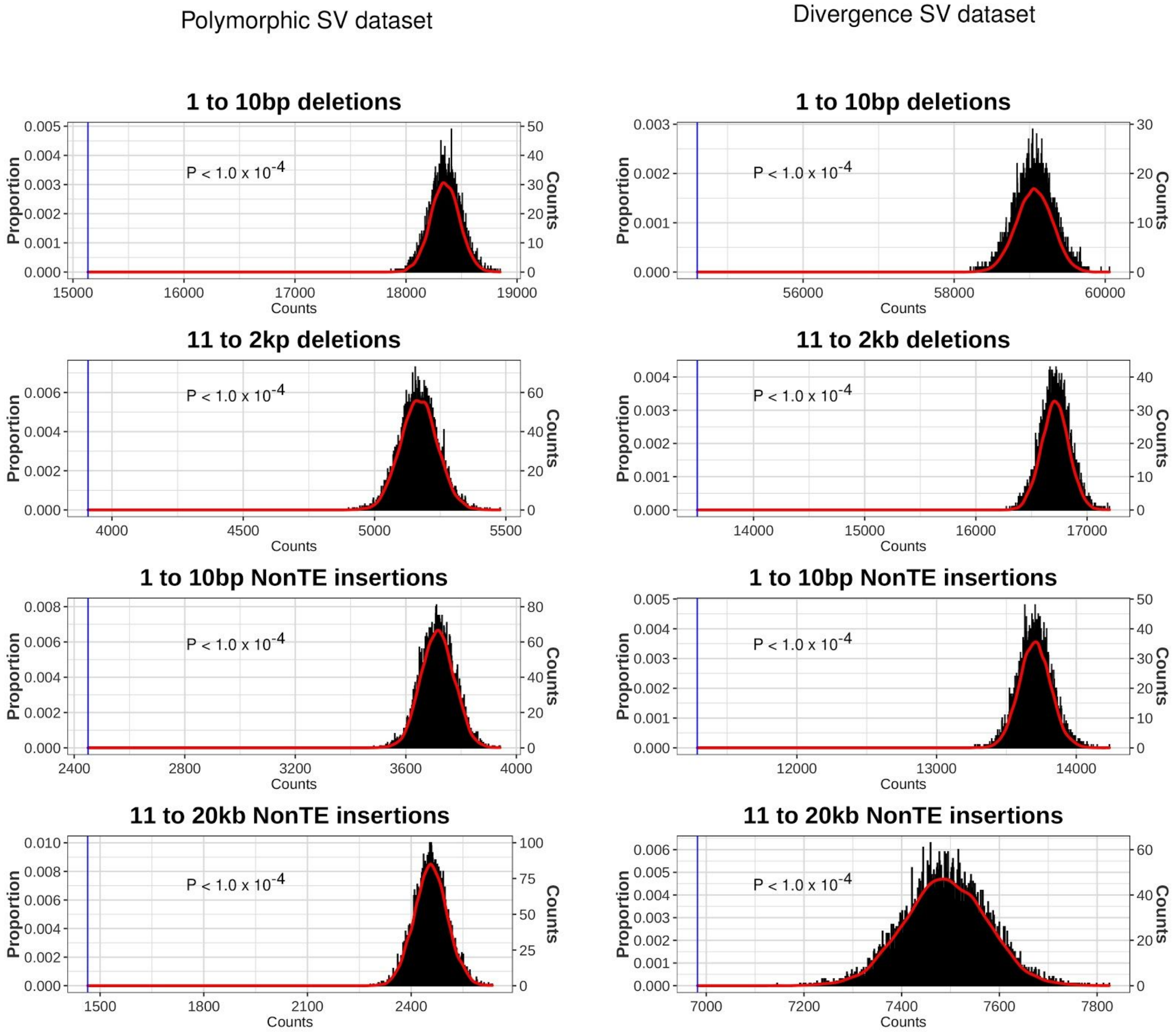


**Supplemental Figure S14:** Permutation tests (n= 10,000) for evaluating if deletions and non-TE insertions are significantly enriched or depleted at TAD boundaries. Left shows results for polymorphic dataset with genomic variants genotyped in 14 *D. melanogaster* species. Right shows results for divergence dataset with genomic variants genotyped in three *D. simulans* clade species (*D. mauritania*, *D. simulans* and *D. sechellia*). We simulated 10,000 samples of TAD boundaries coordinates with size and chromosomal distribution confined by the actual dataset. *P*-values were calculated based on the observation against the random distribution of frequency that SVs that overlapped TAD boundaries for each type of mutation. The blue lines represent the observed value in our real datasets.

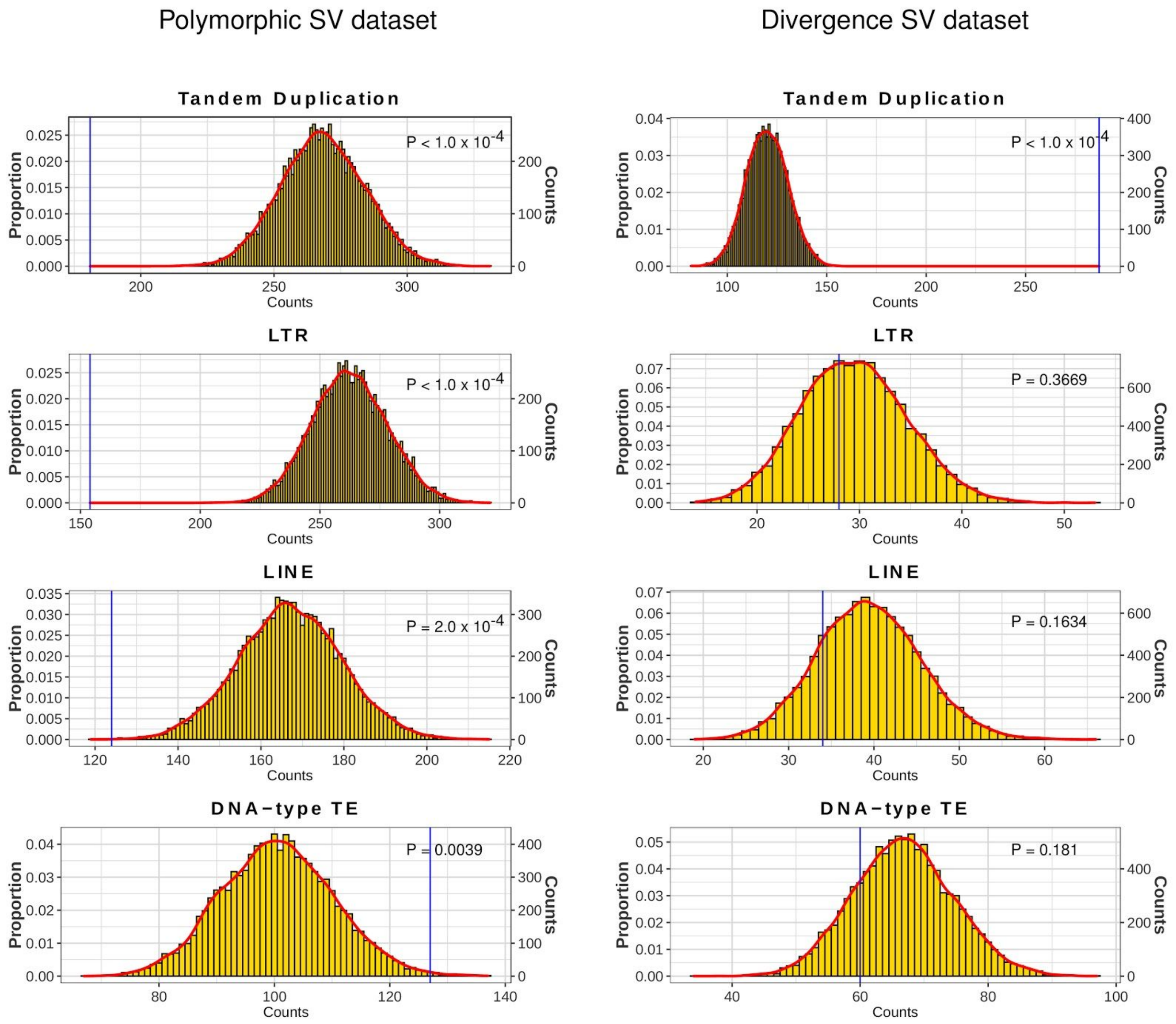


**Supplemental Figure S15:** Permutation tests (n= 10,000) for evaluating if tandem duplications and three classes of TE insertions (LTRs, LINEs and DNA-type elements) are significantly enriched or depleted at TAD boundaries. Left shows results for polymorphic dataset with genomic variants genotyped in 14 *D. melanogaster* species. Right shows results for divergence dataset with genomic variants genotyped in three *D. simulans* clade species (*D. mauritania*, *D. simulans* and *D. sechellia*). We simulated 10,000 samples of TAD boundaries coordinates with size and chromosomal distribution confined by the actual dataset. *P*-values were calculated based on the observation against the random distribution of frequency that SVs that overlapped TAD boundaries for each type of SVs. The blue lines represent the observed value in our real datasets.

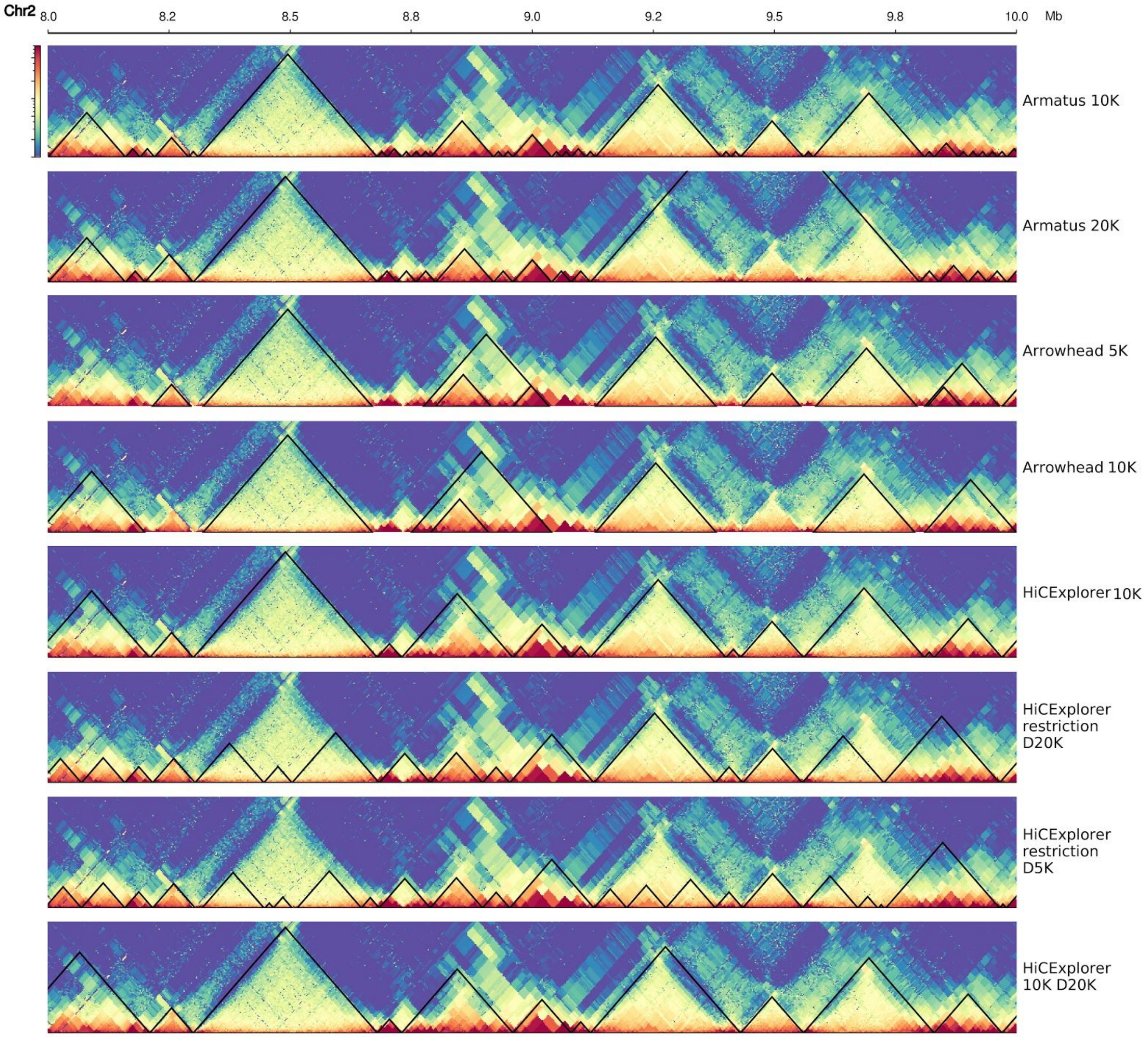


**Supplemental Figure S16:** Comparison of TAD annotation result for Armatus, Arrowhead in the Juicer package and HiCExplorer under different Hi-C contact map resolutions (restriction fragment resolution, 5 kbp, 10 kbp or 20 kbp). Genomic region on *D. pseudoobscura* chromosome 2 from 8,000,000 to 10, 000,000 (bp) was shown as an example.
